# Small Molecules Restore Azole Activity Against Drug-Tolerant and Drug-Resistant *Candida* Isolates

**DOI:** 10.1101/2022.03.31.486631

**Authors:** Philip E. Alabi, Cécile Gautier, Thomas P. Murphy, Xilin Gu, Mathieu Lepas, Vishukumar Aimanianda, Jason K. Sello, Iuliana V. Ene

**Author notes:** equal contributions (Philip E. Alabi, Cécile Gautier).

## Abstract

Each year, fungi cause more than 1.5 billion infections worldwide and have a devastating impact on human health, particularly in immunocompromised individuals or patients in intensive care units. The limited antifungal arsenal and emerging multidrug resistant species necessitate the development of new therapies. One strategy for combating drug resistant pathogens is the administration of molecules that restore fungal susceptibility to approved drugs. Accordingly, we carried out a screen to identify small molecules that could restore the susceptibility of pathogenic *Candida* species to azole antifungals. This screening effort led to the discovery of novel 1,4-benzodiazepines that restore fluconazole susceptibility in resistant isolates of *Candida albicans*, as evidenced by 100-1000-fold potentiation of fluconazole activity. This potentiation effect was also observed in azole-tolerant strains of *C. albicans* and in other pathogenic *Candida* species. The 1,4-benzodiazepines selectively potentiated different azoles, but not other approved antifungals. A remarkable feature of the potentiation was that the combination of the compounds with fluconazole was fungicidal, whereas fluconazole alone is fungistatic. Interestingly, the potentiators were not toxic to *C. albicans* in the absence of fluconazole, but inhibited virulence-associated filamentation of the fungus. We found that the combination of the potentiators and fluconazole significantly enhanced host survival in a *Galleria mellonella* model of systemic fungal infection. Taken together, these observations validate a strategy wherein small molecules can restore the activity of highly used anti-infectives that have lost potency.

**IMPORTANCE:** In the last decade, we have been witnessing a higher incidence of fungal infections, due to an expansion of the fungal species capable of causing disease (*e.g*., *Candida auris*), as well as increased antifungal drug resistance. Among human fungal pathogens, *Candida* species are a leading cause of invasive infections and are associated with high mortality rates. Infections by these pathogens are commonly treated with azole antifungals, yet the expansion of drug-resistant isolates have reduced their clinical utility. In this work, we describe the discovery and characterization of small molecules that potentiate fluconazole and restore the susceptibility of azole-resistant and azole-tolerant *Candida* isolates. Interestingly, the potentiating 1,4-benzodiazepines were not toxic to fungal cells but inhibited their virulence-associated filamentous growth. Furthermore, combinations of the potentiators and fluconazole decreased fungal burdens and enhanced host survival in a *Galleria mellonella* model of systemic fungal infections. Accordingly, we propose the use of novel antifungal potentiators as a powerful strategy for addressing the growing resistance of fungi to clinically approved drugs.

## INTRODUCTION

Underlying more than 1.5 billion infections worldwide and an estimated 1.6 million deaths annually, fungal infections pose a significant threat to human health (1, 2). While most fungal infections are superficial, some can cause life-threatening invasive infections, especially in immunocompromised individuals (1, 3). Among pathogenic fungi, *Candida* species are especially concerning because they are the leading cause of systemic fungal infections with mortality rates reaching 50% (4, 5). The incidence of life-threatening *Candida* infections is on the rise due to a growing proportion of the population with weakened immune systems and increasing numbers of isolates that are tolerant and/or resistant to antifungals (2, 6–9). Most *Candida* infections are caused by *C. albicans* and as much as 7% of isolates of this species are reportedly resistant to drugs (10, 11). Growing incidences of therapeutic failure in *C. albicans* and increasing observations of drug resistant strains of this species and others like *Candida auris* and *Candida glabrata* are an impending public health crisis (9, 10, 12).

The emergence and expansion of drug-resistant *Candida* species are gradually eroding the utility of the current repertoire of antifungals (*e.g.*, azoles, polyenes, echinocandins) used to treat invasive infections (10, 13). New antifungals are desperately needed, but drug discovery efforts are complicated by the phylogenetic conservation of druggable proteins and pathways in fungi and humans. Indeed, the small number of clinically used antifungals having good safety profiles critically perturb phenomena that are unique to fungi. A class of drugs having desirable selectivity and safety profiles are the azoles and the prototypical member of this class of drugs is fluconazole (FLC) (8, 13–18). Azoles are fungistatic agents that act by inhibiting cytochrome P-450DM which catalyzes the conversion of lanosterol to ergosterol, an essential component of the fungal plasma membrane (19). Unfortunately, the azoles are also illustrative examples of how pathogenic fungi can become tolerant and resistant to antifungals. Azole resistance is commonly associated with mutations in the drug target, over-expression of genes encoding efflux pumps, and/or increased copy numbers of beneficial alleles (13, 16, 17, 20). Tolerance to azoles is ascribed to intrinsic heterogeneity in fungal populations and is mediated by multiple stress response pathways (8, 14). About half of *Candida* isolates display tolerance to azoles, and this phenomenon is increasingly associated with persistent candidiasis (8, 14, 21–24), making it critically important for future antifungal drug development.

The small number of drug targets that are unique to fungi and the intrinsic fungal tolerance and resistance to existing drugs make the development of new antifungals a challenging endeavor. An alternative to developing novel antifungals is identifying compounds that can restore drug susceptibility in pathogenic fungi. Indeed, there are several reports that describe molecules that potentiate azole activity against drug-resistant fungi. For instance, a chemical screen using the *Cryptococcus neoformans* overlap^2^ method (O2M) led to the identification of dicyclomine, sertraline, proadifen and chloroquine as azole potentiators (25). Similarly, inhibitors of Hsp90 and calcineurin were reported to suppress resistance and tolerance to FLC and combinations of the inhibitors with FLC were fungicidal (14, 26–31). In another study, potentiation of FLC activity against *C. albicans* and *C. glabrata* by inhibitors of vesicular trafficking was described (32).

Inspired by reports of potentiators of azole activity, we performed a screen of an in-house collection of compounds in search of molecules that could restore FLC susceptibility to resistant and tolerant isolates of *C. albicans*. Here, we report the discovery and characterization of small molecules that restore azole activity against resistant and tolerant strains of *C. albicans*. Interestingly, the azole potentiators are novel 1,4-benzodiazepines (1,4-BZDs) having markedly different structures and activities from previously reported antifungal benzodiazepines. The compounds have no intrinsic antifungal activity, yet their azole potentiation was synergistic and led to 100-1000-fold enhancement of FLC activity, which was higher than that of previously reported potentiators (25, 32). The potentiation was accompanied by a change in azole antifungal activity. Specifically, while FLC alone is fungistatic, several combinations of FLC with potentiators were fungicidal. Another notable feature of the potentiators is that they do not have intrinsic antifungal activity, yet they inhibit filamentation in *C. albicans.* Although the mechanism by which the compounds act is not yet clear, their capacity to potentiate several inhibitors of ergosterol synthesis and an inhibitor of sphingolipid biosynthesis implies that the 1,4-BZDs may act by perturbing lipid homeostasis. Importantly, combinations of FLC and potentiators significantly decreased fungal burdens and enhanced host survival in a *G. mellonella* model of fungal infection.

## RESULTS

### Novel 1,4-BZDs are potentiators of FLC activity

To identify compounds that potentiate FLC activity against drug-resistant and tolerant *C. albicans* strains, we screened an unbiased and structurally diverse library of ∼100 compounds that had been synthesized in our laboratories. The screen included sortin2, dicyclomine, sertraline, proadifen and chloroquine as positive controls, owing to their previously reported azole potentiating activities (Fig. 1A) (25, 32). Careful consideration was also given to the *C. albicans* strains tested in the screen. We selected isolates with different FLC resistance/tolerance levels and from diverse genetic backgrounds; specifically, the FLC-susceptible (yet tolerant) strain SC5314 [Clade 1, (33)], and the FLC-resistant strains SP-945 [Clade 3, (16, 34)] and P60002 [Clade SA, (33)] (Fig. 2A).

**Figure 1.**
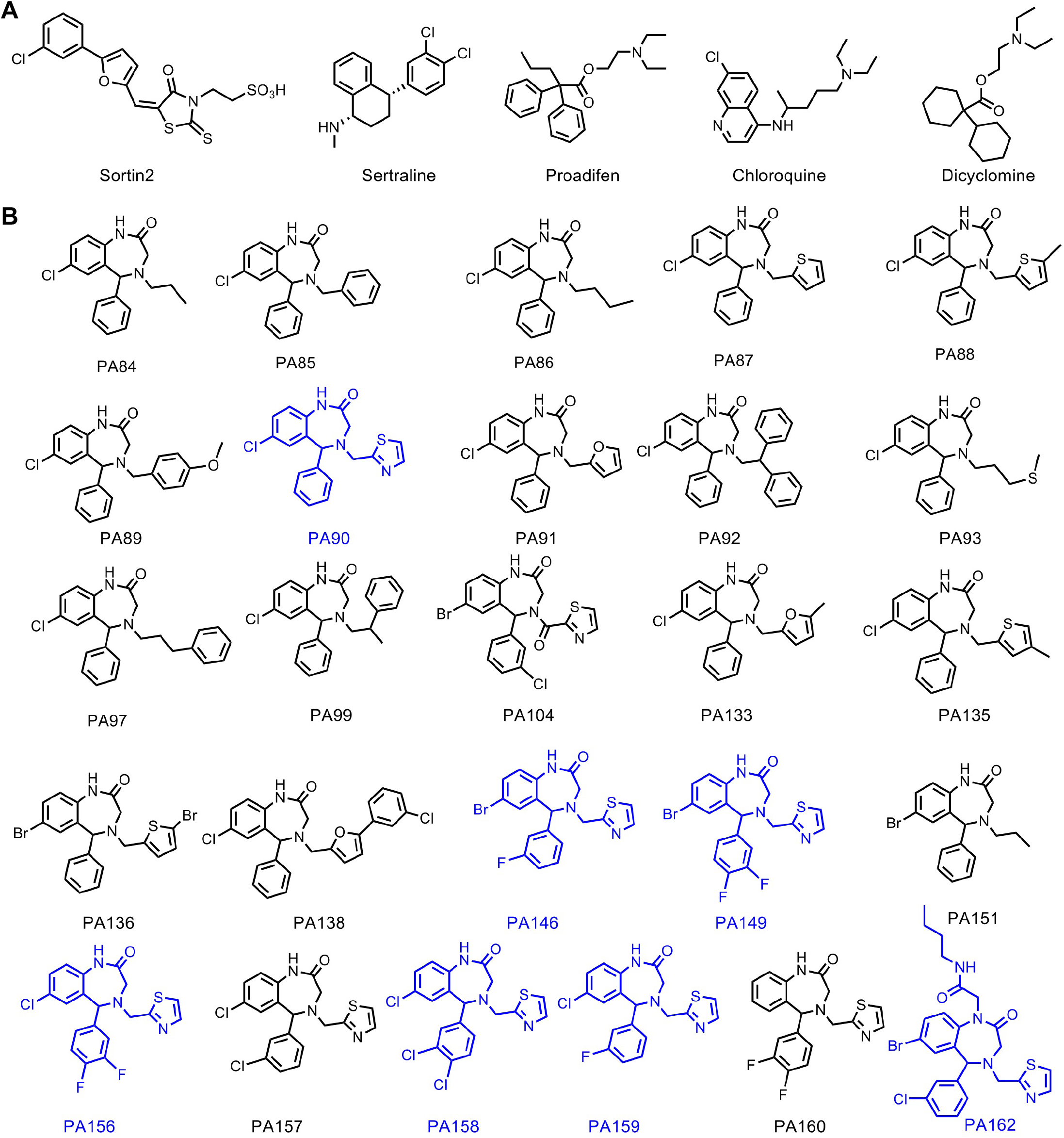
Chemical structures of control compounds (A) and the set of twenty-six 1,4-BZDs (B) tested. The most active 1,4-BZDs are indicated in blue.

**Figure 2.**
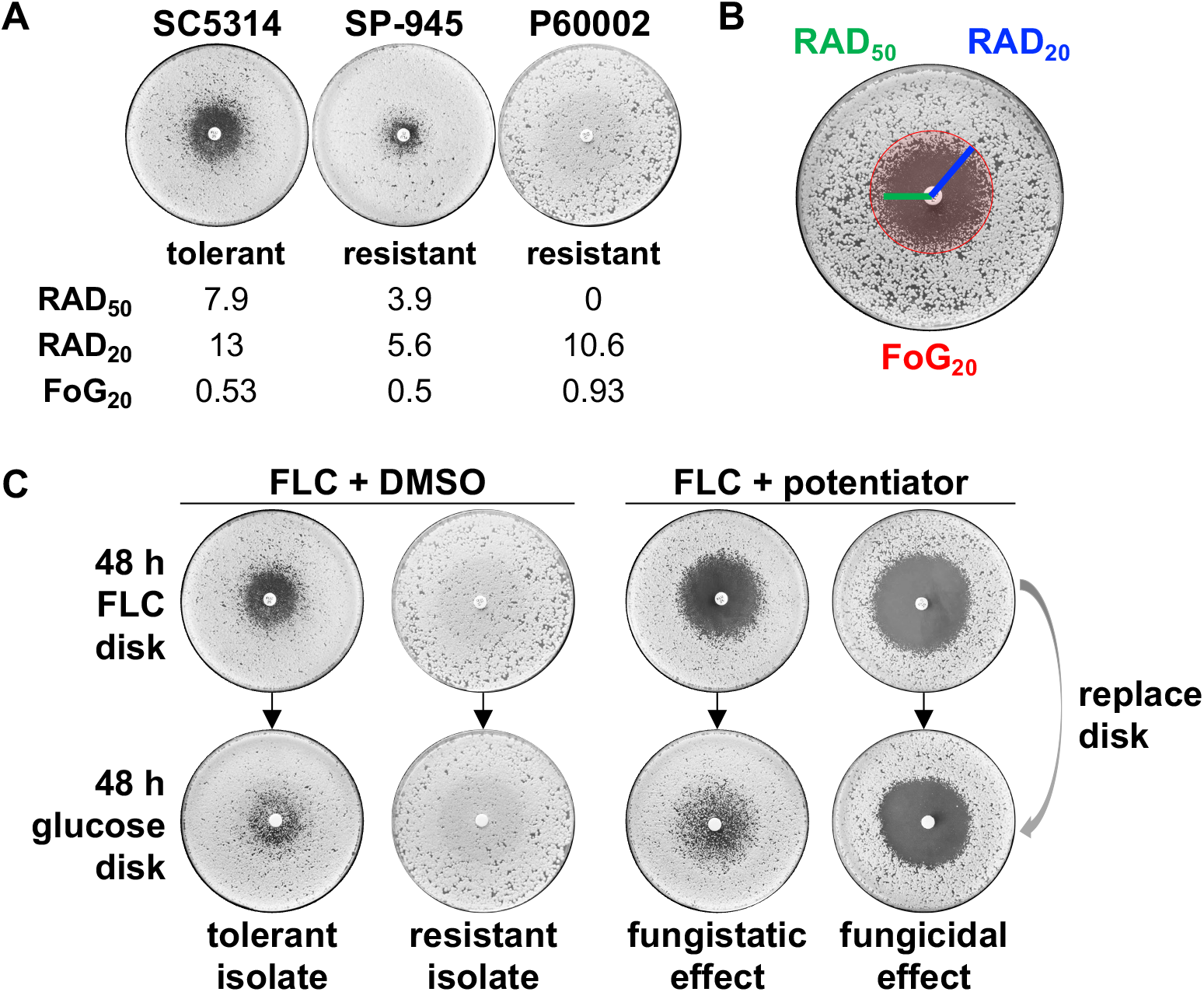
Screening strategy for 1,4-BZDs and control compounds. (A) FLC susceptibility profiles of *C. albicans* isolates SC5314, SP-945, and P60002 on disk diffusion assays, including their mean RAD50, RAD20 and FoG20 levels. (B) A lawn of *C. albicans* cells was plated on YPD agar and grown in the presence of a drug disk (FLC, 25 µg) for 48 h. The size of the radius at 50% and 20% growth inhibition (RAD50 and RAD20, respectively) was measured as a proxy for drug susceptibility. The fraction of growth at 20% inhibition (FoG20) was measured as a proxy for tolerance. Susceptibility and tolerance were determined using *diskImageR*. (C) Following 48 h of growth, the FLC disk was replaced with a glucose disk (G, 8 mg) and the plates were grown for an additional 48 h. For drug combinations, YPD plates contained 1,4-BZDs at 100 µM. Plates show example profiles of tolerant and resistant strains with FLC alone (left, with DMSO used as control). Plates with FLC and potentiators (right) illustrate fungistatic or fungicidal effects of different drug combinations following additional growth on glucose disks.

The screen leveraged a disk diffusion assay in which changes in both drug susceptibility and tolerance could be assessed (35). We purposefully examined both phenomena because a singular focus on susceptibility alone would not account for fungal growth and adaptation to the drug combination over an extended assay time (14). The assays were performed over the course of 48 hours with a FLC disk placed on solid media supplemented with members of the library at 100 μM. Susceptibility was quantified by the clearance of fungal growth around the FLC disk at 50% and 20% growth inhibition (RAD_50_ and RAD_20_, respectively, Fig. 2B) (35). On the other hand, tolerance was assessed by the fraction of growth inside the area of drug inhibition at 20% growth inhibition after 48 h (FoG_20_, Fig. 2B). In combination with FLC, potentiators were expected to effect larger inhibition zones and/or decreased FoG_20_ levels relative to those observed when FLC was administered alone (Fig. 2C). In parallel, it was possible to determine whether the combination of FLC and screened compounds had fungistatic or fungicidal effects by replacing the FLC disk with a glucose disk and by measuring RAD_50_, RAD_20_ and FoG_20_ following an additional 48 h growth period (Fig. 2C).

We were intrigued that several molecules that potentiated FLC activity against all three *C. albicans* strains tested were 1,4-BZDs. Benzodiazepines are privileged scaffolds in pharmacology and drug discovery having features that favor protein binding, including a semi-rigid scaffold, intermediate lipophilicity, and a combination of hydrogen bond donors and acceptors. This scaffold is found in several FDA approved drugs (36–38) and continues to be exploited in drug discovery efforts (39–45). The seven hits in the screen were among a group of twenty-six unique 1,4-BZDs in the library that differed by substitution patterns and the presence of either secondary or tertiary amide moieties (Fig. 1B). Each of the compounds was the product of a diversity-oriented synthetic route. The first building blocks in the synthetic scheme were substituted diaryl ketoacetamides that were derived from substituted phenylacetamides (1a, Fig. S1) in a palladium mediated C-H activation. Acid-promoted N-deacetylation of those intermediates yielded diaryl aminoketones (2a) (46) that were elaborated into 1,4-BZDs having either secondary (3a) or tertiary amide moieties (4a). The former (3a) were accessed via a sequence of amidation followed by cyclization via reductive amination; the latter (4a) were prepared via a tandem Ugi-four component reaction and cyclization via reductive amination. In either case, the advanced intermediates were elaborated into the aforementioned twenty-six 1,4-BZDs via acylations or reductive aminations (5a, 6a, 7a, Fig. S1).

From the perspective of assessing structure activity-relationships, it was fortuitous that the seven potentiators were among twenty-six 1,4-BZDs in our compound collection. In fact, we noted that most of the potentiators were 1,4-BZDs having secondary amides (PA90, PA146, PA149, PA156, PA158, PA159); whereas only a single screening hit had a tertiary amide (PA162). Aside from the secondary and tertiary amide distinction, there were other clear structural correlates for potentiation activity. For example, the potentiating compounds had large halogen substituents at the 7-position of the fused benzo moiety of the 1,4-BZD core. More specifically, compounds having either bromine or chlorine substituents were active, whereas those having a fluorine or a hydrogen atom at the 7-position were less active. It was also interesting that the potentiating compounds, except for PA90, had halogen substituents on the exocyclic phenyl moiety. Compounds having fluorine or chlorine substituents at the 3- and/or 4-position of the exocyclic phenyl moiety were superior potentiators. Finally, all the potentiating compounds had a methylene 2-thiazole at N-4 of the 1,4-BZD core structure. This was critical as compounds having aliphatic or other aromatic substituents in the same N-4 position did not potentiate FLC activity. Taken together, these observations indicate that a class of 1,4-BZDs having 7-Br/7-Cl substituted benzo groups, 3 and/or 4 halogen substituted exocyclic phenyl and N-methylenethiazyl group can significantly potentiate the activity of FLC against azole-resistant *C. albicans* strains.

Interestingly, the seven screening hits (PA90, PA146, PA149, PA156, PA158, PA159 and PA162) restored FLC susceptibility and suppressed FLC tolerance in both resistant and tolerant *Candida* strains. The influence of the seven 1,4-BZDs over azole susceptibility was evidenced by RAD_50_ values that increased by 2.2 to 387-fold and by 1.4 to 2.4-fold increases in RAD_20_ values across the three *C. albicans* strains (Fig. 3A and S2A). The compounds also showed significant effects on FLC tolerance, with FoG_20_ levels decreasing 2.9 to 4.4-fold across the three strains; notably, most of the compounds efficiently suppressed tolerance (Fig. 3B and S2B). The capacity of the seven 1,4-BZDs to potentiate FLC against *C. albicans* strains with different resistance and tolerance levels was striking because previously reported FLC potentiators had strain specific effects. For example, proadifen only potentiated FLC against P60002 and sertraline potentiated FLC against SP-945, while showing reduced activity against other isolates (Fig. S2A-B). Additionally, the 1,4-BZDs exhibited greater potency than the previously reported potentiators in our experiments (Fig. S2A-B).

**Figure 3.**
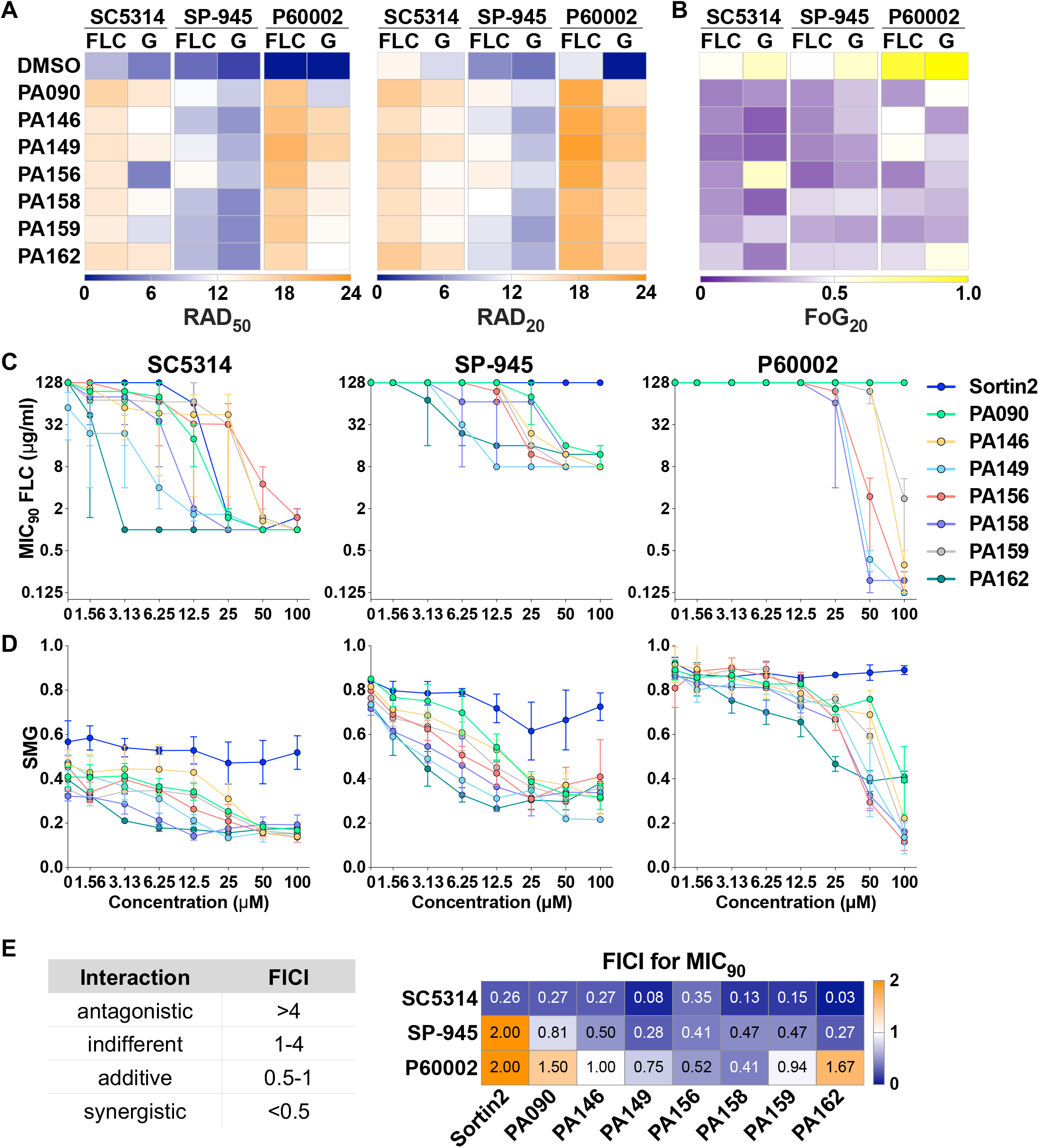
Impact of the seven 1,4-BZD/FLC drug combinations on FLC susceptibility, tolerance and cell survival of *C. albicans*. Compounds were screened using FLC (25 µg) disk diffusion assays on YPD plates containing 100 µM 1,4-BZDs or the same volume of DMSO (vehicle). Panels show FLC susceptibility (RAD50 and RAD20, A) and tolerance (FoG20, B) following 48 h of growth. After 48 h, FLC disks were replaced with glucose disks (G, 8 mg), grown for an additional 48 h, and susceptibility and tolerance were quantified again. Heatmaps show average values for 3 or more biological replicates. (C-D) Checkerboard assays showing the impact of drug combinations of FLC (starting at 128 µg/mL) with the seven 1,4-BZDs and sortin2 (starting at 100 µM) on FLC susceptibility (MIC90, C) and tolerance (SMG, D) for *C. albicans* isolates SC5314, SP-945 and P60002. (E) FICI90 scores for combinations of potentiators and FLC based on MIC90 values. Left-side table indicates the type of FLC-potentiator interaction based on the FICI score. Heatmap shows average values from 3-4 biological replicates.

To more broadly examine the potentiation phenomenon, we carried out azole potentiation experiments with fungal cells grown under standard CLSI conditions (47) in RPMI media (buffered to pH 7) and incubated at 30, 35 or 37°C (Fig. S3). Under these conditions, the 1,4-BZDs significantly enhanced FLC susceptibility in a similar fashion to that observed with YPD media (MIC_50_, Fig. S3A). We found it curious that the impact of potentiators on tolerance were less pronounced on RPMI media than in YPD media (SMG, Fig. S3B). It is also noteworthy that four of the seven 1,4-BZDs (PA156, PA158, PA159 and PA162) exhibited significant potency against strains grown in RPMI irrespective of the growth temperature (Fig. S3).

### 1,4-BZDs are synergistic with FLC against resistant and tolerant strains

After identification of seven 1,4-BZDs that potentiate FLC activity against *C. albicans*, we sought to determine whether the compounds were acting additively or synergistically with this antifungal. Accordingly, we performed the standard checkerboard assay in liquid growth media that is used for assessing the mode of potentiation (25). In these assays, we measured the influence of 1,4-BZDs on both FLC susceptibility (MIC_50_ and MIC_90_) and FLC tolerance (supra-MIC growth (14), SMG) starting at 128 µg/mL for FLC and 100 µM for the 1,4-BZDs in 2-fold serial dilutions (Fig. S4A). Sortin2 was included as a control in these experiments (32). These assays revealed significant FLC susceptibility restoration with the two resistant *C. albicans* isolates (SP-945, P60002) and FLC potentiation with the tolerant isolate (SC5314, Fig. 3C-D, S4A and S5A). PA149, PA158 and PA162 effected the largest changes in susceptibility and tolerance across the 3 strains (Fig. 3C-D). In fact, combinations of FLC with 6.25-25 µM of these 1,4-BZDs lowered MICs 100 to 1000-fold (Fig. 3C and S5A). In contrast, sortin2 showed minimal FLC potentiation across strains, with significant effects observed only for SC5314 (drug-tolerant) at 25 µM (MIC_90_, Fig. 3C). With respect to azole tolerance, the strongest potentiators were PA149, PA156, PA158 and PA162 (Fig. 3D).

For all seven 1,4-BZDs, we calculated fractional inhibitory concentration indices (FICIs) at both 50% and 90% inhibition levels to determine if they were acting additively or synergistically with FLC (described in Tables S2-4). For most interactions examined, FICI_90_ scores were <0.5 against both drug-resistant and drug-tolerant strains, indicating synergistic interactions between FLC and 1,4-BZDs. Synergies were most pronounced when the MIC_90_ was measured for resistant strain SP-945 and tolerant strain SC5314 (Fig. 3E). For P60002, owing to its high resistance levels (MIC_50_ >128 µg/mL FLC), synergistic interactions were most obvious at 50% inhibition (FICI_50_, Fig. S5B). Overall, these assays revealed that FLC potentiation by the 1,4-BZDs is synergistic and that PA149, PA158 and PA162 were particularly potent at restoring FLC susceptibility to drug-resistant and drug-tolerant *C. albicans* strains.

### Combinations of 1,4-BZDs and FLC are fungicidal

The apparent potentiation of FLC activity by the 1,4-BZDs raised questions about the manner of growth inhibition. Specifically, we wanted to determine if the combinations were fungistatic or fungicidal. Thus, after growth on FLC with 1,4-BZDs, the FLC disk was replaced with a glucose disk, and RAD_50_, RAD_20_ and FoG_20_ were measured again after 48 h of growth (Fig. 2C). Growth rebound in the initial zone of FLC inhibition upon removal of the antifungal was a marker of stasis, whereas absence of growth indicated cidality. We found that growth on glucose disks resulted in modest differences in RAD_50_, RAD_20_ and FoG_20_ values across the seven 1,4-BZDs (Fig. 3A-B and S2C). This contrasted observations with previously reported potentiators, which elicited mostly fungistatic effects (Fig. S2A-B).

Cidality was further studied via time-kill experiments wherein fungal cells were incubated over 24 hours in liquid media supplemented with a combination of FLC and potentiators and enumerated at different time points. Control experiments were carried out in media supplemented with either DMSO (no drug) or FLC alone. In all experiments, aliquots of cell cultures were seeded on YPD plates to quantify colony forming units (CFUs) at 0, 4, 8, 12, and 24 h (Fig. 4A). In control experiments with DMSO or FLC alone, CFU measurements revealed that all three isolates significantly increased in density over the course of the incubation period (Fig. 4B). In sharp contrast, treatment with drug combinations of FLC and 1,4-BZDs resulted in up to a 260-fold reduction in the number of live cells relative to the starting inoculum (with an average reduction across compounds of 43-fold and 46-fold for SC5314 and P60002, respectively, Fig. 4B). We were surprised to that the potentiator-drug combinations effected a much weaker cidal effect with against SP-945, with CFU counts closer in number to those observed in experiments with FLC alone (Fig. 4B). These observations indicate that drug combinations of FLC and 1,4-BZDs have potent fungicidal effects, but these effects vary in intensity in a strain-dependent manner.

**Figure 4.**
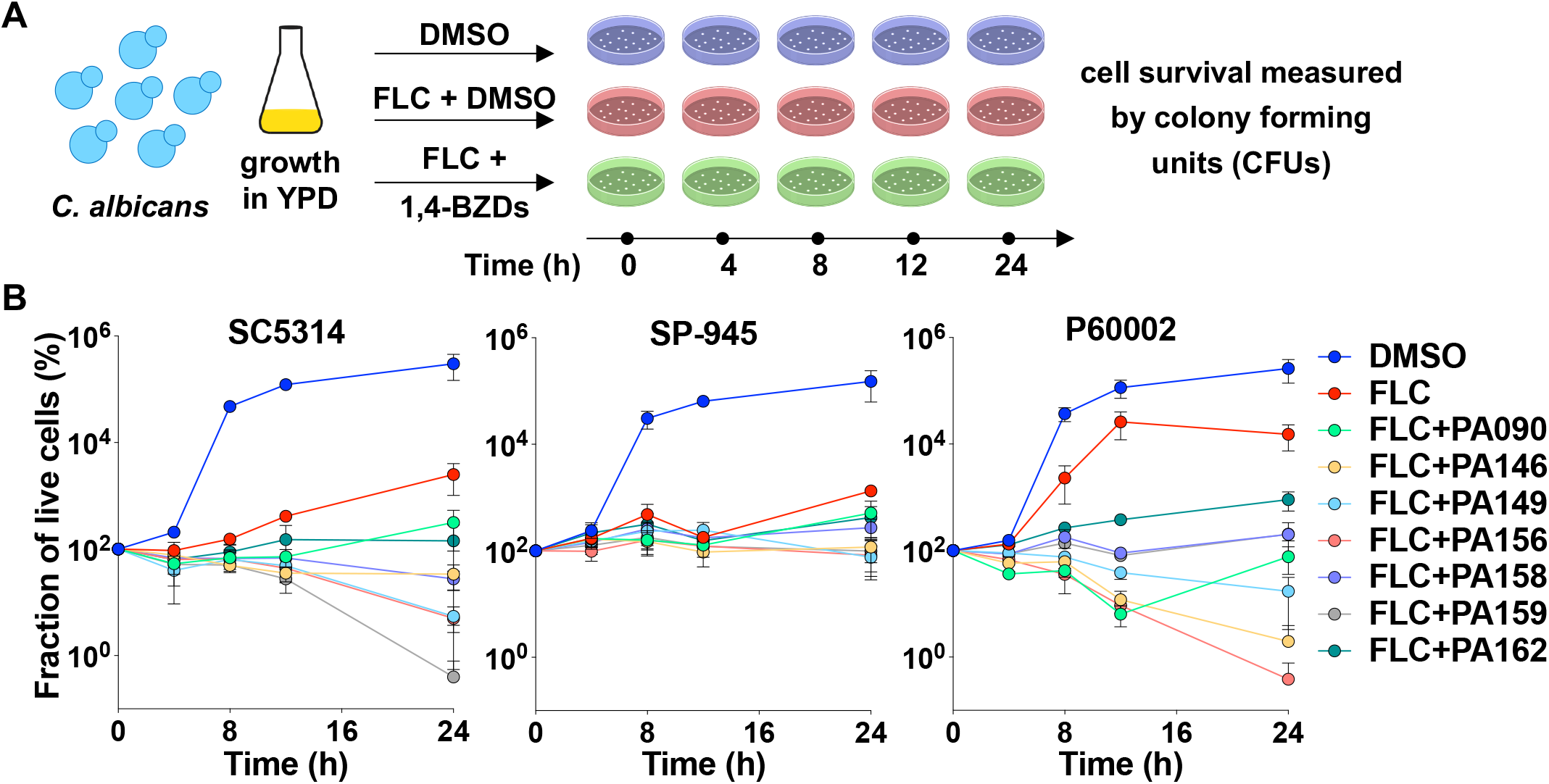
Time-kill assays in which *C. albicans* strains were cultured in YPD media with DMSO (control), FLC (128 µg/mL) or drug combinations of FLC (128 µg/mL) and 1,4-BZDs (100 µM) over 24 h. Cells were plated at 0, 4, 8, 12 and 24 h during growth for CFU determination (A). The percent of live cells incubated with different treatments is shown over time relative to the starting inoculum measured at 0 h (B). Plots show an average of three independent experiments, error bars show ± SEM.

### 1,4-BZDs are not intrinsically toxic

Negative control experiments in the checkerboard assays (those lacking FLC) indicated that the 1,4-BZDs alone did not have any inherent antifungal activity. To further explore these observations, we examined the influence of the potentiating 1,4-BZDs alone on fungal growth. Specifically, we monitored the growth of the three *C. albicans* isolates in liquid media supplemented with 1,4-BZDs at both 30°C and 37°C for 24 h (Fig. 5A). Sortin2 (32) was also included for comparison in this experiment. We found that fungal growth was not affected by either sortin2 or any of the PA90, PA146, PA156, PA158 or PA162 compounds tested, even at the highest concentrations used in FLC restoration/potentiation assays (100 µM, Fig. 5B). Moreover, calculation of doubling times in the presence of 1,4-BZDs revealed no significant differences relative to the no drug condition across the tested compounds (Fig. 5C). These observations recapitulated those in the disk diffusion assays, in which the outer edges of the agar plates containing 1,4-BZDs showed no inhibition of *C. albicans* growth relative to regions close to the FLC disk (Fig. S2C) and point to negligible fungistatic or fungicidal effects of 1,4-BZDs in the absence of FLC.

**Figure 5.**
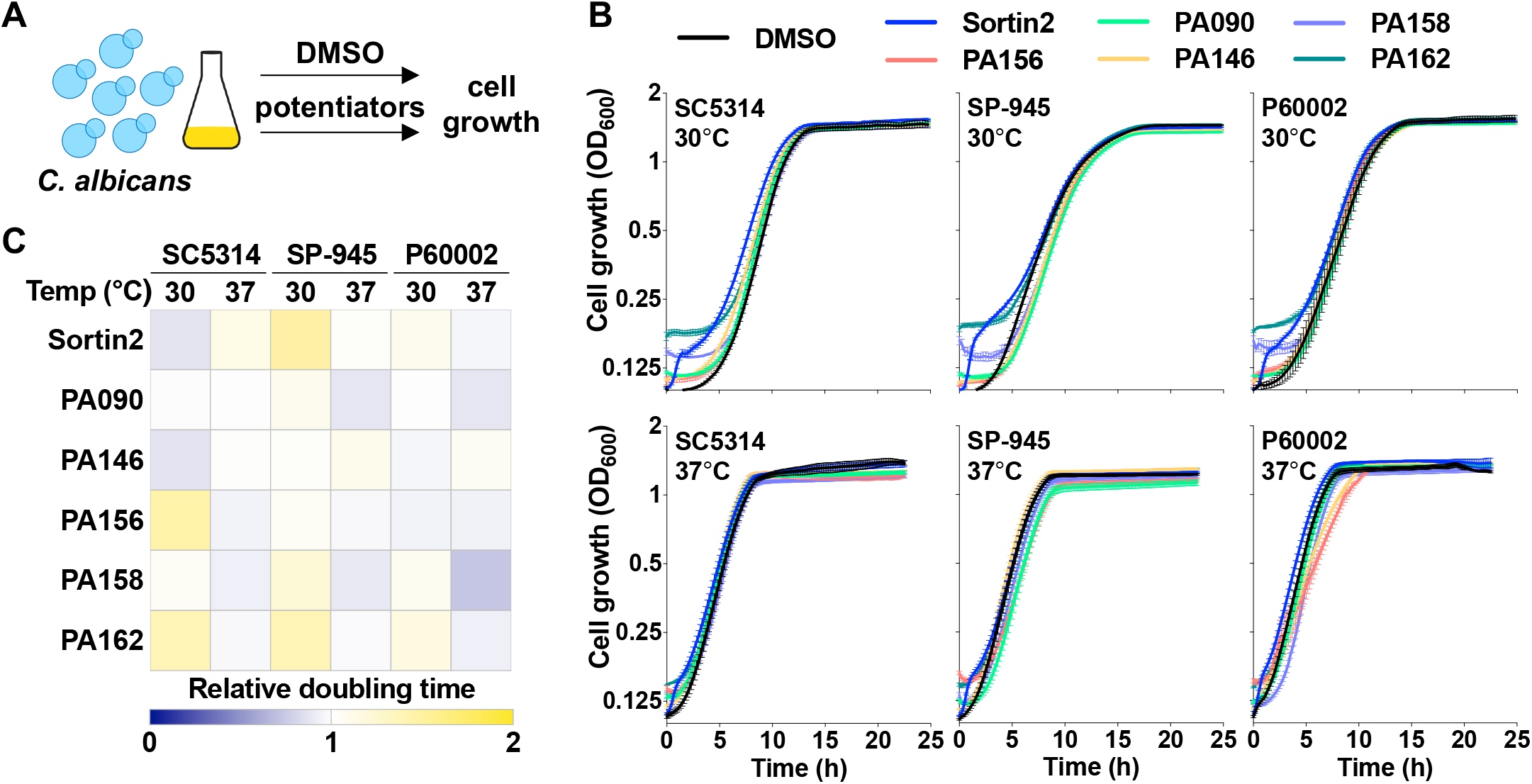
Impact of potentiators on *C. albicans* cell growth. (A, B) Growth curves of *C. albicans* isolates (SC5314, SP-945 and P60002) on YPD at 30 and 37 °C in the presence of DMSO, sortin2 (100 µM), or 1,4-BZDs (100 µM). Lines show average values from 3 biological replicates, error bars represent ± SEM. (C) Heatmap shows relative doubling times of *C. albicans* cells in the presence of potentiators, normalized to DMSO controls.

In an effort to assess the potential of the 1,4-BZDs for safe use as therapeutics in mammals, we tested their cytotoxicity against human peripheral blood mononuclear cells (PBMCs). Blood samples were collected from healthy donors and cells were isolated for culturing either with DMSO (non-toxic vehicle), 1,4-BZDs, or Triton (a cytotoxic detergent). After incubation with the 1,4-BZDs and the controls, the PBMCs were processed to measure the release of LDH (lactate dehydrogenase, an indicator of cytotoxicity). We were encouraged to find that most 1,4-BZDs were weakly cytotoxic at concentrations at which they potentiated FLC activity against *Candida* (5-25 µM, Fig. 6A). Indeed, incubation of PBMCs with PA158, PA159 or PA162 resulted in minimal or undetectable levels of LDH release at concentrations of up to 100 µM. As PA159 and PA162 were the least cytotoxic, these compounds were also tested for their ability to damage red blood cells (RBCs) by measuring hemolytic titers following incubation of RBCs with 1,4-BZDs. We found that, at 50 µM, both compounds exhibited hemolytic titers similar to those of negative controls (DMSO or PBS). Encouragingly, these titers modestly increased when 1,4-BZDs were used at high concentrations (200 µM, Fig. 6B). These data indicate that the 1,4-BZDs do not exert obvious host toxicities that would preclude their further optimization for mammalian use.

**Figure 6.**
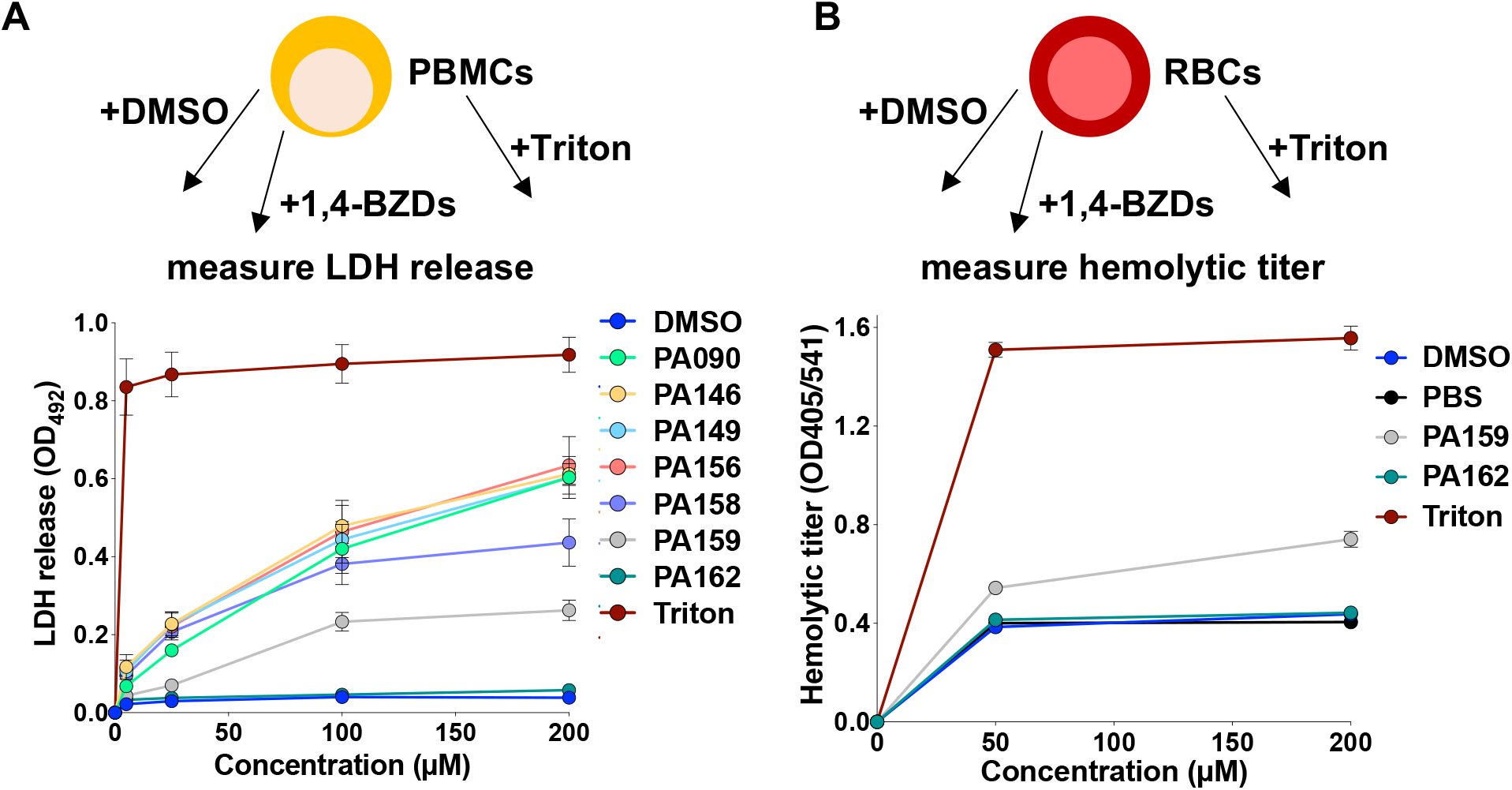
Cytotoxicity of 1,4-BZDs on human peripheral blood mononuclear cells (PBMCs, A). Lytic activity of the 1,4-BZDs was assessed with red blood cells (RBCs, B). PBMCs and RBCs were incubated with DMSO, PBS, Triton (positive control), or 1,4-BZDs at different concentrations and treatment toxicity was inferred by quantifying LDH release for PBMCs (A) or by measuring hemolytic titers for RBCs (B). Assays were performed with blood samples from two donors, each performed with two technical replicates, error bars show ± SEM.

### 1,4-BZDs potentiate azole antifungal activity in several *Candida* species

To gain further insights into FLC potentiation, we assessed the capacity of the 1,4-BZDs to potentiate FLC activity against phylogenetically diverse *Candida* species and more distantly related yeast, including *Candida dubiliniensis, Candida tropicalis, Candida parapsilosis, Candida metapsilosis, Candida orthopsilosis, Candida auris, Candida duobushaemulonii, Candida haemulonii, Candida krusei, Candida lusitaniae, Candida glabrata, Kodameae ohmeri* and *Saccharomyces cerevisiae*. Initially, we determined the baseline FLC susceptibilities of isolates of these species in liquid growth media. Twenty-four of the 40 isolates examined had MIC_50_s greater than 4 μg/mL, being classified as FLC-resistant by current MIC clinical breakpoints (except for *C. glabrata* where this breakpoint is 16 μg/mL (48) (marked with R, Fig. 7A). Twenty-two of the 40 isolates had baseline SMG levels greater than or equal to 0.5 indicating that these isolates were tolerant to FLC (marked with T, Fig. 7B). Considering these parameters, we assessed the capacity of the 1,4-BZDs (PA90, PA149, PA156, PA158 and PA162) to potentiate FLC activity against the isolates at 100 µM. The assays revealed that selected 1,4-BZDs were able to potentiate FLC against isolates of *C. dubliniensis, C. tropicalis, C. haemulonii, C. duobushaemulonii*, *K. ohmeri*, *C. glabrata* and *S. cerevisiae* (Fig. 7A-B). In contrast, the 1,4-BZDs did not potentiate FLC against *C. auris*, *C. krusei, C. lusitaniae, C. parapsilosis, C. metapsilosis*, and *C. orthopsilosis* (Fig. 7A-B). Interestingly, potentiation activities correlated with the phylogenetic relatedness of these species to *C. albicans*. The 1,4-BZDs displayed higher FLC potentiation against close relatives of *C. albicans* (*C. dubliniensis* and *C. tropicalis*) relative to more phylogenetically distant species (*e.g*., *C. auris*) (Fig. 7C).

**Figure 7.**
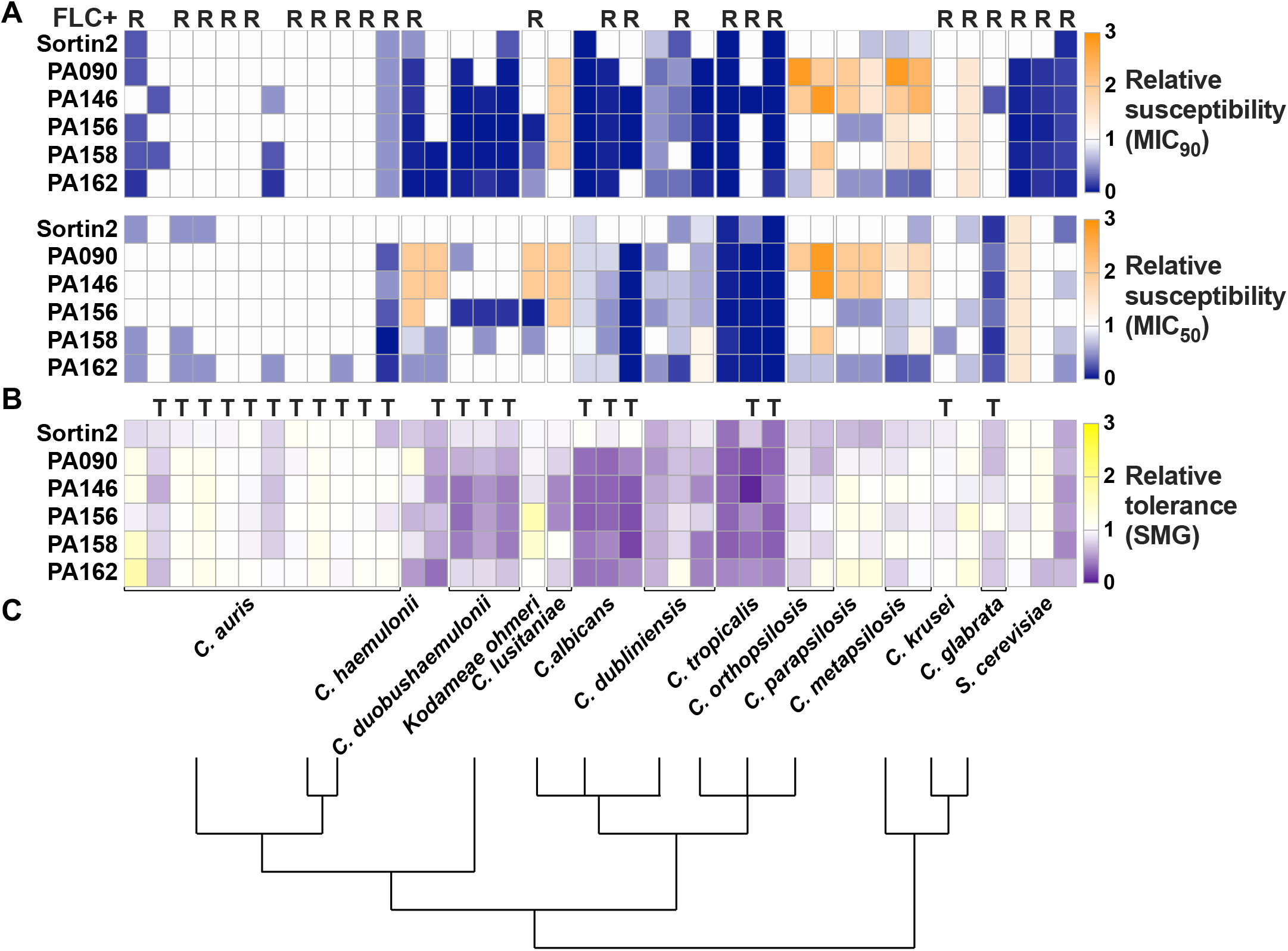
Changes in FLC susceptibility (MIC50 and MIC90, A) and tolerance (SMG, B) of other yeast species using combinations of potentiators (100 µM) and FLC (from 0 to 256 µg/mL). Values are shown relative to DMSO controls. R, FLC resistant isolates (MIC50 ≥ 16 µg/mL for *C. glabrata* and MIC50 ≥ 4 µg/mL for the rest of the species); T, FLC tolerant isolates (SMG ≥ 0.5). Heatmaps show average values from 3-4 biological replicates. (C) Lines indicate phylogenetic relationships between different species, adapted with permission from (112).

### 1,4-BZDs potentiate the azole class of antifungal drugs

Due to the synergistic potentiation of FLC activity by the 1,4-BZDs, we queried the specificity of the phenomenon by determining if the compounds could potentiate the activities of other antifungal agents. Initially, we explored the capacity of PA149, PA156, PA158, PA159 and PA162 (100 µM) to potentiate other azole drugs, including itraconazole (ITC), ketoconazole (KCA), posaconazole (POS) and voriconazole (VOR) in disk diffusion assays with *C. albicans* strains SC5314 and P60002. The 1,4-BZDs tested restored the susceptibility of the resistant isolate P60002 (Fig. 8A) and suppressed the tolerance of both strains (Fig. 8B). Thus, azole/potentiator combinations led to increased susceptibility (higher RAD_50_ and RAD_20_, Fig. 8A) and decreased tolerance (lower FoG_20_, Fig. 8B). Again, the 1,4-BZDs compounds outperformed sortin2 which showed minimal potentiation in combination with other azoles.

**Figure 8.**
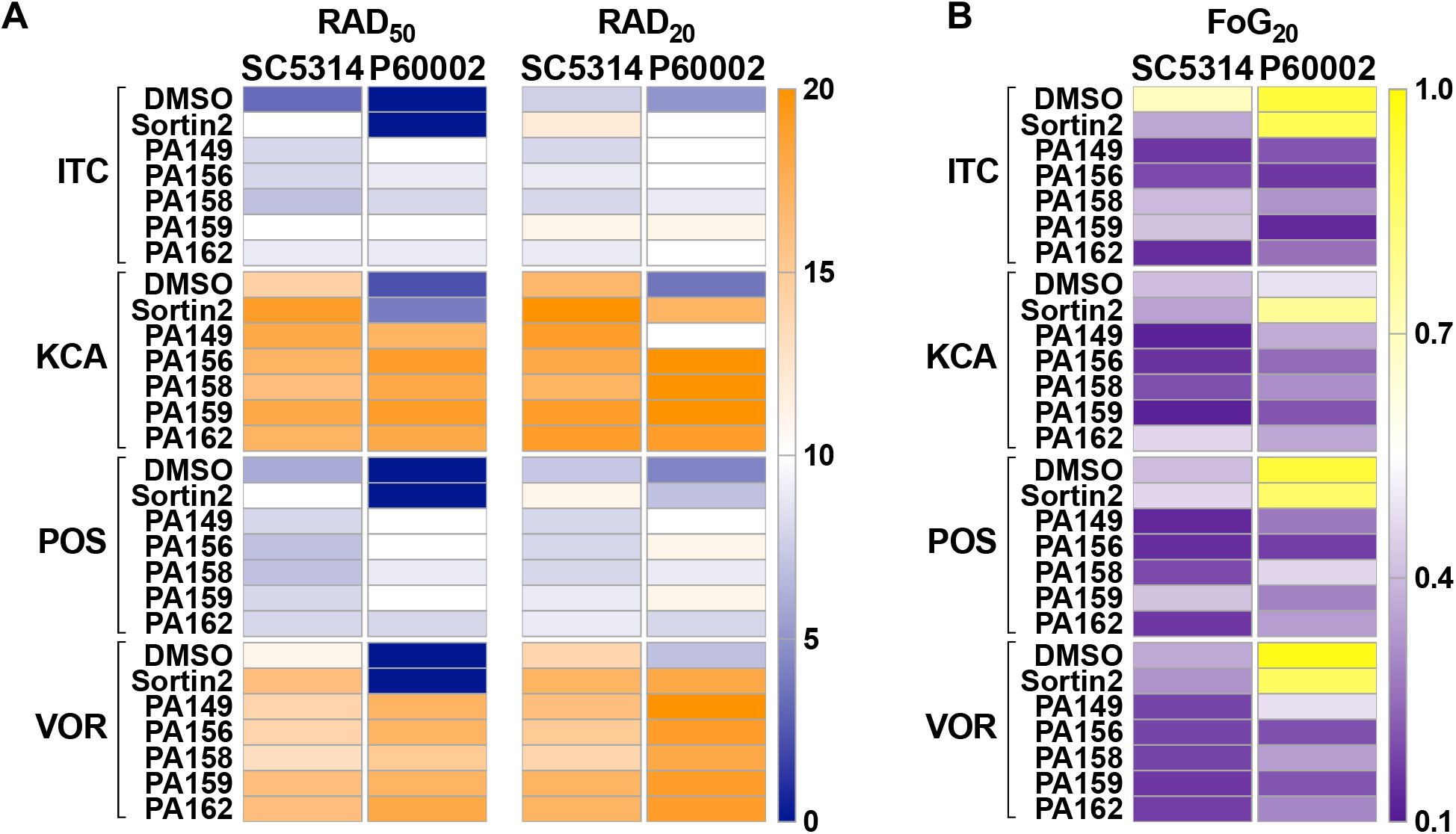
1,4-BZDs potentiate other azole antifungals. Heatmaps show drug susceptibility (RAD50 and RAD20, A) and tolerance (FoG20, B) when *C. albicans* isolates were plated with combinations of potentiators and other azole drugs. Compounds were screened using disk diffusion assays on YPD plates containing 1,4-BZDs (100 µM), sortin2 (100 µM), or DMSO (no drug) and disks of itraconazole (ITC, 50 µg), ketoconazole (KCA, 10 µg), posaconazole (POS, 5 µg), or voriconazole (VOR, 1 µg). Heatmaps show average values from 3 biological replicates.

Three of the potentiating 1,4-BZDs (PA90, PA146 and PA162) were also tested for their capacity to potentiate non-azole antifungals, including flucytosine (AFY), griseofulvin (AGF), amphotericin B (AMB), caspofungin (CAS) and nystatin (NY). These drugs have structures distinct from those of the azoles and do not target ergosterol biosynthesis. Their mechanisms of action include inhibition of DNA synthesis (AFY), perturbation of the fungal cell membrane (AMB, NY), disruption of cell wall β-1,3 glucan synthesis (CAS), and microtubule disassembly (AGF) (49, 50). In a marked difference from the azoles, we found that the 1,4-BZDs did not significantly potentiate the non-azole antifungal drugs (Fig. S6A-B). These results suggest that the 1,4-BZDs could be broadly used as potentiators of the azole anti-fungal drugs.

### Cidality of 1,4-BZDs requires perturbation of membrane constitution

The azole-specific potentiation by the 1,4-BZDs suggested that the potentiation may be contingent on perturbation of biogenesis of ergosterol or other constituents of the fungal cell membrane. Accordingly, we assessed the capacity of the 1,4-BZDs to potentiate small molecules that perturb the biosynthesis of cell membrane components. We expanded our analysis to include compounds that are not clinically used as anti-fungal drugs. We selected terbinafine (TERB), which targets Erg1 (51) upstream of Erg11, and fenpropimorph (FEN), which targets Erg2 and Erg24 (52), both downstream of Erg11 (the target of FLC, Fig. 9A). We also included myriocin (MYO), an inhibitor of serine palmitoyltransferase, involved in the synthesis of sphingolipids (53, 54), which are integral components of fungal membranes. Initially, we measured the activities of the aforementioned molecules against the three *C. albicans* isolates and found that the susceptibility (MIC_50_) and tolerance (SMG) levels did not correlate with those observed for FLC (Fig. S7A) or with the pathway targeted. For example, isolate P60002 is highly resistant to FLC but highly susceptible to terbinafine and intermediately susceptible to fenpropimorph, even though all three drugs target the same pathway (Fig. S7A). Because FLC, terbinafine, and fenpropimorph all target ergosterol biosynthesis, we anticipated that they would all be potentiated by the 1,4-BZDs. Indeed, the compounds significantly potentiated the activities of the modulators of ergosterol biosynthesis; however, the degree of potentiation did not reach that observed with FLC, especially with respect to tolerance (SMG, Fig. 9B). Consistent with the apparent dependence of the potentiation on perturbation of ergosterol biosynthesis, we found that a *Candida* strain lacking *ERG5*, encoding the sterol desaturase involved in the last step of ergosterol synthesis (55), was more sensitive to 1,4-BZDs than it corresponding parental strain (Fig. 9A-B). In contrast, deletion of *UPC2*, encoding a key transcriptional regulator of ergosterol biosynthetic genes (56), did not impact the activities of the potentiators. We reason this could be due to compensatory upregulation of ergosterol biosynthesis genes by other transcription factors (57). In parallel assessments of the correlation between the action of 1,4-BZDs and perturbation of membrane constitution, we carried out potentiation experiments with myriocin, an antifungal inhibitor of sphingolipid biogenesis. We were intrigued to find that the activity of myriocin was potentiated by the seven 1,4-BZDs. Curiously, the compounds increased susceptibility of the strains to myriocin without significantly altering tolerance (Fig. 9A-B). Though they do not reveal the potentiation mechanism, these experiments suggest that activity of the 1,4-BZDs is dependent on the constitution of the cell membrane.

**Figure 9.**
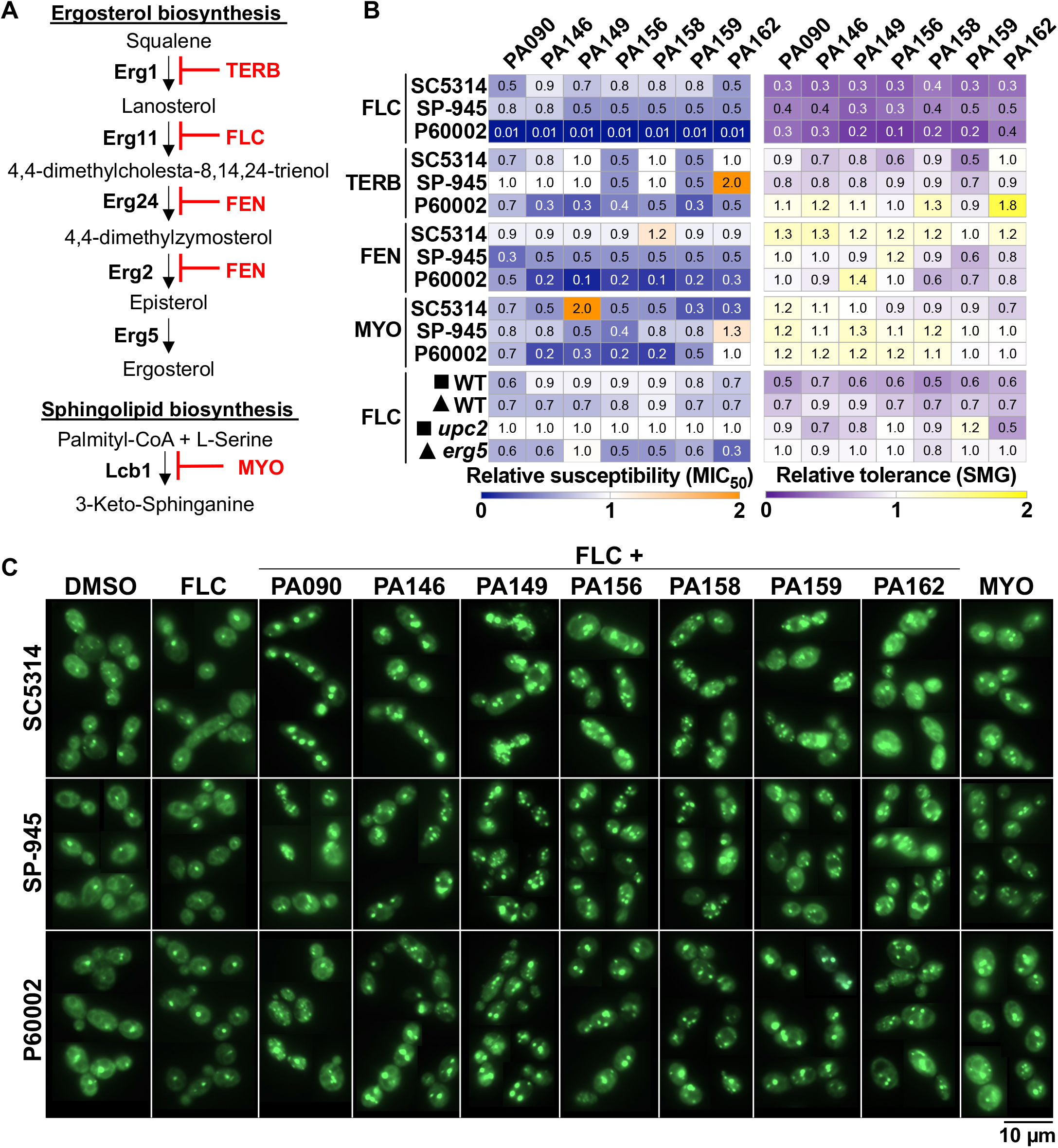
1,4-BZDs potentiate inhibitors of ergosterol and sphingolipid biosynthesis. (A) Terbinafine (TERB) and fenpropimorph (FEN) target ergosterol biosynthesis via Erg1 and Erg24/Erg2, respectively, while myriocin (MYO) targets sphingolipid biosynthesis via Lcb1. (B) Changes in FLC, TERB, FEN or MYO susceptibility (MIC50) and tolerance (SMG) when *C. albicans* isolates were treated with combinations of 1,4-BZDs (at 100 µM) and serial dilutions of FLC (0 to 128 µg/mL), TERB (0 to 32 µg/mL), FEN (0 to 32 µg/mL), or MYO (0 to 128 µg/mL), relative to DMSO controls. FLC data is included for reference. Mutants of Upc2, the central regulator of ergosterol synthesis genes, and Erg5, a cytochrome P450 enzyme catalyzing the last step in ergosterol biosynthesis, were also tested with FLC and 1,4-BZDs; symbols next to strain names denote their corresponding lineage/WT controls, constructed in the SC5314 background. All values are shown relative to DMSO controls and indicate the average from 3-4 biological replicates. (C) *C. albicans* isolates stained with a Bodipy dye following 4 h incubation with either DMSO, FLC (128 µg/mL), MYO (32 µg/mL), or combinations of FLC (128 µg/mL) and 1,4-BZDs (100 µM). Images show representative cells for the respective conditions, at 60X magnification. Scale bar, 10 µm.

### Combinations of 1,4-BZDs and FLC alter lipid homeostasis

Given the capacity of 1,4-BZDs to potentiate antifungals that inhibit the biogenesis of certain cell membrane constituents, we also sought to assess the influence of the compounds on lipid homeostasis. Towards this end, we used a molecular probe that stains neutral and non-polar lipids (BODIPY 493/503, Invitrogen) and has been previously validated in this species for observing lipid droplets (58). These organelles are dynamic lipid storage structures that support lipid homeostasis and help mediate cellular stress responses (59). Treatment of *C. albicans* cells with FLC alone did not reveal changes in lipid droplet size and distribution relative to non-treated cells (DMSO control), with most cells displaying single lipid droplets and uniform cytoplasmic staining (Fig. 9C). In contrast, treatment with myriocin, an inhibitor of sphingolipid biosynthesis used here as a positive control, resulted in the accumulation of lipid droplets indicating disruption of lipid homeostasis (Fig. 9C). Surprisingly, treatment with combinations of FLC and 1,4-BZDs resulted in a large proportion of cells displaying an increase in both the size and number of lipid droplets per cell, suggesting that the combination of FLC and 1,4-BZDs interferes with lipid homeostasis (Fig. 9C). While these observations do not clarify the mechanism of azole potentiation, it is clear that combinations of FLC and 1,4-BZDs radically impact lipid homeostasis, leading to significant changes in the structure and organization of membrane-bound compartments in fungal cells.

### Absence of Cdr1 efflux pump negatively affects potentiation by 1,4-BZDs

Given that 1,4-BZDs potentiate azoles, it is possible that they could either directly or indirectly enhance the intracellular accumulation of azoles to an extent that they are active against drug-resistant and drug-tolerant strains. Mechanistically, intracellular accumulation of azoles could occur via enhancement of uptake or inhibition of efflux. The latter phenomenon underlies the resistance of *Candida* to many drugs including the azoles (60–63). Indeed, multiple genes encoding efflux pumps have been associated with FLC resistance (60, 61, 64) and their impact on potentiation could inform on the mechanism of action of 1,4-BZDs. Therefore, we tested the capacity of 1,4-BZDs to potentiate FLC activity against mutants lacking ABC (ATP-Binding Cassette; *e.g*., *ADP1*, *CDR1*, *CDR2*, *MDL2*, *SNQ2*) or MSF (Major Facilitator Superfamily; *e.g*., *FLU1*, *MDR1*) transporter genes (63). These transporters are primarily regulated by Tac1, Mrr1 and Upc2 (the latter also being a central regulator of ergosterol biosynthesis) (56, 65, 66); therefore, we also included strains lacking these transcription factors in our analyses (Fig. 10A). As expected, we observed the increased FLC susceptibility of the strains lacking *CDR1* (either alone or in combination with other deletions) and the reduced FLC tolerance of strains containing *CDR1*, *CDR*2 or *FLU1* deletions (Fig. S7B). This aligns with reports that Cdr1 is a major azole efflux pump in *C. albicans* (63, 67), although compensatory effects were often observed between transporters or their regulators (Fig. S7B). Next, we examined FLC potentiation by 1,4-BZDs using MIC liquid assays wherein changes in both FLC susceptibility (MIC_50_) and tolerance (SMG) relative to no potentiator controls (DMSO). In several mutants, FLC potentiation by the 1,4-BZDs was altered relative to corresponding parental strains (WT). Strains lacking *FLU1*, both *CDR1* and *CDR2*, or *TAC1*, the transcription factor regulating the expression of *CDR1* and *CDR2* (65), showed increased susceptibility (decreased MIC_50_) to FLC-1,4-BZD combinations relative to control strains (Fig. 10B). The apparent conservation of FLC potentiation by the 1,4-BZDs in strains having and lacking efflux pumps implies that the mechanism of the phenomenon is predominantly efflux-independent.

**Figure 10.**
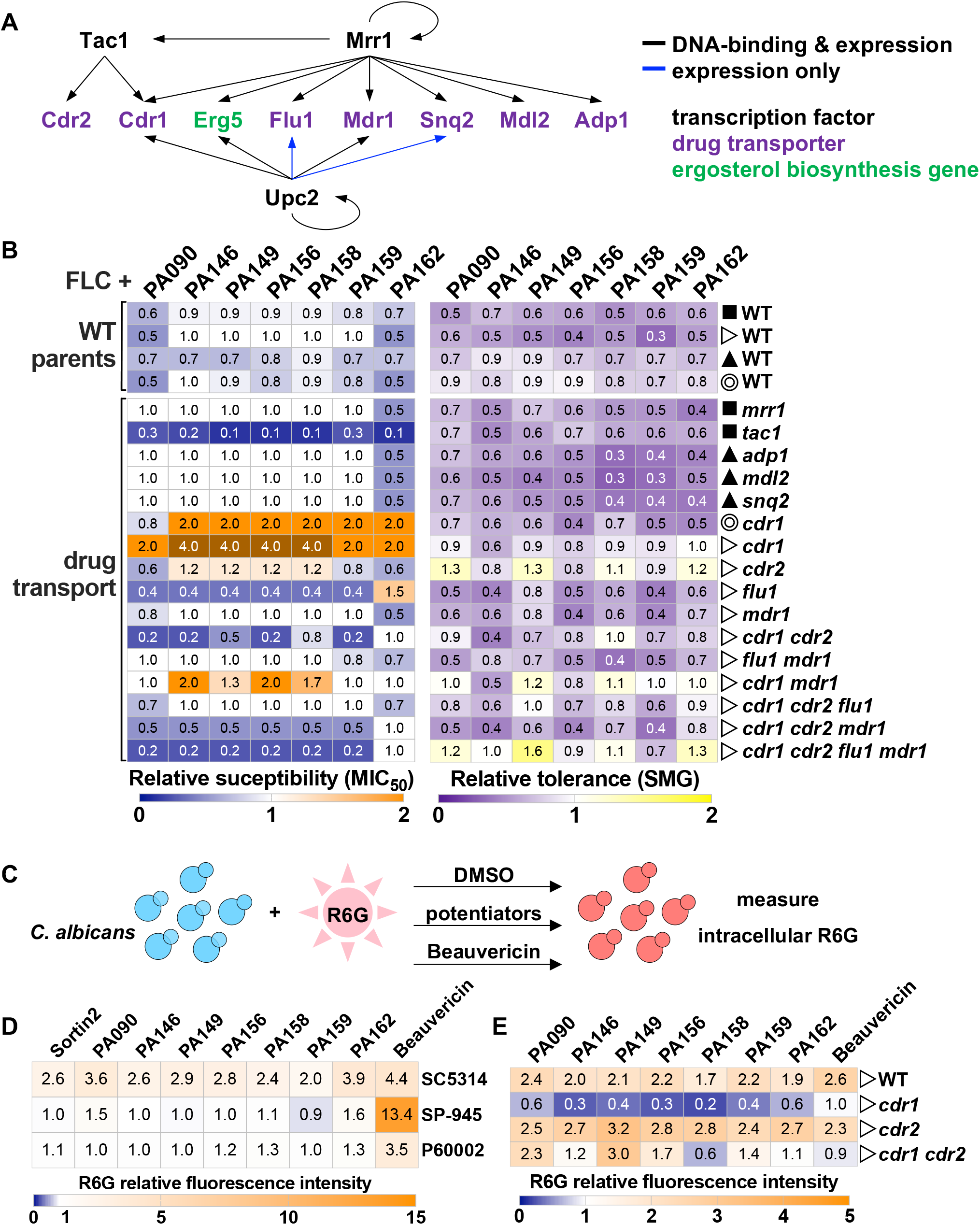
Impact of drug combinations on *C. albicans* mutants of drug transporters and their regulators. (A) Schematic shows transporters from the classes of ABC (ATP-Binding Cassette, Adp1, Cdr1, Cdr2 and Mdl2) and MSF (Major Facilitator Superfamily, Flu1 and Mdr1) as well as their corresponding transcription factor regulators (Tac1, Mrr1 and Upc2). (B) Impact of 1,4-BZDs on the FLC susceptibility (MIC50) and tolerance (SMG) of different single and combined deletion mutants. Symbols denote the corresponding WT strains for each mutant, all constructed in the SC5314 background. Heatmaps show average values from 3 biological replicates, relative to corresponding DMSO controls. (C) Impact of 1,4-BZDs on intracellular rhodamine 6G accumulation. (D-E) Rhodamine 6G fluorescence intensity levels were measured after 4 h incubation with rhodamine 6G (1 µg/mL) and treatment with either DMSO (no drug), sortin2 (100 µM), 1,4-BZDs (100 µM), or beauvericin (10 µg/mL, as positive control). Strains examined include SC5314, SP-945, P60002 (D), as well as *cdr1*, *cdr2*, *cdr1 cdr2* mutants with the corresponding WT parental strain (E). Heatmaps show average values from 3 or more biological replicates, relative to corresponding DMSO controls.

The surprising exception to efflux-independent FLC potentiation activity of the 1,4-BZDs is the increased MIC_50_ levels of two independently constructed *cdr1* mutants (Fig. 10B). It is not clear whether the Cdr1 transporter impacts the activity of the potentiators either directly or indirectly. These observations are difficult to explain given the complex roles that Cdr1 plays in fungal physiology. In addition to mediating azole efflux, Cdr1-3 are general phospholipid translocators in *C. albicans* and actively contribute to the biogenesis of the plasma membrane lipid bilayer, therefore influencing membrane integrity and phospholipid signaling (68). It is noteworthy that disruption of phospholipid homeostasis affects FLC susceptibility (69). The fact that both ergosterol and sphingolipids are essential for the correct localization of Cdr1 to lipid rafts in *C. albicans* adds further complexity to ascribing a role for Cdr1 in FLC resistance (70, 71). As drug combinations of FLC and potentiators alter lipid homeostasis, one could imagine that they could affect Cdr1 localization, thereby altering drug efflux (and therefore drug susceptibility). Given the complex relationship between Cdr1, membrane integrity and FLC susceptibility, the results observed with *cdr1* do not lend to easy interpretation.

The multiplicity of *C. albicans* drug efflux pumps alongside their intricate regulatory interactions and roles in membrane integrity complicate the interpretation of the genetic analyses of the influence of 1,4-BZDs over efflux. Thus, we performed experiments wherein we directly compared the 1,4-BZDs with beauvericin, a known fluorescent efflux substrate and promiscuous efflux inhibitor which mainly acts on Cdr1 and Cdr2 pumps (62, 72, 73) (Fig. 10C). We found that beauvericin significantly enhanced R6G accumulation in strains SC5314, SP-945 and P60002, but accumulation was either unchanged or reduced upon treatment of the strains with the seven potentiators (as well as for sortin2, Fig. 10D). Specifically, 1,4-BZDs increased R6G levels in isolate SC5314 but did not impact its accumulation in strains SP-945 and P60002 (Fig. 10D), where the 1,4-BZDs were most effective at potentiating FLC activity (Fig. 9B). From these biochemical experiments, it can be argued that 1,4-BZDs do not influence drug efflux.

Because the potentiators had intriguing effects on strains lacking *CDR1*, we specifically examined R6G accumulation in those mutants. Consistent with previous reports, *cdr1*, *cdr2*, and *cdr1 cdr2* mutants displayed 1.7-2.7-fold higher R6G accumulation relative to the WT strain in the absence of treatment (Fig S7D). The 1,4-BZDs differentially influenced R6G accumulation in the *cdr* null strains. While the increase in R6G accumulation was similar for the WT and *cdr2* strains, the opposite trend was observed for *cdr1*, with R6G levels decreasing upon treatment with 1,4-BZDs (Fig. 10E). An intermediate phenotype was observed for the *cdr1 cdr2* double mutant (Fig. 10E). The paradoxical decrease in R6G accumulation in the *cdr1* null effected by 1,4-BZDs mirrors the enhancement of FLC resistance in the same genetic background (Fig. 10B). Apparently, the deletion of *CDR1* effects a physiological state that suppresses the intrinsic activity of the 1,4-BZDs.

### 1,4-BZDs are potent inhibitors of *C. albicans* filamentation

The absence of toxicity for the 1,4-BZDs did not rule out the possibility that the compounds elicit other effects on the physiology of *C. albicans*. Accordingly, we used microscopy to assess the influence of the compounds on fungal cell morphology. SC5314, which is a highly filamentous strain (33), exhibited extensive filamentation, with ∼89% of the population growing as hyphae or pseudohyphae following 2 h induction in liquid YPD media at 37°C (elongated cells, Fig. 11A-B). The other strains were less filamentous; only ∼40% of P60002 and ∼6% of SP-945 cells were filamenting under these conditions (Fig. 11A-B). We quantified cell morphologies of the *C. albicans* strains in the presence of two 1,4-BZDs, PA158 or PA162, which were selected due to their high potency (Fig. 3) and divergent chemotypes (Fig. 1B). Interestingly, treatment with 1,4-BZDs (at 100 µM) significantly reduced the number of hyphal and pseudohyphal cells for all *Candida* strains (Fig. 11A-B). In contrast, treatment with FLC alone (10 µg/mL) modestly affected filamentation, while a combination of 1,4-BZD (100 µM) and FLC (10 µg/mL) resulted in similar filamentation levels as those observed when the strains were treated with 1,4-BZDs alone (Fig. 11A-B). These results are consistent with our observations that 1,4-BZDs perturb membrane integrity and lipid biosynthesis which are essential for the development of filaments (74–76). Importantly, these observations are notable because transition to a filamentous morphology is known to impact pathogenicity and commensalism in *C. albicans* (74, 77, 78). By targeting lipid homeostasis, 1,4-BZDs could directly interfere with this key developmental process.

**Figure 11.**
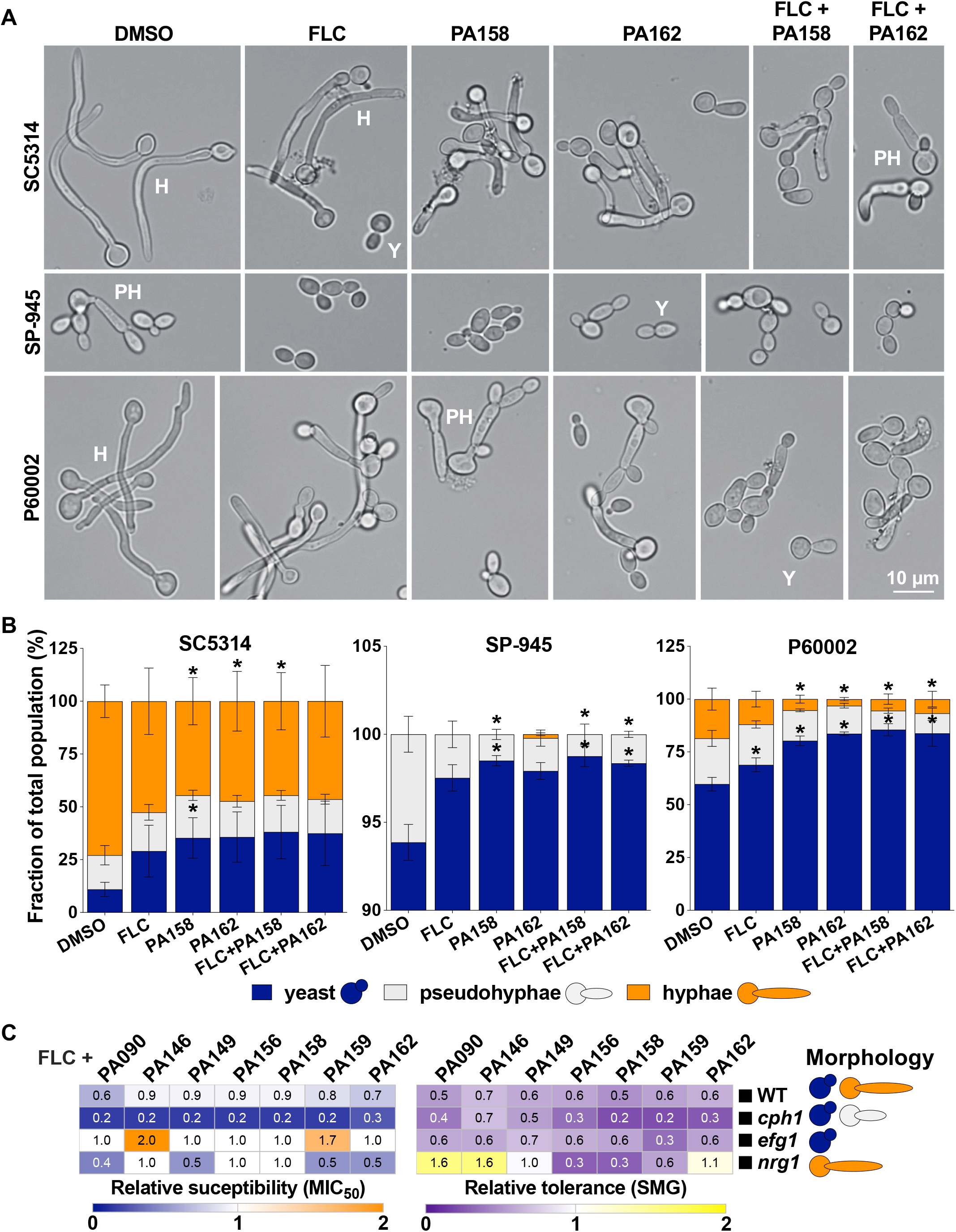
Impact of drug combinations on *C. albicans* filamentation. (A) Cell morphologies of strains SC5314, SP-945, P60002 treated with either DMSO, FLC (10 µg/mL), 1,4-BZDs (PA158 or PA162, 100 µM), or a combination of FLC (10 µg/mL) and 1,4-BZDs (100 µM). Cells were incubated in YPD at 37°C for 2 h prior to imaging. Images show representative cells for the respective conditions, at 40X magnification. Scale bar, 10 µm. (B) The fraction of each morphology was quantified for at least 200 cells per condition from 4 biological replicates. Asterisks indicate significant differences relative to DMSO, *t* test, * *P* < 0.05. Error bars represent ± SEM. (C) Impact of 1,4-BZDs (100 µM) on FLC susceptibility (MIC50) and tolerance (SMG) for mutants affecting cell morphology (see schematic on the right side) and corresponding parental strain. Changes in MIC50 and SMG are shown relative to levels observed with DMSO treatment. Heatmaps show average values from 3-6 biological replicates.

To determine whether the FLC potentiation activity of the 1,4-BZDs is dependent on filamentation, we assessed their activity in strains that were restricted to a particular cell morphology because they lacked key regulators of filamentation such as Nrg1, Efg1 and Cph1 (79–81). We found that 1,4-BZDs can alter the FLC susceptibility and tolerance of strains restricted to yeast or filamentous forms, similar to effects observed with the corresponding parental strain (WT, Fig. 11C). As is the case for FLC potentiation, the mechanism by which the compounds inhibit filamentation is unclear. Nevertheless, because filamentation is critical for *C. albicans* pathogenicity (78, 82) and its inhibition has emerged as a strategy for suppressing virulence (83–85), these observations suggest that the 1,4-BZDs could have utility beyond azole potentiation.

### 1,4-BZDs improve *G. mellonella* survival during *C. albicans* systemic infection

Finally, we examined the impact of 1,4-BZD/FLC combinations on *G. mellonella* survival during systemic *C. albicans* infection *G. mellonella* has been successfully utilized as a model for systemic infection for diverse pathogenic fungi. By extension, it is also frequently used for testing antifungal efficacy (86, 87). On these grounds, *G. mellonella* larvae were systemically infected with SC5314, SP-945 or P60002 and treated with either a vehicle control (PBS/DMSO at matching volume), FLC, 1,4-BZDs, or 1,4-BZDs/FLC combinations (Fig. 12A-E). The larvae were then incubated at 37°C and survival was monitored daily for 14 days. Control groups of larvae were used to assess the impact of these compounds on host survival in the absence of fungal infection (Fig. 12B). Treatment with FLC, sortin2 or 1,4-BZDs alone did not affect *G. mellonella* viability, indicating that these compounds are not overtly toxic to this organism (Fig. 12B). Experiments revealed that the different *C. albicans* isolates display distinct virulence dynamics and respond differently to FLC treatment. This was not surprising given the differences in genetic background, filamentation (Fig. 11), and azole susceptibilities (Fig. 2) of the three isolates. Infections by all strains resulted in significant host lethality, with median survival times (defined as the time at 50% group lethality) of 2, 1 and 4 days for SC5314, SP-945 and P60002, respectively (average of 2.33 ± 1.53 days). Treatment with FLC alone increased median survival times to 9.33 ± 1.53 days across strains (Fig. 12C-E). Strikingly, treatment with combinations of FLC and potentiators significantly increased survival relative to FLC alone (see detailed statistics in Table S5). Indeed, *Candida* infected larvae treated with FLC and either PA158 or PA162 had median survival times of 12.17 ± 0.76 and 12.33 ± 0.58 days, respectively. These observations also stand in contrast to results observed when infected larvae were treated with a combination of FLC and sortin2. In this case, median survival times increased to 10 ± 1 days (Fig. 12C-E) and this combination only improved the survival of larvae infected with SP-945 (see Table S5). Collectively, these observations indicate that the ability of 1,4-BZDs to potentiate FLC *in vitro* translates to *in vivo* infections.

**Figure 12.**
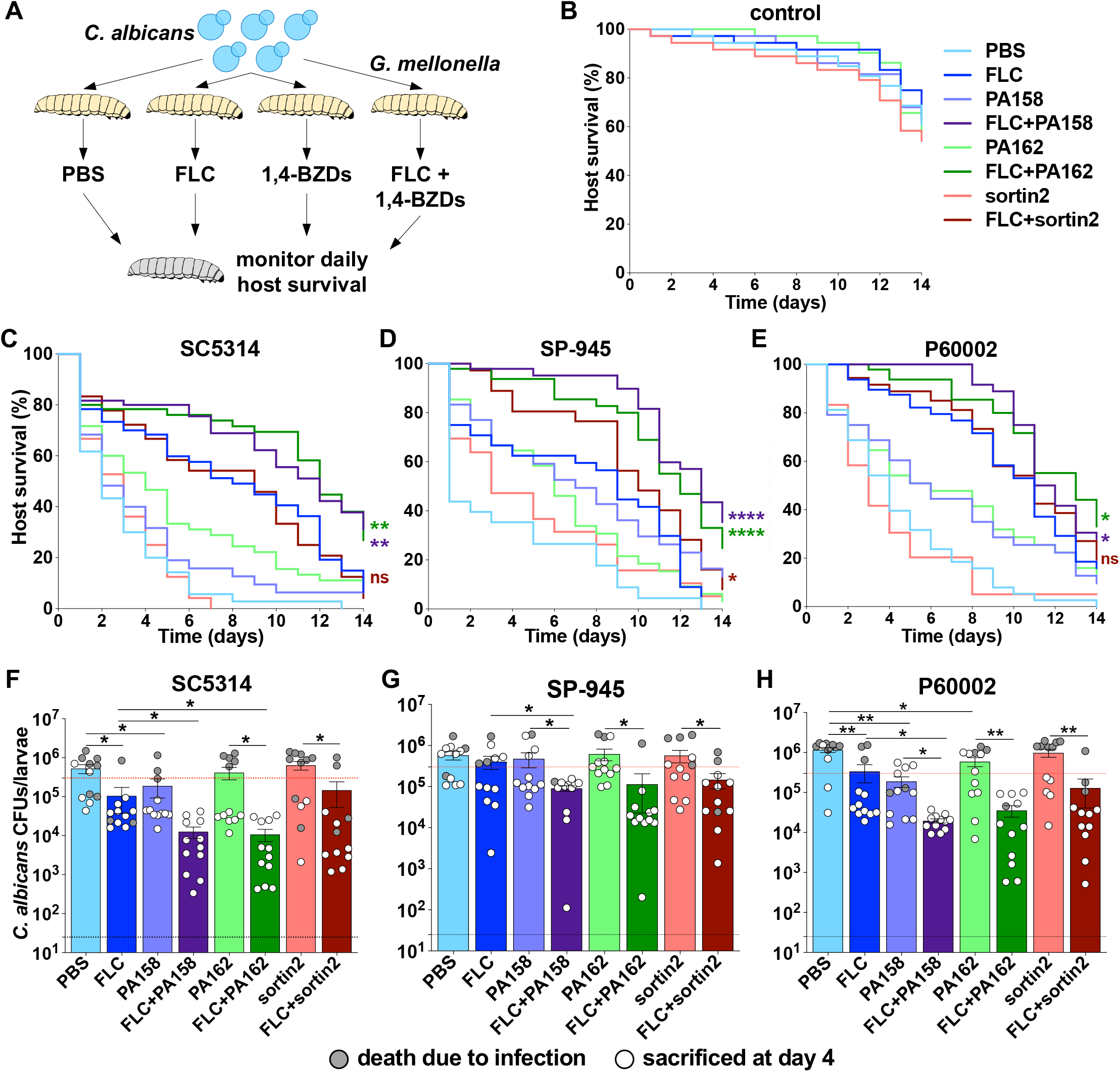
Impact of drug combinations on *G. mellonella* survival during systemic fungal infection. (A) Larvae of *G. mellonella* were systemically infected with *C. albicans* isolates (3 × 10^5^ cells/larva) and subsequently treated with either no drug (DMSO as vehicle), FLC (20 µg/mL), sortin2 (50 µM), 1,4-BZDs (PA158 or PA162, 50 µM), or a combination of FLC (20 µg/mL) and 1,4-BZDs (50 µM) in volumes of 10 µL/larva. Additional control groups of larvae were systemically injected with PBS and treated with either DMSO, FLC, sortin2, or 1,4-BZDs in the absence of fungal infection (B). Survival of larvae infected with SC5314 (C), SP-945 (D) or P60002 (E) was monitored daily for 10 days for 3-5 groups of larvae with 12 larvae per group for each condition (n = 36-60 larvae/group). Asterisks indicate significant differences for comparisons between FLC and drug combinations (FLC vs FLC + sortin2, FLC vs FLC + PA158, FLC vs FLC + PA162), colored according to the corresponding line, based on Log-rank (Mantel-Cox) tests. All statistical comparisons between each two groups are included in Table S5. (F-H) Fungal burdens recovered from larvae treated with single agents or drug combinations before or at 4 days post infection (n = 12 larvae/group). Grey dots show larvae that had succumbed to infection before or at 4 days, white dots show larvae sacrificed on day 4. Error bars represent ± SEM, asterisks indicate significant differences based on unpaired t-tests, * *P* < 0.05; ** *P* < 0.01.

Because fungal burdens are associated with mortality in this infection model (88, 89), we sought to assess the impact of 1,4-BZD/FLC combinations on this parameter. We measured fungal burdens 4 days post infection (or at the time of death for larvae which had succumbed to infection before this time point) by homogenizing the larvae and plating them for CFUs. We found that larvae that did not survive the initial 4 days displayed overall higher fungal burdens than those that survived (average of 12.6-fold higher, Fig. 12F-H), consistent with reports indicating that fungal load drives mortality in this organism. Importantly, we found that combinations of FLC and 1,4-BZDs significantly decreased fungal burdens relative to treatment with FLC alone (average of 10-fold and 7.5-fold lower fungal burdens for PA158 and PA162, respectively). This was in contrast with combinations of FLC and sortin2, which only decreased fungal burdens by 2-fold (Fig. 12F-H). Except for the P60002-infected larvae, differences between PBS and 1,4-BZD treated groups were not significant, emphasizing the synergistic effect of the drug combination for reducing growth of *C. albicans in vivo* (Fig. 12F-H). Together, these observations indicate that the *in vivo* FLC potentiation by 1,4-BZDs correlates with a reduction in fungal burdens.

It was also noteworthy that treatment of infected larvae with 1,4-BZDs alone significantly increased median survival (to 5 ± 2.64 days with PA158 and 5.33 ± 1.15 days with PA162 relative to 2.33 ± 1.53 days with DMSO and 3 days with sortin2 treatment). Because these differences could not be explained by a reduction of fungal burdens in these groups (Fig. 12F-H), the enhanced survival rates effected by treatment with 1,4-BZDs as single agents could be explained by their capacity to inhibit filamentation (Fig. 11). To test this hypothesis, we examined fungal cells present in *G. mellonella* tissue homogenates following staining with calcofluor white. This dye binds to chitin molecules in the fungal cell wall and therefore can be used to visualize fungal cells in tissues (90, 91). Detection of *C. albicans* proved challenging for larvae treated with FLC due to low fungal burdens, but >240 cells per host could be visualized for the rest of the groups. As expected, treatment with either PA158 or PA162 resulted in reduced filamentation in both SC5314 and P60002 infected larvae (Fig. S8A-B, an insufficient number of cells was examined for SP-945). We were gratified to find that the inhibition of virulence-associated filamentation by *C. albicans* observed *in vitro* was also observed in the *Galleria* infection model and was correlated with host survival (Figs. 11 and S8).

## DISCUSSION

The treatment of invasive candidiasis has been plagued by the emergence of drug resistant/tolerant species, a limited arsenal of antifungals and the lack of new antifungal molecules. Clinically, refractory invasive candidiasis is usually treated with single antifungal agents (11). The use of adjuvants that restore the activity of current antifungals could represent an effective therapeutic strategy. Here, we performed a small chemical screen in search of small molecules that could restore azole susceptibility to drug-tolerant and -resistant *Candida* species. We discovered seven unique 1,4-BZDs that potentiated FLC activity against azole-resistant and tolerant strains of *C. albicans.* Notably, these 1,4-BZDs are structurally distinct from all the benzodiazepine drugs, including anxiolytics and sedatives (*e.g*., diazepam, lorazepam, alprazolam, oxazepam, clonazepam). It is further noteworthy that the potentiators are distinct from other benzodiazepines reported to have antifungal activity either as single agents or in combination with azoles. For example, 3*H*-1,5-benzodiazepine derivatives were reported to have antifungal activity against *Aspergillus niger* and *C. albicans* (92). Hydroxylated analogs of 1,5-benzodiazepines (active against *Sporothrix schencki* and *Sporothrix brasiliensis*), substituted 1,5-benzodiazepines (active against *Aspergillus fumigatus* and *C. albicans*), and 2,4-disubstituted-1,5-benzodiazepines (active against *A. niger* and *C. albicans*), were reported to have IC_50_ of 0.4 – 900 μM (93–97). In 2014, Jassen Pharmaceuticals patented novel potent pyrrolo[1,2-a][1,4]-benzodiazepine analogs (IC_50_ 0.1 – 8 μM) with both *in vitro* and *in vivo* antifungal activities across diverse fungi including *C. parapsilosis*, *C. neoformans*, *A. fumigatus*, *Trichophyton rubrum* and *Scedosporium apiospermun* (98–100). A recent study described a tethered benzodiazepine and tetrazole scaffold having novel polypharmacological antifungal effects (101). This compound series exhibited potent activity against *C. albicans*, with >98% growth inhibition at 0.5 μM, while altering hyphal morphology, chitin deposition and membrane integrity (101). Finally, Midazolam, a 1,4-benzodiazepine that is clinically used as a sedative, was reported to have antifungal activity against planktonic and biofilm *Candida* cells. Though it did not synergize with FLC, this drug inhibited cell growth by inducing damage to the mitochondrial membrane and phosphatidylserine externalization (102). None of the above benzodiazepines share a substitution pattern and biological activities with the potentiators reported herein. Nevertheless, these precedents reinforce the idea that benzodiazepines are privileged scaffolds in drug discovery and support the need for further exploration of this class for novel antifungal pharmacology.

Remarkably, the 1,4-BZDs effected up to 1000-fold potentiation of FLC activity (MIC_50_) and cleared tolerance at concentrations of 6.25-25 µM. They potentiated the activity of a variety of other azoles (itraconazole, posaconazole, ketoconazole, voriconazole), but did not potentiate antifungal agents from other classes (caspofungin, amphotericin B, flucytosine, nystatin, griseofulvin). Though the azoles are known to be fungistatic in *Candida* species, we found that a combination of FLC with the potentiators was cidal. Checkerboard assays using 1,4-BZD/FLC combinations revealed that the compounds acted synergistically with FLC. The antifungal activity of this drug combination was especially notable because the 1,4-BZDs have no inherent antifungal activity. From a therapeutic standpoint, it is meaningful that the compounds exhibited negligible cytotoxicity to human cells and did not induce lysis of red blood cells.

At this stage, we do not have a molecular mechanism to describe how the 1,4-BZDs potentiate azole activity. The fact that they only potentiate azole activity against phylogenetically related species (*e.g*., *C. albicans, C. tropicalis*, *C. dubliniensis*) suggests that the mode of action is genetically encoded. In any case, several observations suggest that their mode of action both involves and depends on perturbation of lipid homeostasis. For example, the 1,4-BZDs selectively potentiated antifungals that act via inhibition of ergosterol and sphingolipids biosynthesis pathways, which are two important constituents of the fungal membrane. Moreover, we found that disruption of a gene underlying ergosterol biosynthesis increases susceptibility to combinations of FLC and potentiators. In the same vein, it is noteworthy that disruption of the gene encoding the membrane-bound Cdr1 efflux pump that has a role in phospholipid translocation and other facets of membrane physiology abrogates the capacity of the 1,4-BZDs to potentiate FLC. Another observation connecting the potentiators and lipid homeostasis is their perturbation of fungal morphology. The 1,4-BZDs alone dramatically alter the ability of *C. albicans* to filament and their combination with FLC significantly enhances the size and distribution of lipid droplets. The former observation is particularly significant because it has implications for filamentation-associated pathogenicity. While additional mechanistic studies are needed to define the precise mechanism of potentiation, the observations reported herein strongly link 1,4-BZD potentiation of FLC with lipid homeostasis in *C. albicans*.

Irrespective of the mechanistic ambiguities, our findings have implications for the treatment of infections by azole-resistant *Candida* species. We were excited to find that combinations of 1,4-BZDs and FLC reduced *C. albicans* fungal burdens and improved host survival in the *G. mellonella* model of systemic fungal infection. Some of the efficacy in this *in vivo* infection model can be ascribed to the fact that the 1,4-BZDs alone reduce filamentation, which is a key contributor to infection progression and lethality in this animal model (103). It remains to be established what filamentation pathways are modulated by 1,4-BZDs, either directly or indirectly. *In vivo* studies using models of mammalian fungal infection will be required to determine the efficacy of these molecules against *Candida* infection either alone or in combination with azoles.

Our observations support additional optimization of the different 1,4-BZD chemotypes as well as the molecular characterization and identification of the genes and pathways that underlie the effects of the 1,4-BZDs. Understanding the processes targeted by these compounds on their own and in combination with azoles will provide new opportunities for the development of antifungal therapies.

## MATERIALS AND METHODS

### Synthesis of molecules

The 1,4-BZDs were prepared via a diversity-oriented synthetic scheme (Fig. S1). The scheme begins with substituted acetanilides (1): purchased or synthesized from aniline (104). We accessed the respective 2-aminoketones (2a) from the acetanilides through a combination C-H activation and subsequent acid-mediated removal of the acetyl-group (46, 105). The 2-aminoketones were converted to the substituted 5-phenyl-1,3,4,5-tetrahydro-2H-benzo[e][1,4]diazepin-2-one (3a, secondary amine) or the N-butyl-2-(2-oxo-5-phenyl-2,3,4,5-tetrahydro-1H-benzo[e][1,4]diazepin-1-yl)acetamide (4a) through a one-pot tandem reaction (involving amide formation for 3a and Ugi reaction for 4a), followed by intramolecular imine formation and reduction (106, 107). The secondary amine was further converted to the final products through acylation (acylation product 5a), or reductive amination (reductive amination products 6a and 7a) (108–111). The detailed procedures, characterization of 1,4-BZDs, and drugs used in this study are provided in the Supplemental Material.

### Growth of fungal isolates

Strains used in this study are listed in Table S1. Fungal isolates were cultured overnight in 3 mL of liquid YPD (2% glucose, 2% bacto-peptone, 1% yeast extract, 2% dextrose) at 30°C with shaking (200 rpm). Cell densities were measured using optical densities of culture dilutions (OD_600_) in sterile water using a SPECTRAmax 250 Microplate Spectrophotometer. For growth curves, *C. albicans* cells were seeded in 96-well plates at a concentration of 2 x 10^5^ cells/mL and grown in YPD at 30°C or 37°C with orbital shaking for 24 h. Cell densities were measured every 15 min (OD_600_) using a Biotek Epoch 2 plate reader and doubling times were calculated using the exponential growth phase of each curve.

### Disk diffusion assays

For disk diffusion assays, *Candida* cultures were diluted to 1 x 10^6^ cells/mL in sterile water and 100 μL of each diluted culture was spread on a YPD plate using glass beads. The plates were allowed to dry and a disk containing 25 μg FLC (Liofilchem, 9166) was placed in the center of each plate. The plates were incubated for 48 h at 30°C, at which point they were photographed using a digital camera. To examine cidality, the FLC disk was replaced with a disk containing 8 mg of glucose and plates were grown for an additional 48 h. Disk images were analyzed using the R package *diskImageR* (35). The radius of inhibition (RAD) measures the distance (mm) from the center of the disk to the point where 20% or 50% reduction in growth occurs (RAD_20_ and RAD_50_, respectively), quantifying the extent of the zone of inhibition around the FLC disk. The fraction of growth (FoG_20_) calculates the area under the curve from the center of the disk to the 20% cutoff of inhibition. At least three independent disk assays were performed for each experiment.

### MIC testing

MIC testing was performed using broth microdilution assays in 96-well plates. FLC (Sigma Aldrich, PHR1160) was serially diluted (1 in 2) in liquid YPD to final concentrations of 0 – 128 μg/mL. Each well contained 125 μL liquid YPD volume with *C. albicans* cells at a concentration of 2 x 10^5^ cells/mL. Plates were incubated at 30°C with shaking (200 rpm) and cell densities (OD_600_) were measured at 0, 24 and 48 h using a SPECTRAmax 250 Microplate or a BioTek Epoch2 Spectrophotometer. Plates were examined for contamination after each incubation period. MIC_50_/MIC_90_ values were determined after 24 h growth by identifying the drug concentration leading to less than or equal to 50%/90% growth relative to growth without drug (0 μg/mL FLC). Supra-MIC values, or SMG, were determined after 48 h growth by measuring the average growth levels above the MIC_50_ levels relative to growth without drug. At least 3 independent cell culture replicates were assayed for each experiment.

For MIC assays using 1,4-BZDs in combination with other drugs, the respective drugs were serially diluted (1 in 2) in liquid YPD with the highest concentrations for myriocin (Sigma Aldrich, M1177) at 128 μg/mL, for terbinafine (Sigma Aldrich, PHR1298) at 32 μg/mL, and for fenpropimorph (Sigma Aldrich, 36772) at 32 μg/mL. The 1,4-BZDs were maintained at 100 μM in these experiments.

For experiments examining the potentiation effect of 1,4-BZDs in RPMI, *C. albicans* cells were grown in RPMI-MOPS medium buffered to pH 7 (1,04% RPMI with glutamine and phenol red, without sodium bicarbonate (Sigma Aldrich, R6504), 3,45% MOPS, 2% D(+) Glucose, pH adjusted with NaOH) and the MIC assays were carried out as those described in YPD with the only modification that growth was carried out at 30°C, 35°C and 37°C.

### Checkerboard assays and FICI determination

For checkerboard assays 96 well plates were set up with drug combination containing FLC (0-128 µg/mL, X axis) and 1,4-BZDs (0-100 µM, Y axis) and serially diluted. Each well was seeded with *C. albicans* cells at a concentration of 2 x 10^5^ cells/mL in 125 μL liquid YPD volume. Plates were incubated at 30°C with shaking (200 rpm) and cell densities (OD_600_) were measured at 0, 24 and 48 h using a SPECTRAmax 250 Microplate Spectrophotometer. Plates were examined for contamination after each incubation period. FLC MIC_50_/MIC_90_ values for each row were determined after 24 h growth by identifying the drug concentration leading to less than or equal to 50%/90% growth relative to growth without drug (0 μg/mL FLC). Fractional inhibitory concentration indices (FICI_50_ and FICI_90_) were calculated as previously described (25) using FLC MIC_50_ and MIC_90_ levels, respectively (described in Tables S2-4). 3-4 independent assays were performed for each strain and for each 1,4-BZD.

### Time-kill assays

*C. albicans* cultures were grown overnight in liquid YPD at 30°C with shaking (200 rpm). Next, 5 x 10^4^ cells from each isolate were cultured at 30°C (200 rpm) for 48 h in a volume of 200 µl of YPD + DMSO, YPD + FLC (128 μg/mL), or YPD + FLC (128 μg/mL) + 1,4-BZD (100 μM) in 96 well plates. At different time points during growth (0, 4, 8, 12, 24, 36 and 48 h), 10 μL of each culture was diluted into PBS and the cells were plated on YPD agar and incubated at 30°C for CFU determination. The CFUs were counted at 48 h and for each condition the number of colonies was used to determine survival proportions (%) relative to the 0 h time point. Experiments were performed with three biological replicates.

### Cytotoxicity assays

The LDH release assay is a colorimetric assay, based on the cytosolic lactate dehydrogenase released into culture supernatant from damaged peripheral blood mononuclear cells (PBMCs). The assay was performed using the Cytotoxicity Detection Kit (Roche, 11644793001) following the instruction of the manufacturer. Briefly, blood samples from healthy donors were obtained from the Établissement Français du Sang (EFS Paris, France) with written informed consent following the guidelines provided by the Ethics Committee of Institut Pasteur (convention 12/EFS/023). PBMCs were isolated from blood samples by Ficoll (Eurobio, France) density-gradient separation. Isolated PBMCs were washed, resuspended in RPMI medium (at a concentration of 2 x 10^6^/mL) and seeded into 96-well tissue culture plates (100 μL/well). Each well was added with a concentration range of compounds diluted in DMSO (20 μL/well) and the total volume was made up 200 μL/well by adding 80 μL/well of RPMI. DMSO alone was used as the negative control, while Triton-X100 (1%) served as the positive control. After incubating the culture plate at 37°C in a 5% CO_2_ incubator for 1 h, the plate was subjected to centrifugation (1200 rpm, 10 min) and 100 μL of supernatants from each well were transferred to a 96-well flat bottom microplate. Freshly prepared cytotoxicity reaction mixture was then added to each well and incubated for 30 min at ambient temperature. The color developed was measured at 492 nm using a plate reader (Tecan Infinite 200PRO, Tecan). The optical densities read represented the amount of lactate dehydrogenase released, and thereby a proxy for cytotoxicity.

For hemolytic assays, human blood samples (2 mL) were mixed with 8 mL of PBS (pH 7.4) and subjected to centrifugation at 1000 rpm for 3 min. The RBC pellets obtained were washed 4 times with PBS (8 mL, each time). From the final suspension, 200 μL of RBCs (from the bottom of the tube) were treated to 9.8 mL PBS, from which 100 μL/well were transferred to 96-well cell culture plates. The RBC suspensions were treated with different concentrations of compounds (in a volume of 20 μL/well) diluted in DMSO and incubated at 37°C with 5% CO_2_ for 30 min. After incubation, the culture plates were centrifuged at 1200 rpm (10 min), the supernatant from each well was transferred to a 96-well flat bottom microplate and the color was measured at 541 nm using a plate reader (Tecan Infinite 200PRO, Tecan). DMSO treated RBCs served as the control, and Triton-X100 (1%) treated RBCs was used as a positive control.

### Flow cytometry

*C. albicans* cultures were grown overnight in liquid YPD at 30°C with shaking (200 rpm). Cultures were diluted 1:100 in 2-3 mL YPD medium and compounds were added to a final concentration of 10 µg/mL for FLC, 10 µg/mL for Beauvericin, 1 µg/mL for Rhodamine 6G, 100 µM for 1,4-BZDs, and 100 µM for sortin2. Cells were incubated at 30°C with shaking and harvested after 4 h, washed twice with sterile PBS and diluted to a density of ∼10^6^ cells/mL in PBS. Fluorescence was measured in 200 µL aliquots of cell suspension and data were collected from 100,000 cells per sample using a Guava easyCyte 3^rd^ generation or a Miltenyi MACSQuant Analyzer 10 Flow Cytometer. Cell populations were gated by SSC/FSC to eliminate small debris particles. Experiments were performed with at least three biological and two technical replicates. Analyses were performed using FlowJo 10.8. Differences between conditions/treatments were evaluated using *t* tests, * *P* < 0.05.

### *In vitro* filamentation assays

For filamentation, *C. albicans* cells were grown overnight in liquid YPD at 30°C with shaking, washed with sterile PBS and resuspended in fresh YPD at a concentration of 10^5^ cells/mL. 1 mL of cell suspension was added to 24-well plates and the plates were incubated at 37°C without shaking for 2 h in the presence of DMSO (no drug), FLC alone (10 µg/mL), 1,4-BZDs (100 µM) or a combination of FLC (10 µg/mL) and 1,4-BZDs (100 µM). Wells were examined for distribution of cells on the surface of the wells and images of approximately 500-1000 cells were captured at a 40X magnification using a Zeiss AxioVision Rel. 4.8 microscope. Image analysis was performed using ImageJ to quantify the proportion of yeast, pseudohyphal, and hyphal cells present in each condition. Assays were performed with 4 biological and 2 technical replicates, differences between treatments were evaluated using *t* tests, * *P* < 0.05.

### Fluorescent imaging of lipid droplets

*C. albicans* cells were grown overnight in liquid YPD at 30°C with shaking and resuspended in fresh YPD at a concentration of 10^5^ cells/mL in combination with different treatments: DMSO (no drug), FLC (128 µg/mL), combinations of FLC (128 µg/mL) and 1,4-BZDs (100 µM), or Myriocin (32 μg/mL) as positive control. Cells were then incubated for 4 h at 30°C with shaking and washed twice with sterile PBS to remove media. To stain lipids, the cells were incubated with 1 μg/mL Bodipy 493/503 (D3922, Invitrogen) for 10 min at 30°C with shaking and washed twice with PBS before imaging. Samples were imaged at a 60X magnification using a GFP filter on a Zeiss AxioVision Rel. 4.8 microscope with >200 cells observed per sample. All assays were performed with biological triplicates.

### G. mellonella infections

*G. mellonella* larvae were obtained from Vanderhorst Wholesale (Saint Marys, OH, USA) or from La Ferme aux Coleos (Cherbourg-en-Cotentin, France) and the infection protocol was adapted from (14). Larvae were kept at 15°C and used within 1 week of delivery. Larvae without signs of melanization and an average weight of 0.25 g were selected in groups of 12 larvae. *C. albicans* inoculums were prepared from cultures grown overnight at 30°C in liquid YPD and washed twice with sterile PBS. Cell densities (OD_600_) were measured using a SPECTRAmax 250 Microplate Spectrophotometer and adjusted to 3 × 10^7^ cells/mL in sterile PBS. Each larva was first injected with 3 × 10^5^ *C. albicans* cells via the last left pro-leg using a 10 µL glass syringe and a 26S gauge needle (Hamilton, 80300). A second 10 µL injection with either FLC (20 µg/mL), 1,4-BZDs (50 µM), FLC (20 µg/mL) + 1,4-BZDs (50 µM), or sterile vehicle (DMSO) was done via the last right pro-leg 1.5–2 h post *C. albicans* infection. The inoculum size was confirmed by plating on YPD for colony forming units (CFUs). Infected and controls groups of larvae were maintained at 37°C for 10 days and survival was checked daily. Larvae were recorded as dead if no movement was observed upon contact. Virulence assays were performed in 2-4 independent experiments (*n* = 24-48 total larvae per condition). Control groups of larvae included mock PBS injections (no *C. albicans*) followed by treatment with DMSO, FLC (20 µg/mL) or 1,4-BZDs (50 µM). No significant killing of larvae was observed in either of the control conditions. Statistical differences between larval groups were tested using log-rank (Mantel-Cox) and Gehan-Breslow-Wilcoxon tests (* *P* < 0.05, ** *P* < 0.01, *** *P* < 0.001, **** *P* < 0.0001) and are fully described in Table S5.

An additional set of virulence assays was performed to quantify fungal burdens in *G. mellonella* larvae. After 4 days of infection (or at the time of death for larvae that had died before this time point), the larvae were homogenized in sterile PBS + antibiotics (PBS 1X, 500 µg/mL penicillin, 500 µg/mL ampicillin, 250 µg/mL streptomycin, 250 µg/mL kanamycin, 125 µg/mL chloramphenicol, and 125 µg/mL doxycycline), and dilutions of homogenates were plated onto YPD to assess the abundance of fungal cells from each treatment group by CFU determination. For imaging *C. albicans* cells in the host, each slide was prepared with 9 μL of *Galleria* homogenate to which 1 μL calcofluor white stain was added (calcofluor white M2R, 10 mg/L, Evans blue, 5 mg/L, Sigma Aldrich). Images were taken with a Leica M80 Stereomicroscope at a 60X magnification. Images were captured with a DMC 2900 camera, using the Leica Application Suite (LAS) imaging software. The number of cells in each morphology was determined from 3 larvae per group, by examining between 240 to 618 fungal cells per larva (average 411 cells/larva). For FLC-treated groups, the number of cells observed under the microscope was not sufficient for statistical analyses (<50 cells per larva).

### Statistical analyses

Unless otherwise specified, statistical analyses were carried out in GraphPad Prism 9 using two-tailed t-tests (* *P* < 0.05).

## ACKNOWLEDGEMENTS

We thank Richard Bennett (Brown University), Dominique Sanglard (Lausanne University Hospital), Brian LeBlanc (Brown University), Joseph Bliss (Brown University) and Arnaldo Colombo (Federal University of Sao Paolo) for the generous gifts of strains. This work was supported by Brown University UTRA and SPRINT awards (TPM), NIH NIAID R21AI139592 (IVE), Île de France Dim One Health (IVE), and Institut Pasteur (IVE). IVE is a CIFAR Azrieli Global Scholar in the CIFAR Program Fungal Kingdom: Threats & Opportunities. JKS and PEA acknowledge generous funding from the Chan Zuckerberg Biohub and Brown University.

## Supplemental Figure Legends

**Figure S1.** Synthetic scheme for accessing the different compounds tested. **a.** Pd(OAc)_2_ 0.1eq., TBHP 4eq., TFA 0.3eq, substituted benzaldehyde or substituted benzyl alcohol 2eq., 40°C, 16 h; **b.** excess Conc. HCl, EtOH, 75°C, 16 h; **c.** Glycine N-carboxyanhydride (NCA) 1.3eq., TFA 2eq., Toluene (0.1 M), 65°C, 30 min; **d.** Et_3_N 2eq., 85°C, 30 min; **e.** NaBH_3_CN, AcOH, MeOH, room temperature, 16 h; **f.** n-butylisocyanide 1.1eq., formaldehyde 1eq., Boc-Glycine 1eq., MeOH, 24 h; **g.** TFA 2eq., Toluene, 70°C; **h.** NaBH_3_CN, AcOH, MeOH, 16 h; **i.** R_1_COCl, NaHCO_3_, CHCl_3_, 0°C to room temperature, 12 h; **j** or **k** R_1_CHO, NaBH_3_CN, AcOH, MeOH, room temperature, 16 h.

**Figure S2.** Impact of potentiator/FLC drug combinations on FLC susceptibility, tolerance and viability of *C. albicans*. Compounds were screened using FLC disk diffusion assays (25 µg/disk) on YPD plates containing 100 µM of potentiator or the corresponding volume of DMSO (vehicle control). Heatmaps show FLC susceptibility (RAD_50_ and RAD_20_, A) and tolerance levels (FoG_20_, B) following 48 h of growth. After 48 h, FLC disks were replaced with glucose disks (gluc, 8 mg/disk), grown for an additional 48 h, and susceptibility and tolerance were quantified again. Heatmaps show average values for 3 or more biological replicates, arrows indicate the top seven 1,4-BZDs. (C) Representative images for FLC (at 48 h) and glucose (at 96 h) disk diffusion assays for strains SC5314, SP-945 and P60002 for DMSO, sortin2 and the top seven 1,4-BZDs (used at 100 µM).

**Figure S3.** Impact of 1,4-BZD/FLC drug combinations on FLC susceptibility and tolerance when *C. albicans* cultures were grown in RPMI at different temperatures (30, 35 and 37°C). YPD data at 30°C is included for reference. Compounds were screened using liquid MIC assays containing 100 µM of 1,4-BZDs and a gradient of FLC concentrations (0 to 128 µg/mL). Heatmaps show average values for changes in FLC susceptibility (MIC_50_, A) and tolerance (SMG, B) relative to levels observed with DMSO alone. Experiments were performed with three biological replicates.

**Figure S4.** 3D plots showing inhibition of fungal growth due to varying 1,4-BZDs (0 to 100 µM) and FLC (0 to 128 µg/mL) concentrations in checkerboard assays. Sortin2 was included for reference. Plots show an average of 2-4 biological replicates.

**Figure S5.** Summary of checkerboard assays showing the impact of drug combinations of FLC and potentiators on FLC susceptibility (MIC_50_) for *C. albicans* isolates SC5314, SP-945 and P60002. (B) FICI_50_ scores for combinations of potentiators and FLC based on 50% growth inhibition. Left-side table indicates the type of FLC-potentiator interaction based on the FICI_50_ score. FICI_50_ scores show an average of 2-4 biological replicates.

**Figure S6.** Changes in antifungal susceptibility (RAD_50_ and RAD_20_, A) and tolerance (FoG_20_, B) using combinations of a subset of 1,4-BZDs and non-azole antifungals. Compounds were screened using disk diffusion assays on YPD plates containing 1,4-BZDs (100 µM) or DMSO (no drug) and using disks of flucytosine (AFY, 1 µg), griseofulvin (AGF, 10 µg), amphotericin B (AMB, 10 µg), caspofungin (CAS, 5 µg), and nystatin (NY, 100 IU). Heatmaps indicate average values from 3 biological replicates.

**Figure S7.** Susceptibility profiles (MIC_50_ in µg/mL, relative values for SMG) of (A) *C. albicans* isolates SC5314, SP-945 and P60002 to FLC, terbinafine (TERB), fenpropimorph (FEN), myriocin (MYO). (A-C) FLC susceptibility profiles (MIC_50_ in µg/mL, relative values for SMG) of different gene deletion mutants and their WT controls. Symbols next to gene names denote the corresponding parental strains. Heatmaps show average values from 3-4 biological replicates. (D) Impact of 1,4-BZDs on intracellular rhodamine 6G accumulation for *cdr1*, *cdr2*, and *cdr1 cdr2* mutants and corresponding WT parental strain. Rhodamine 6G fluorescence intensity levels were measured after 4 h incubation with rhodamine 6G (1 µg/mL) and treatment with DMSO (no drug). Heatmaps show average values from 3 or more biological replicates.

**Figure S8.** *C. albicans* filamentation in *G. mellonella* hosts. Representative images (A) and quantification (B) of *C. albicans* SC5314 and P60002 cell morphologies observed in homogenates of *G. mellonella* larvae treated with either PBS (no drug), PA158 or PA162 (4 days post infection). Cells were stained with calcofluor white, and images were taken at 60X magnification, scale bar is 20 µm. Histogram shows average fractions of cell populations from 3 larvae per group, with >240 fungal cells examined for each larva. Asterisks indicate significant differences relative to PBS treated larvae, *t* test, * *P* < 0.05.

## Supplemental Tables

**Table S1.** Yeast strains used in this study. For the different species, the isolates are listed in the order in which they appear in Fig. 6.

**Table S2.** Checkerboard assays and FICI calculations for strain SC5314.

**Table S3.** Checkerboard assays and FICI calculations for strain SP-945.

**Table S4.** Checkerboard assays and FICI calculations for strain P60002.

**Table S5.** Statistical comparisons for *Galleria mellonella* experiments.

