## Supplemental Material for "Small Molecules Restore Azole Activity Against Drug-Tolerant and Drug-Resistant *Candida* Isolates"

[5]Current address: Cell Biology Program, Sloan Kettering Institute, New York, NY

† equal contributions

*1.7 Synthesis of substituted 5 substituted phenyl-4-(thiazole-2-carbonyl)-1,3,4,5-tetrahydro-2H-benzo[e][1,4]diazepin-2-one 3 (tertiary amide, 5) from secondary amine via acylation.* ..*8*

*1.8 Reductive aminations of the secondary amines (3, 4) to afford the tertiary amines (6, 7)*..........................................................................................................................*9*

**2.0 CHARACTERIZATION DATA OF FINAL ANALOGS: $^{1}$H NMR, $^{13}$C NMR, HRMS 10**

**REFERENCES.........................................................................................................63**

### 1.0 Synthetic protocols

Analytical thin-layer chromatography (TLC) was performed with silica gel GF254 plates. Column chromatography was performed with silica gel (300–400 mesh) eluting with solvent mixtures (ethyl acetate (EtOAc)/hexane, acetone/hexane). Nuclear magnetic resonance (NMR) spectroscopy, high resolution mass spectrometry (HRMS, with relevant soft ionization techniques) were used to characterize the synthesized analogs and intermediates. $^{1}$H NMR and $^{13}$C NMR spectra were recorded at 400 MHz and 600 MHz (Bruker) with $CDCl_3$ as solvent. Chemical shifts are reported in ppm with relevant solvent calibration as standards. FLC-Cy5, was synthesized and characterized using previously described protocols (1). Dicyclomine (Sigma Aldrich), sertraline (Sigma Aldrich), proadifen (Sigma Aldrich), sortin2 (Sigma Aldrich), chloroquine (Sigma Aldrich), antifungal disks (Liofilchem), fluconazole (Sigma Aldrich), rhodamine 6G (Sigma Aldrich), beauvericin (Sigma Aldrich) were all used as received.

#### 1.1 Synthetic scheme for accessing the 1,4-BZD analogs (Figure S1)

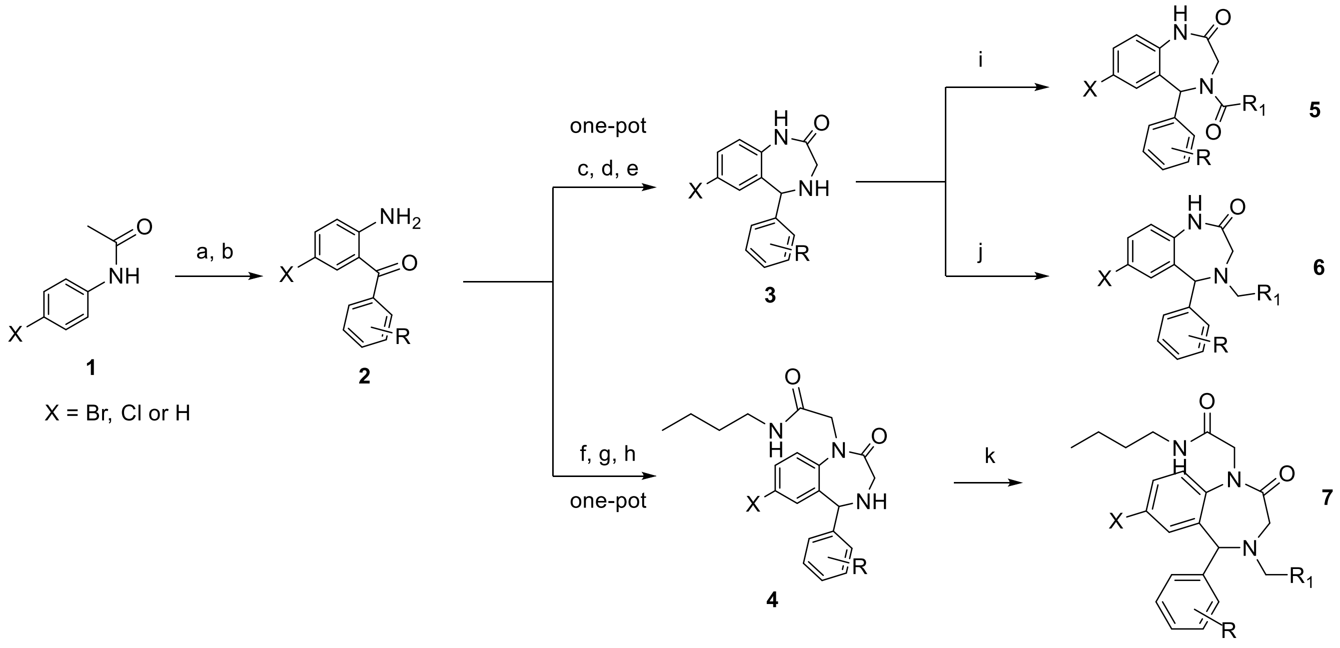

The scheme begins with substituted acetanilides **1**, purchased or synthesized from aniline (2). The respective 2-aminoketones **2** were accessed from the acetanilides through C-H activations and subsequent acid-mediated deacetylation (3, 4). Further conversion of the 2-aminoketones to the substituted 5-phenyl-1,3,4,5-tetrahydro-2H-benzo[e][1,4]diazepin-2-one **3** (secondary amine) or the N-butyl-2-(2-oxo-5-phenyl-2,3,4,5-tetrahydro-1H-benzo[e][1,4]diazepin-1-yl)acetamide **4** via a one-pot tandem reaction involving three transformations. These transformations were amide formation or Ugi reactions, followed by intramolecular imine formation and imine reduction to obtain the secondary amines (5, 6). The conversion of the secondary amines via acylation (product **5**), or reductive amination (reductive products **6** and **7**) afforded the different analogs (7-10).

#### 1.2 Synthesis of acetanilides (1) from anilines (0)

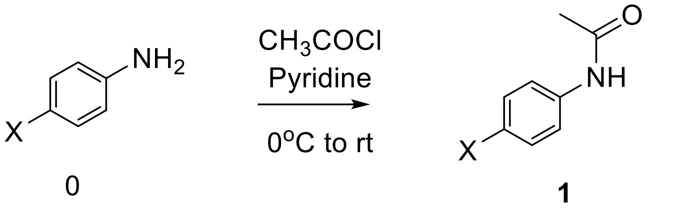

Acetyl chloride (1.1eq) was added to a solution of substituted aniline **0** (4 mM), Pyridine (1.1eq) in dry $CH_2Cl_2$ (20 mL) at 0 °C, the mixture was allowed to cool to ambient temperature and stirred until the consumption of starting material was observed (monitored by TLC). After completion of reaction, the crude reaction mixture was sequentially washed and extracted with 2 N solution of HCl, brine and subsequently dried over $Na_2SO_4$. After filtration, the crude solution (organic layer) was concentrated in vacuo and purified by flash column chromatography (using EtOAc/hexane) to afford substituted acetanilide **1**. The yield was between 70-95% (2).

#### 1.3 Synthesis of N-acetylaminoketones (1a) from acetanilides (1)

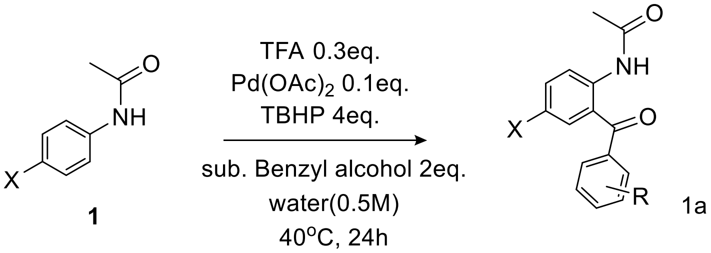

Acetanilide (1eq), $H_2O$ (0.5 M solution of acetanilide **1**), TFA (0.3eq), substituted benzylic alcohols **or** substituted benzaldehydes (2eq) and TBHP (4eq. using a 70 % solution in $H_2O$) were added to a 50 mL $Pd(OAc)_2$ charged round bottom flask (0.1eq). A rubber septum was used to seal the reaction vessel and the mixture was stirred at 40 $^oC$. After completion (monitored disappearance of acetanilide by TLC, usually ~16 h for most reactions), the reaction mixture was dissolved in EtOAc and the organic layer was extracted with water, brine and dried over sodium sulfate. The organic layer was concentrated in vacuo. The concentrated crude was purified by flash column chromatography using EtOAc and hexane solvent mixtures. The yield of products **1a** was 25-85 %. Higher yields were observed for Deshaloacetanilides.

#### 1.4 Synthesis of aminoketones (18) from N-acetyl-aminoketones (17)

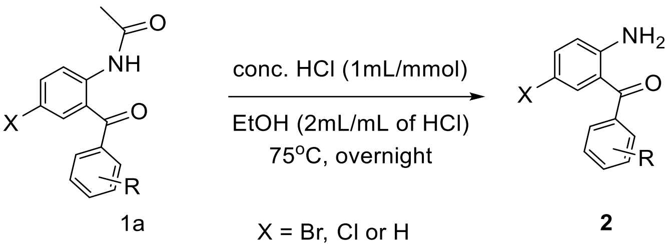

Concentrated HCl (1 mL of HCl per mM of **1a**) was added to a solution of the keto acetamide **1a** in ethanol (EtOH, 2 mL/mM) and the mixture was stirred at 75 °C for 16 h, the reaction crude was cooled to room temperature and the solution was subsequently brought to a pH of 8 using saturated $NaHCO_3$ solution at room temperature. EtOAc was added to the aqueous solution followed by extraction of the biphasic mixture. The organic layer was washed with water, brine and dried over $Na_2SO_4$ to yield a crude reaction solution. The solution was concentrated in vacuo and purified using flash column chromatography (5-8% EtOAc/hexane as solvent mixture) to afford mostly yellowish solids, aminoketones **2** (the yield was 75-93%) (3).

#### 1.5 Synthesis of substituted 5-phenyl-1,3,4,5-tetrahydro-2H-benzo[e][1,4]diazepin-2-one 3 (secondary amine, 3) from amino ketones via tandem cyclization, imine formation and reduction

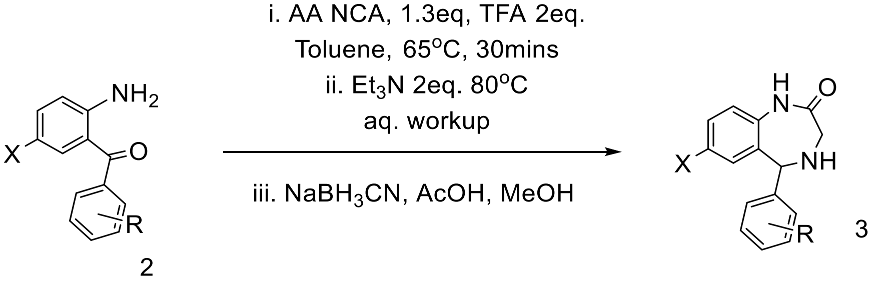

The synthesis was adapted from (6). Aminoketone (**2**) and toluene (0.2 M solution of aminoketone) were loaded onto a 20 mL round bottom flask. 2,2,2-Trifloroacetic acid (2eq.) was added and solid glycine N-carboxyanhydride (Gly NCA, 1.3eq.) was added subsequently. The reaction was heated to 60 $^oC$ for 1 h. We ensured the reaction was properly vented to allow the release of $CO_2$ during this period, usually done through a nitrogen/argon inlet and an air needle running through the septum. Next, neat triethylamine ($Et_3N$, 2eq.) was added to the reaction mixture, the reaction was heated to 80 $^oC$, allowed to stir for another 1 h and monitored by TLC/MS for complete conversion to the benzodiazepine-imine product. On completion, the reaction was cooled to room temperature, followed by evaporation of the toluene and addition of EtOAc to the reaction mixture, this crude solution of EtOAc was extracted with water, brine and dried over sodium sulphate. The crude solution was then concentrated in vacuo. Methanol (MeOH, 0.2 M) was then added to the crude mixture and excess acetic acid (3eq.), the mixture was cooled to 0 $^oC$. Sodium cyanoborohydride (4eq.) was then added to the cooled mixture, allowed to warm slowly to room temperature and stirred overnight (16 h) at room temperature. After completion of reaction, sodium bicarbonate was added to the crude mixture to bring the pH to 8, and then extracted with EtOAc. The organic layer was further extracted with water, brine, dried over sodium sulphate and filtered. The filtrate was subsequently concentrated in vacuo and purified by flash column chromatography (using a 20% and 50% solution of EtOAc/hexane) to yield the product **3** (the yield was 40-82%).

**1.6 Synthesis of substituted N-butyl-2-(2-oxo-5-phenyl-2,3,4,5-tetrahydro-1H-benzo[e][1,4]diazepin-1-yl)acetamide 4 from amino ketones via tandem C-H activation and reduction**

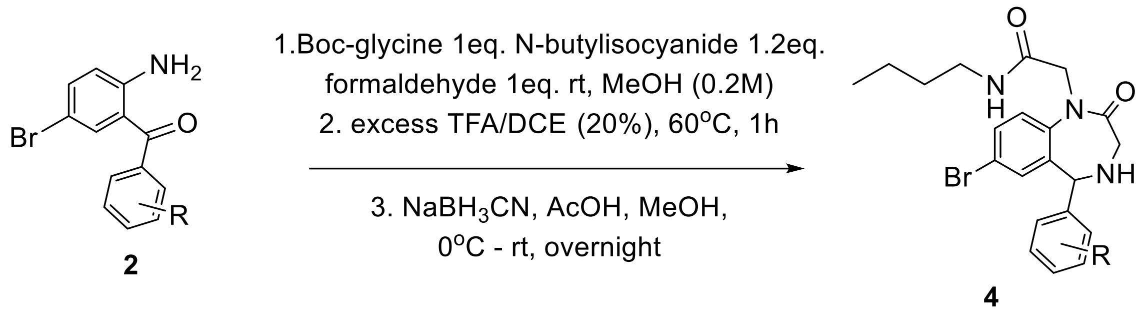

one-pot

The synthesis was adapted from (5). At ambient temperature, n-butylisocyanide (1.3eq.), boc-glycine (1eq.), and formaldehyde (1eq. using a formalin solution) were added to a solution of aminoketone (1eq) **2** in MeOH (0.3 M) and stirred at room temperature for 2-3 days. After completion, DCM (3 mL) and PS-p-TsOH (polystyrene supported toluene sulfonic acid, 2eq.) were added to the reaction mixture and stirred for 60 minutes. After stirring, the solid residue was filtered off and washed with MeOH, EtOAc and DCM (twice with each solvent). The solution was concentrated in vacuo and a 20% solution of TFA in DCE (3 mL) was added to the concentrated solution and stirred at 60 $^{o}$C for 1 h. After 1 h, the obtained solution was cooled to room temperature, treated with sodium bicarbonate, and extracted with EtOAc. The organic layer was washed with brine, dried over sodium sulphate, and concentrated in vacuo to afford the crude mixture. Next, MeOH (0.2 M) and excess acetic acid (3eq.) were added to the crude mixture which was cooled to 0 $^{o}$C. sodium cyanoborohydride (4eq.) was added to the cooled solution and the mixture was allowed to warm slowly to room temperature and stirred for 16 h at room temperature. After this reaction, $NaHCO_3$ was added to the crude mixture, to bring the pH to 8, and the aqueous solution was then extracted with EtOAc, washed with water, brine, dried over sodium sulphate and filtered. The filtrate was concentrated in vacuo and purified by flash column chromatography (20-70% gradient EtOAc/hexane) to yield the product (yield 64%).

**1.7 Synthesis of substituted 5 substituted phenyl-4-(thiazole-2-carbonyl)-1,3,4,5-tetrahydro-2H-benzo[e][1,4]diazepin-2-one 3 (tertiary amide, 5) from secondary amine via acylation.**

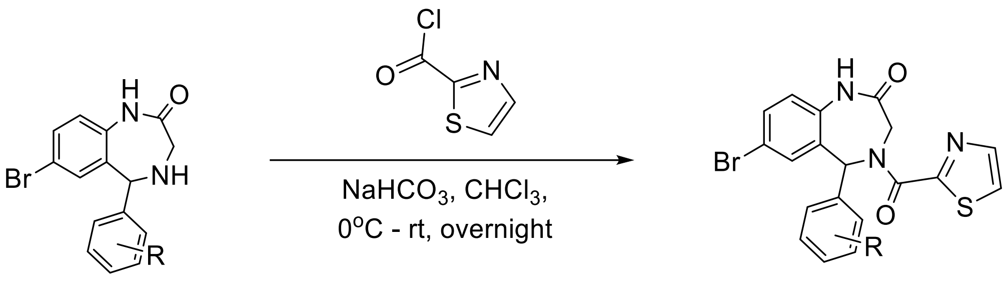

**3** 5

0.33 mM of the secondary amine **3** were added to a round bottom flask with a Teflon coated stir bar. $CHCl_3$ (0.2 M) was added, and the mix was brought to 0 $^{o}$C under a nitrogen atmosphere. Next, 3eq. of sodium bicarbonate was added to the solution followed by the slow addition of 1.3eq. of 2-thionyl chloride and allowed to stir for 16 h. On completion of the reaction, the residue was filtered off through celite and the filtrate was concentrated in vacuo to afford the crude product. The crude product was purified by flash column chromatography (usually with 15-20% EtOAc/hexane solvent mixture). The yield was 84%.

**1.8 Reductive aminations of the secondary amines (3, 4) to afford the tertiary amines (6, 7)**

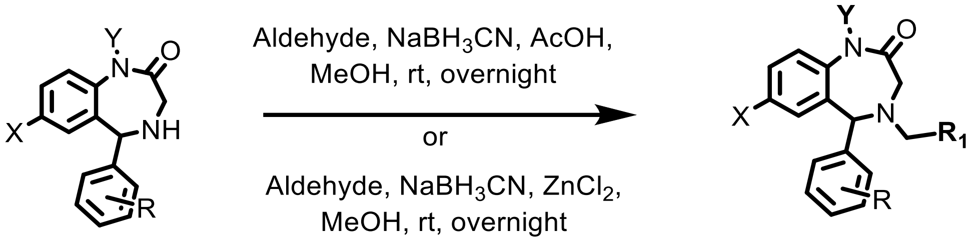

**3** Y = H
**4** Y = $CH_2CONHCH_2CH_2CH_2CH_3$

**6** Y = H
**7** Y = $CH_2CONHCH_2CH_2CH_2CH_3$

The reductive aminations were performed using either of the two different protocols:

1. 1.5eq of the suitable aldehyde was added to a 0.33 mM solution of the secondary amine **3** or **4** in MeOH (0.3 M) and stirred at 0 °C. 2eq. of $ZnCl_2$ (using a 1 M $ZnCl_2$ solution in MeOH or THF) was added to this mixture, followed by the addition of 3eq. of sodium cyanoborohydride in small batches. After the complete addition of sodium cyanoborohydride, the mixture was allowed to warm to room temperature and stirred overnight. On completion of the reaction, sodium bicarbonate was added to the reaction mixture and the mixture was extracted with EtOAc, washed with water, brine and dried over sodium sulphate. The crude reaction solution was concentrated in vacuo and purified by Preparative TLC plates (on silica), using a 40% acetone/hexane solvent mixture in most cases. The isolated product yield was 43-82% (8, 9).
2. 1.5eq of the aldehyde was added to a 0.33 mM solution of the secondary amine **4** in MeOH (0.3 M) and stirred at 0 °C. 3eq. of AcOH and 3eq. of sodium cyanoborohydride were added to the mixture, the mixture was allowed to warm to room temperature and stirred for 16 h. After reaction completion (monitored by TLC and MS), sodium bicarbonate was added to the reaction mixture and the mixture was extracted with EtOAc, washed with water, brine and dried over sodium sulphate. The organic layer was concentrated in vacuo and purified by preparative

TLC plates (on silica), using a 40% acetone/hexane solvent mixture. The isolated product yield was 72% (11).

### 2.0 Characterization data of final analogs: $^{1}$H NMR, $^{13}$C NMR, HRMS

FLC-Cy5 was synthesized and characterized using previously described synthetic protocols (1).

**PA104**

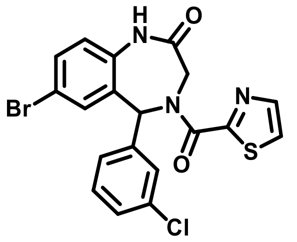

**(7-bromo-5-(3-chlorophenyl)-4-(thiazole-2-carbonyl)-1,3,4,5-tetrahydro-2H-benzo[e][1,4]diazepin-2-one)**

**$^{1}$H NMR** (600 MHz, Chloroform-*d*) δ 8.95 (d, *J* = 91.5 Hz, 1H), 7.84 (dd, *J* = 6.1, 3.2 Hz, 1H), 7.57 – 7.43 (m, 2H), 7.39 – 7.32 (m, 1H), 7.16 (t, *J* = 4.6 Hz, 2H), 7.07 – 6.93 (m, 1H), 6.93 – 6.81 (m, 2H), 5.59 – 4.12 (5.59 (d, *J* = 17.0 Hz), 4.64 (d, *J* = 17.1 Hz), 4.51(d, *J* = 16.5 Hz), 4.12 (d, *J* = 16.5 Hz), 2H).

**$^{13}$C NMR** (151 MHz, CDCl3) δ 170.33, 170.23, 163.89, 163.68, 160.13, 160.05, 143.90, 143.83, 140.68, 139.08, 135.24, 135.20, 135.14, 135.04, 134.25, 133.80, 132.80, 132.66, 130.37, 130.24, 129.93, 129.52, 128.60, 128.51, 127.67, 127.55, 125.77, 125.67, 125.42, 123.57, 123.14, 117.66, 117.31, 61.66, 61.02, 60.42, 49.53, 48.10. Peaks are duplicated because of the existence of rotamers.

**HRMS (ESI):** m/z calculated for $C_{19}H_{13}BrClN_3O_2S$ $[M+H]^+$: 460.9600, found: 461.9668.

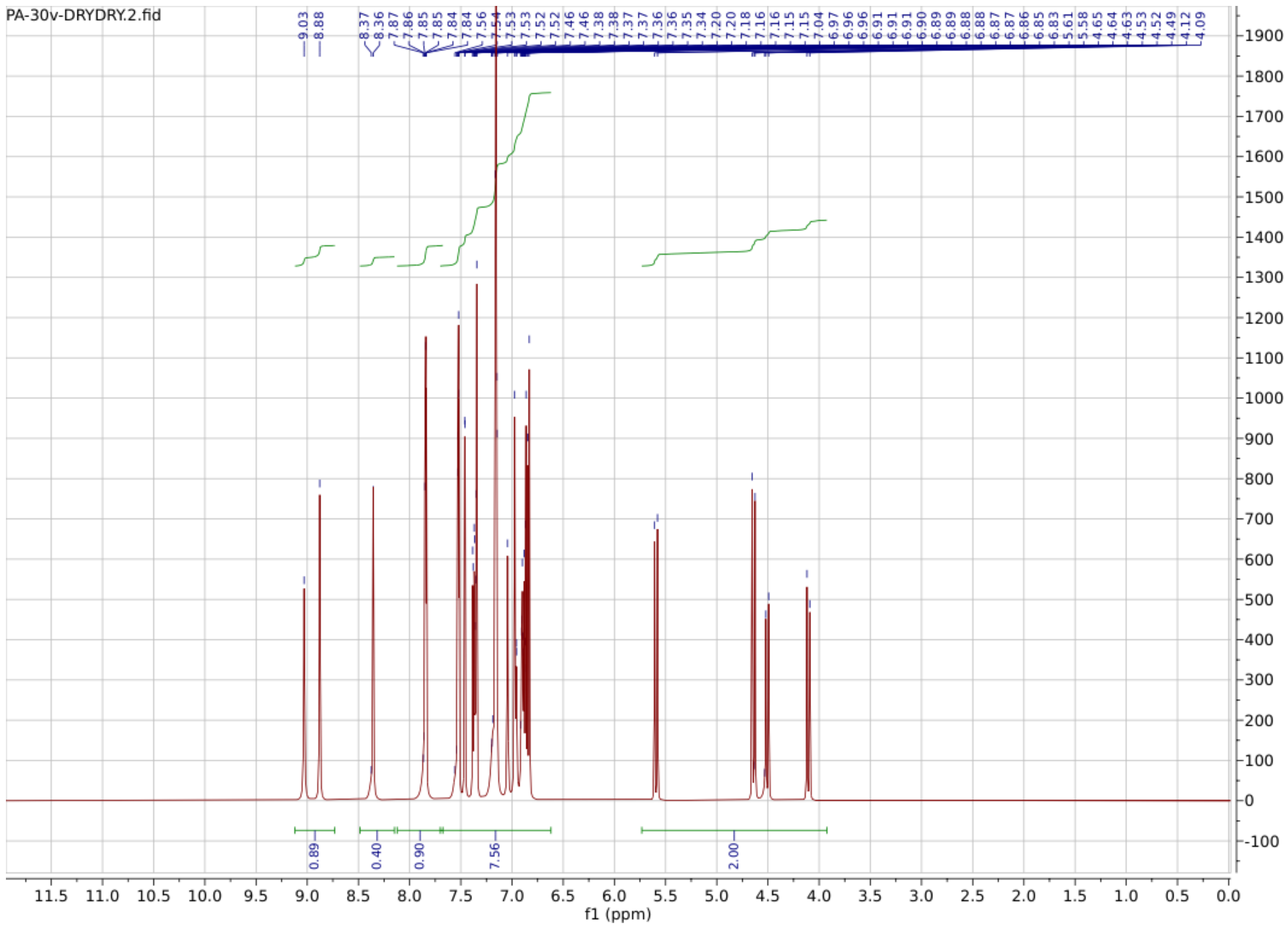
PA-30v-DRYDRY.2.fid
9.03
8.88
8.37
8.36
7.86
7.85
7.85
7.84
7.84
7.56
7.54
7.53
7.53
7.52
7.52
7.46
7.38
7.38
7.37
7.37
7.36
7.36
7.35
7.34
7.20
7.18
7.16
7.16
7.15
7.15
7.04
6.97
6.96
6.96
6.91
6.91
6.91
6.90
6.89
6.89
6.88
6.88
6.87
6.86
6.85
6.83
5.61
5.58
4.65
4.64
4.63
4.53
4.52
4.49
4.12
4.09
0.89
0.40
0.90
7.56
2.00
f1 (ppm)

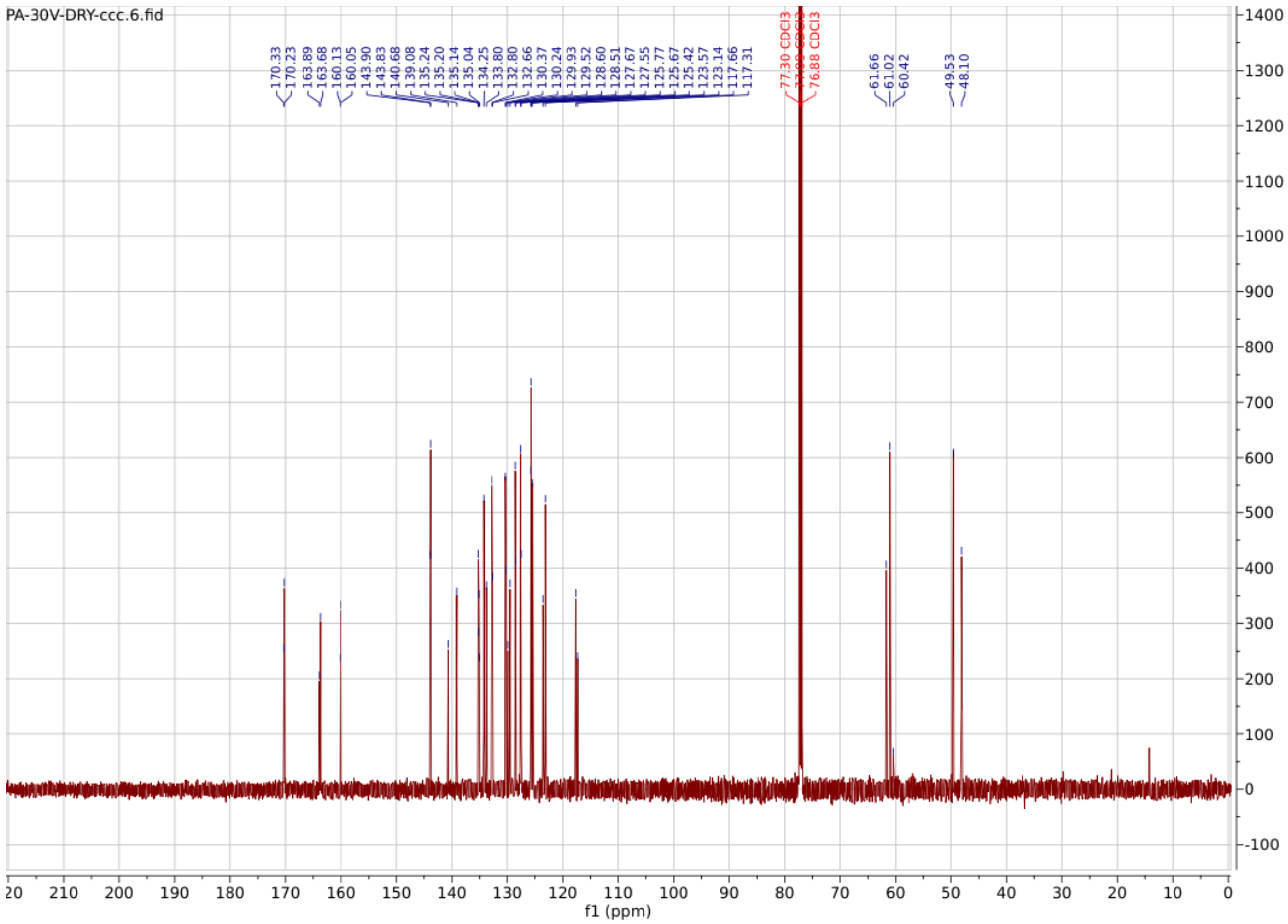

**PA84**

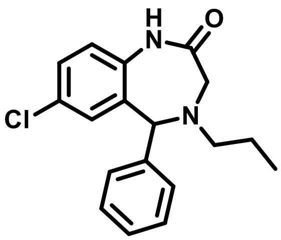

**7-chloro-5-phenyl-4-propyl-1,3,4,5-tetrahydro-2H-benzo[e][1,4]diazepin-2-one**

$^{1}$**H NMR** (600 MHz, $CDCl_3$) δ 8.15 (s, 1H), 7.42 – 7.17 (m, 7H), 6.91 (d, *J* = 8.5 Hz, 1H), 6.86 (d, *J* = 2.4 Hz, 1H), 4.94 (s, 1H), 3.50 (d, *J* = 15.9 Hz, 1H), 3.38 (dd, *J* = 15.8, 1.4 Hz, 1H), 2.71 – 2.57 (m, 2H), 1.63 – 1.52 (m, 2H), 0.90 (t, *J* = 7.3 Hz, 3H). Peak 1.63-1.52 coincides with some water peaks. This should integrate to 2.

**$^{13}$C NMR** (151 MHz, $CDCl_3$) δ 173.09, 140.55, 135.65, 133.03, 131.08, 129.69, 128.68, 128.62, 128.44, 127.86, 121.61, 77.23, 77.02, 76.81, 68.55, 55.44, 52.69, 20.80, 11.54.

**HRMS (ESI):** m/z calculated for $C_{18}H_{19}ClN_2O$ $[M+H]^+$: 314.1186, found: 315.1253.

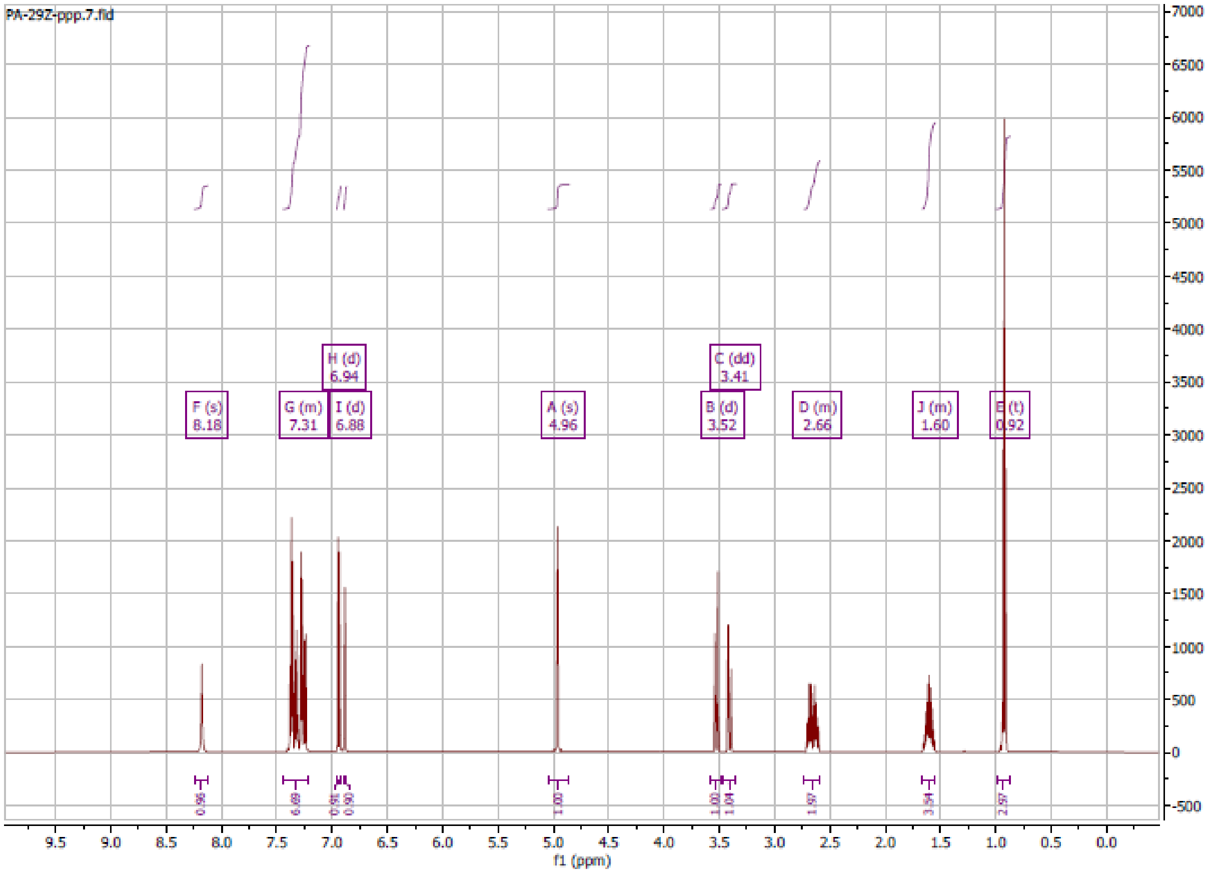

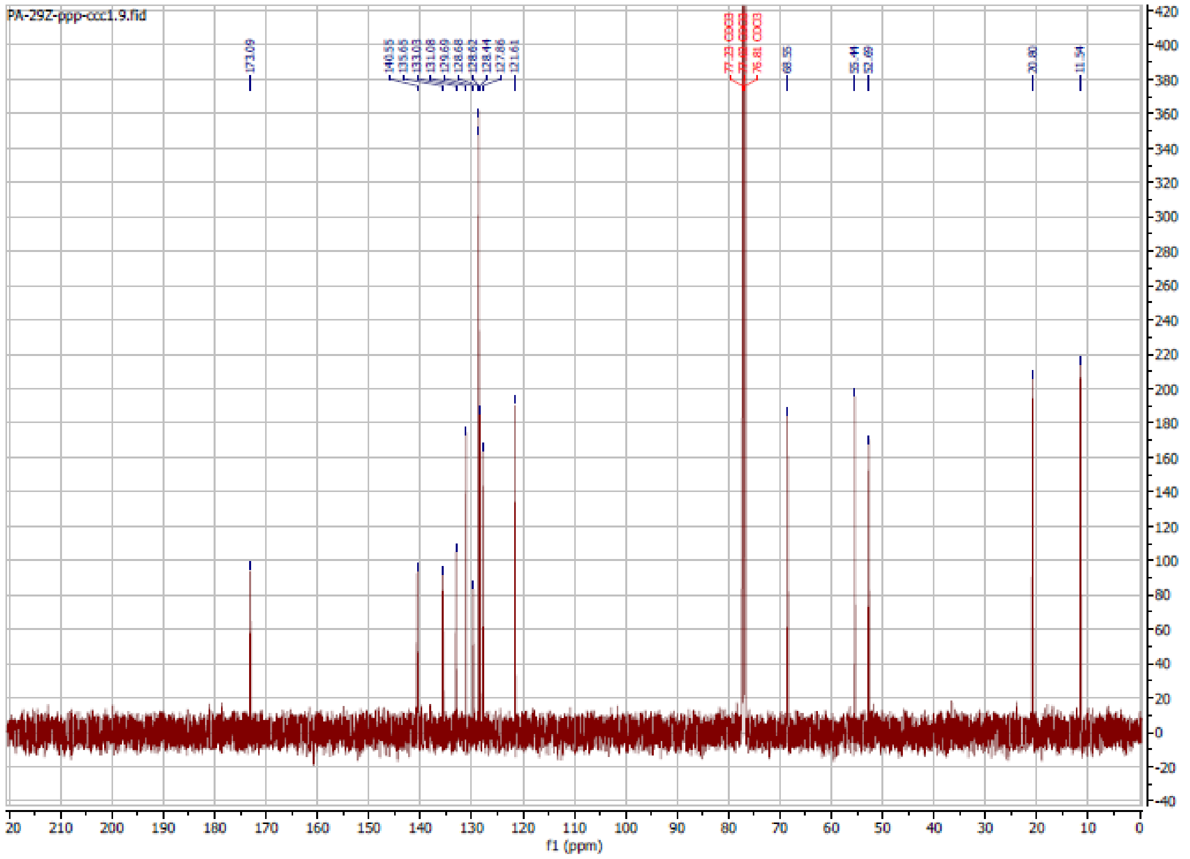

**PA85**

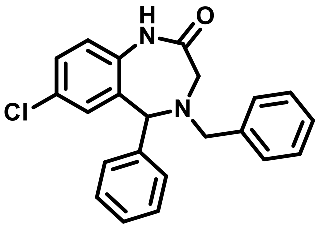

**4-benzyl-7-chloro-5-phenyl-1,3,4,5-tetrahydro-2H-benzo[e][1,4]diazepin-2-one**

**$^1$H NMR** (400 MHz, $CDCl_3$) δ 8.17 (s, 1H), 7.59 – 7.14 (m, 17H, includes chloroform), 7.06 – 6.94 (m, 1H), 6.84 (d, *J* = 2.4 Hz, 1H), 4.97 (s, 1H), 3.95 (d, *J* = 13.3 Hz, 1H), 3.79 (d, *J* = 13.3 Hz, 1H), 3.55 (d, *J* = 15.3 Hz, 1H), 3.34 (d, *J* = 15.3 Hz, 1H).

**$^{13}$C NMR** (151 MHz, $CDCl_3$) δ 172.19, 140.14, 137.92, 135.93, 133.07, 131.04, 130.03, 129.04, 128.84, 128.67, 128.58, 128.48, 128.08, 127.47, 121.76, 77.23, 77.02, 76.81, 67.93, 57.91, 52.35.

**HRMS (ESI):** m/z calculated for $C_{22}H_{19}ClN_2O$ $[M+H]^+$: 362.1186, found: 363.1258.

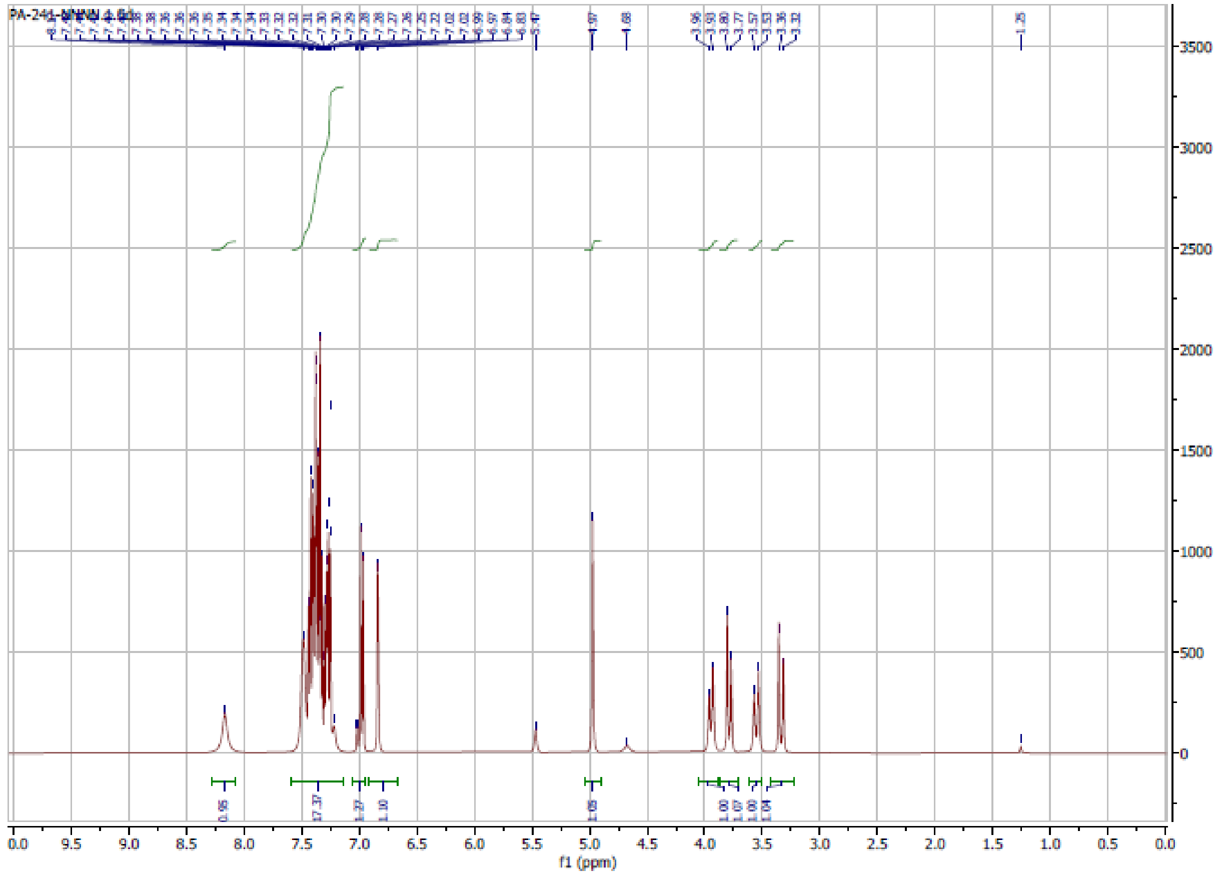

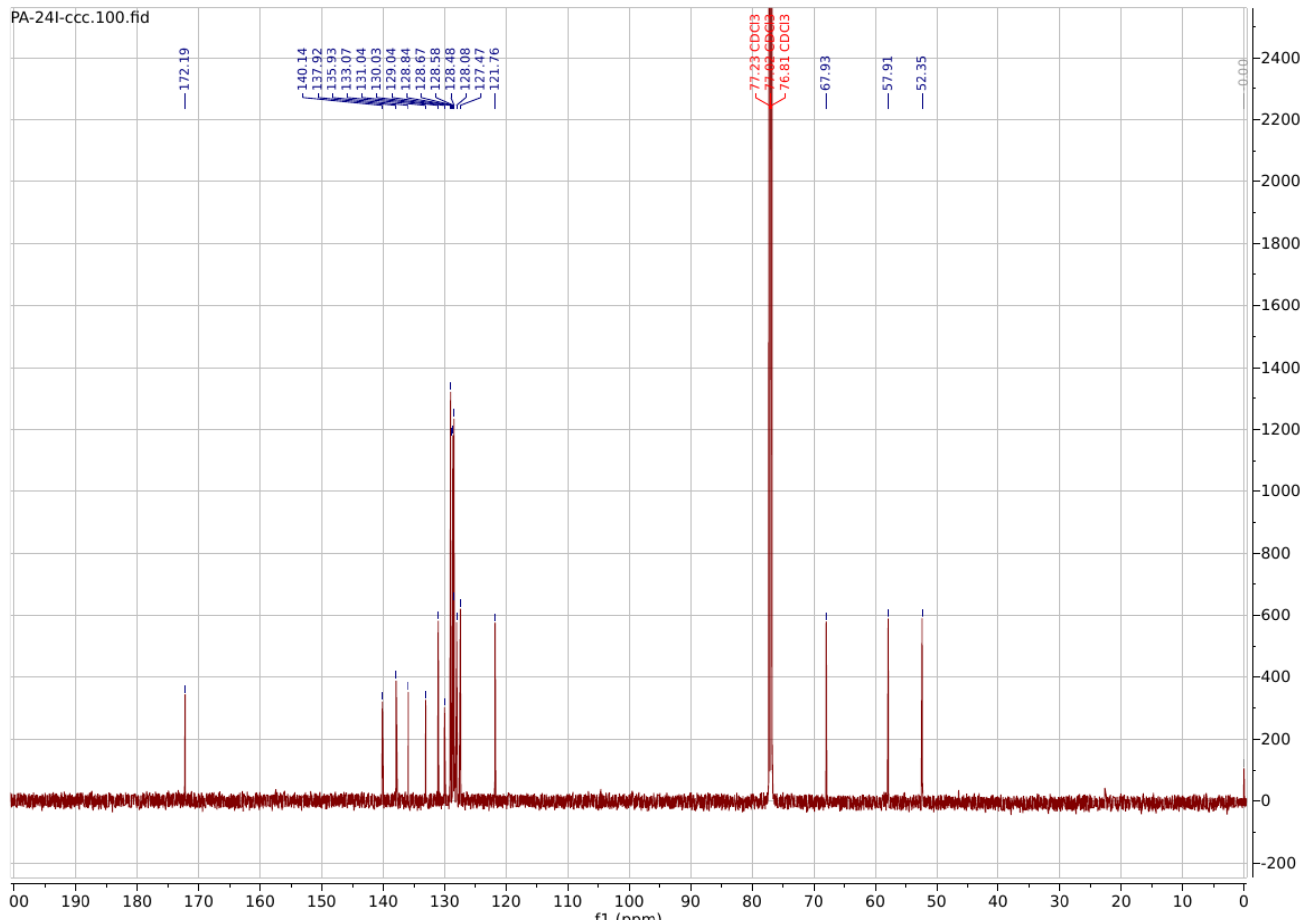

**PA86**

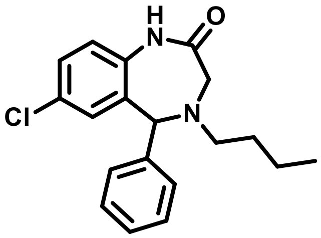

**4-butyl-7-chloro-5-phenyl-1,3,4,5-tetrahydro-2H-benzo[e][1,4]diazepin-2-one**

**$^{1}$H NMR** (600 MHz, Chloroform-*d*) δ 8.40 (s, 1H), 7.40 – 7.18 (m, 7H), 6.96 (d, *J* = 8.5 Hz, 1H), 6.89 (d, *J* = 2.4 Hz, 1H), 4.96 (s, 1H), 3.52 (d, *J* = 15.8 Hz, 1H), 3.42 (dd, *J* = 15.8, 1.4 Hz, 1H), 2.77 – 2.60 (m, 2H), 1.63 – 1.19 (m, 4H), 0.91 (t, *J* = 7.4 Hz, 3H).

**$^{13}$C NMR** (151 MHz, $CDCl_3$) δ 173.24, 140.59, 135.67, 132.97, 131.06, 129.64, 128.67, 128.61, 128.43, 127.83, 121.64, 77.23, 77.02, 76.80, 68.60, 53.33, 52.71, 29.80, 26.93, 20.17, 13.95.

**HRMS (ESI):** m/z calculated for $C_{19}H_{21}ClN_2O$ $[M+H]^+$: 328.1342, found: 329.1409.

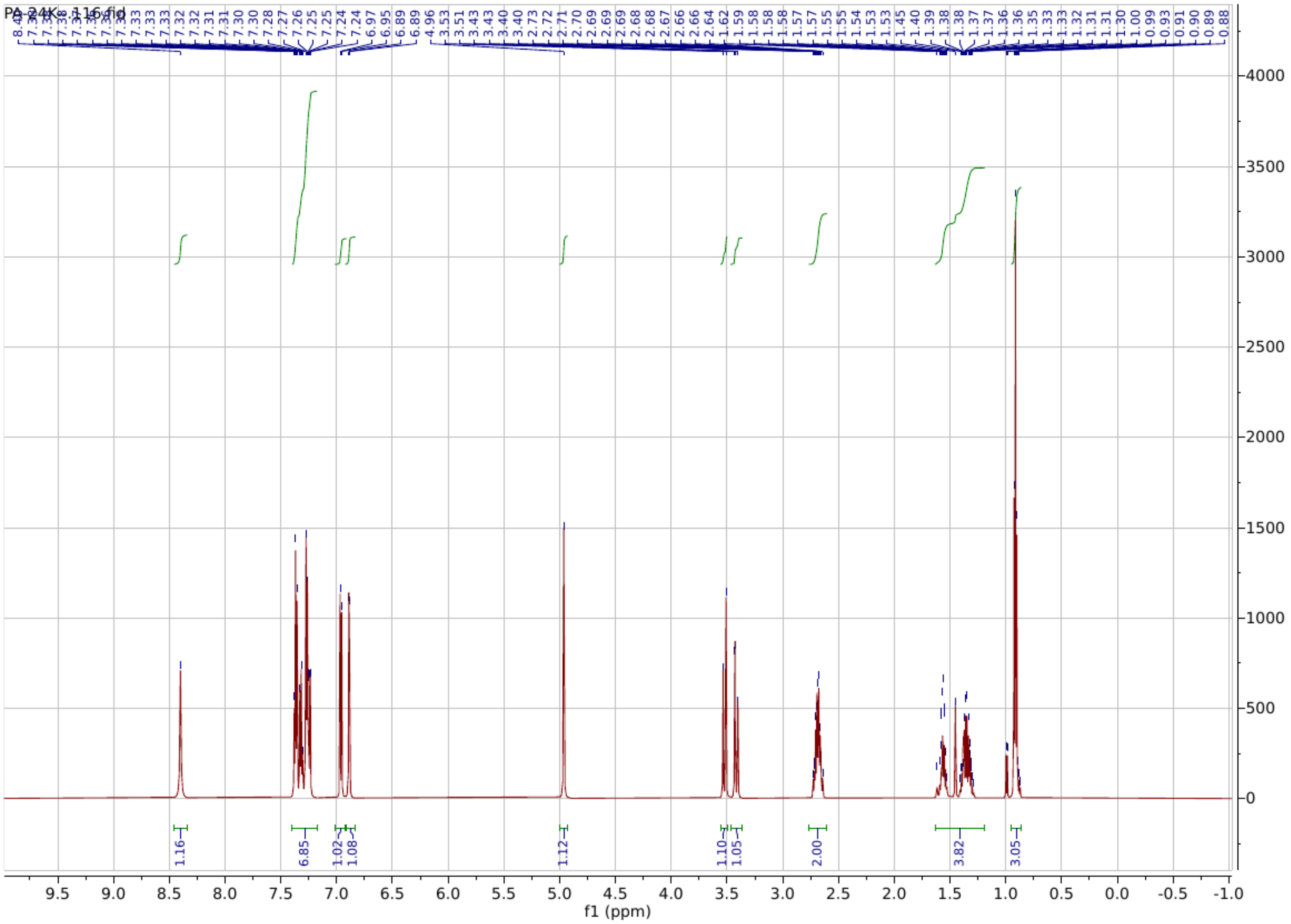

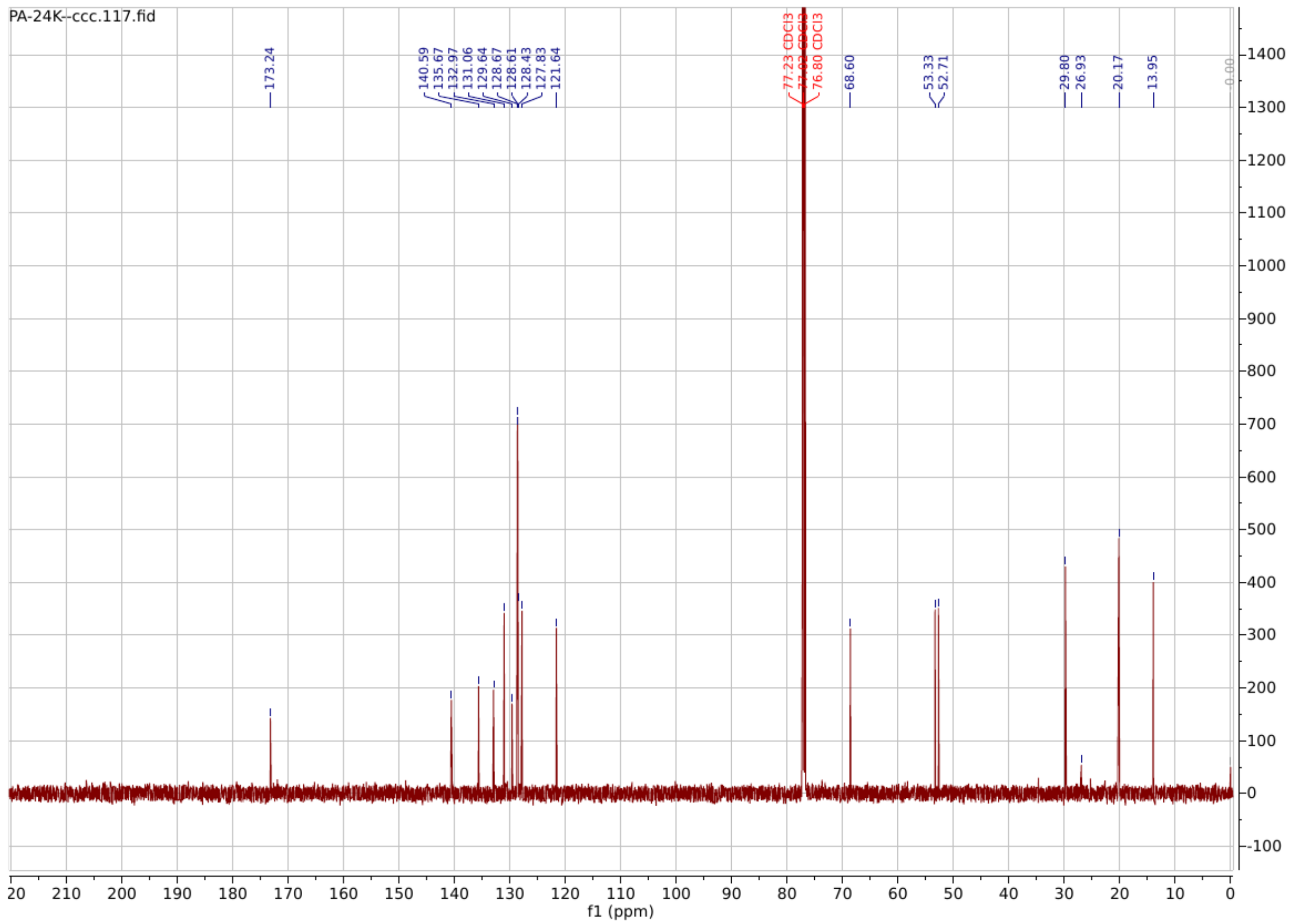

**PA87**

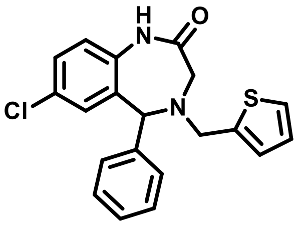

**7-chloro-5-phenyl-4-(thiophen-2-ylmethyl)-1,3,4,5-tetrahydro-2H-benzo[e][1,4]diazepin-2-one**

**$^1$H NMR** (600 MHz, Chloroform-*d*) δ 8.44 (s, 1H), 7.48 – 7.20 (m, 7H), 7.09 – 6.90 (m, 3H), 6.82 (d, *J* = 2.4 Hz, 1H), 5.01 (s, 1H), 4.08 – 3.98 (m, 2H), 3.54 (d, *J* = 15.5 Hz, 1H), 3.44 (dd, *J* = 15.5, 1.5 Hz, 1H).

**$^{13}$C NMR** (151 MHz, $CDCl_3$) δ 172.13, 141.80, 139.86, 135.81, 132.81, 131.08, 130.09, 128.87, 128.67, 128.58, 128.13, 126.58, 126.50, 125.48, 121.84, 77.24, 77.02, 76.81, 67.41, 52.64, 52.54.

**HRMS (ESI):** m/z calculated for $C_{20}H_{17}ClN_2OS$ $[M+H]^+$: 368.8790, found: 369.0819.

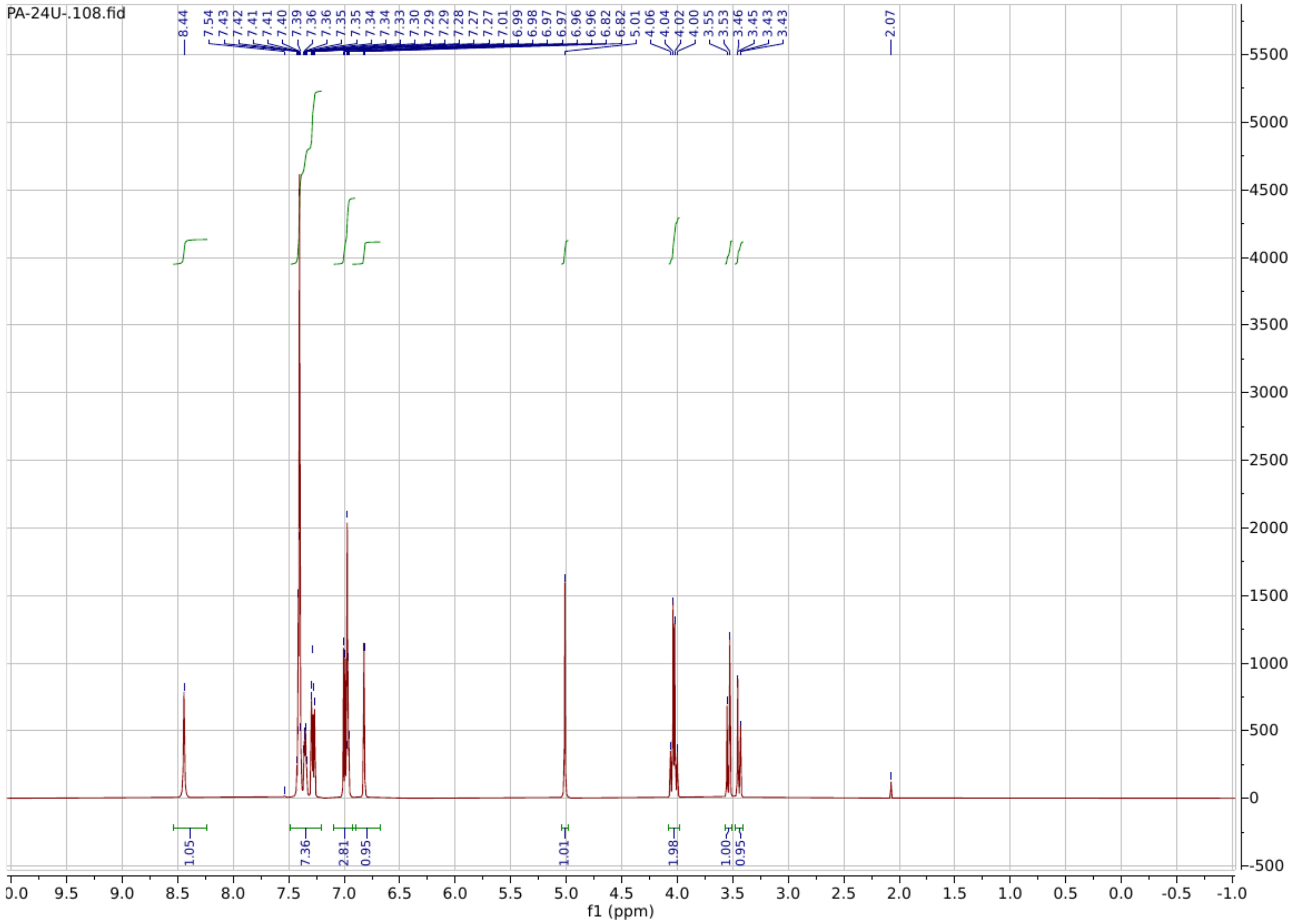

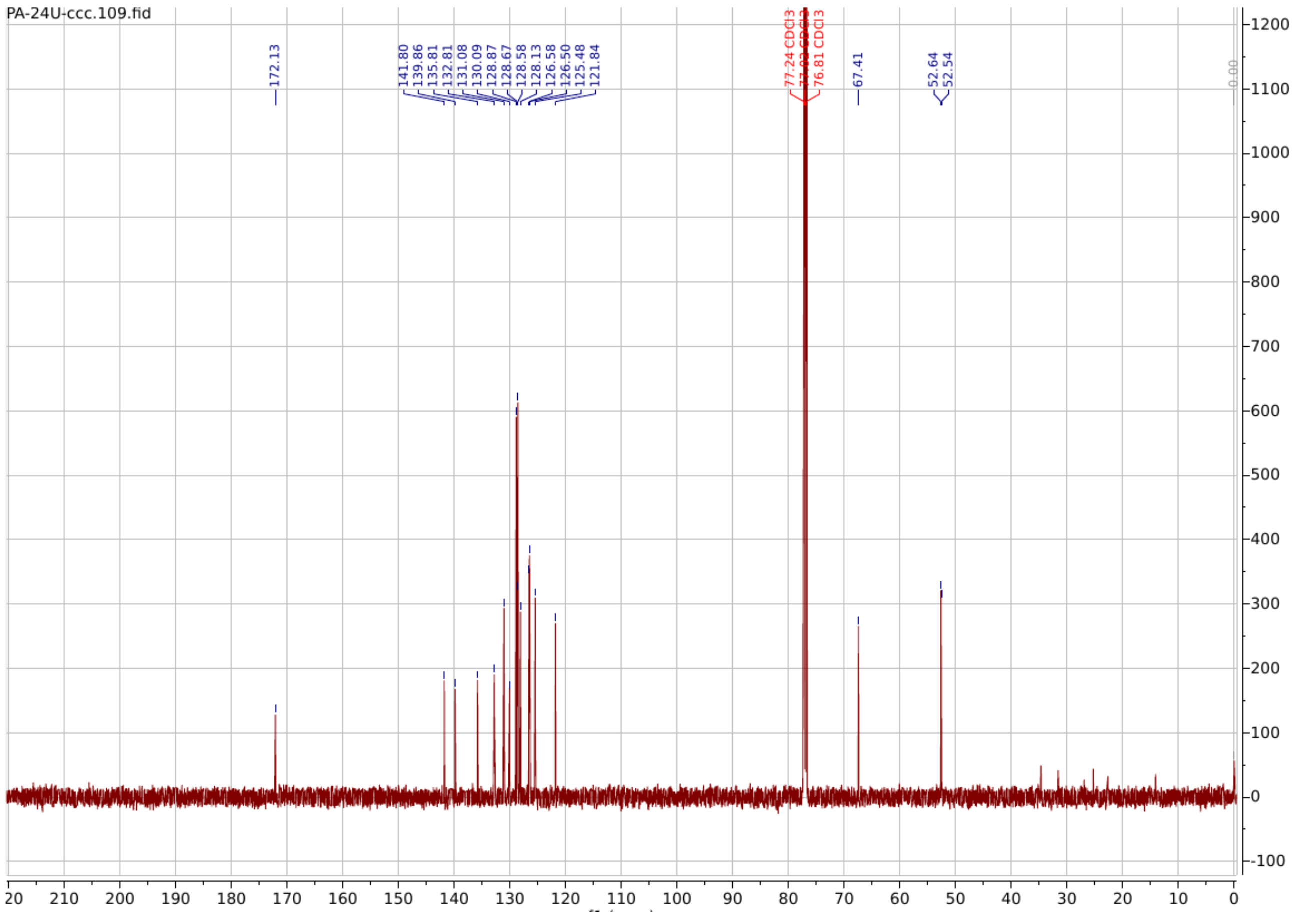

**PA88**

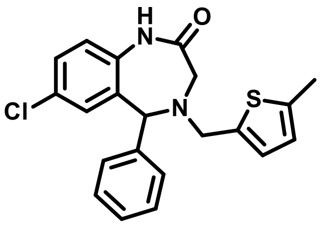

**7-chloro-4-((5-methylthiophen-2-yl)methyl)-5-phenyl-1,3,4,5-tetrahydro-2H-benzo[e][1,4]diazepin-2-one**

**$^{1}$H NMR** (400 MHz, $CDCl_3$) δ 8.06 (s, 1H), 7.60 – 7.26 (m, 8H), 7.04 – 6.76 (m, 3H), 6.60 (dt, *J* = 3.4, 1.1 Hz, 1H), 5.09 (s, 1H), 4.03 (q, *J* = 14.0 Hz, 2H), 3.77 – 3.48 (m, 2H), 2.46 (d, *J* = 1.2 Hz, 3H).

**$^{13}$C NMR** (151 MHz, $CDCl_3$) δ 171.67, 140.05, 139.93, 139.15, 135.69, 132.93, 131.16, 128.84, 128.62, 128.56, 128.09, 126.59, 124.51, 121.67, 77.22, 77.01, 76.80, 67.27, 52.87, 52.35, 15.49.

**HRMS (ESI):** m/z calculated for $C_{21}H_{19}ClN_2OS$ $[M+H]^+$: 382.0907, found: 383.0993.

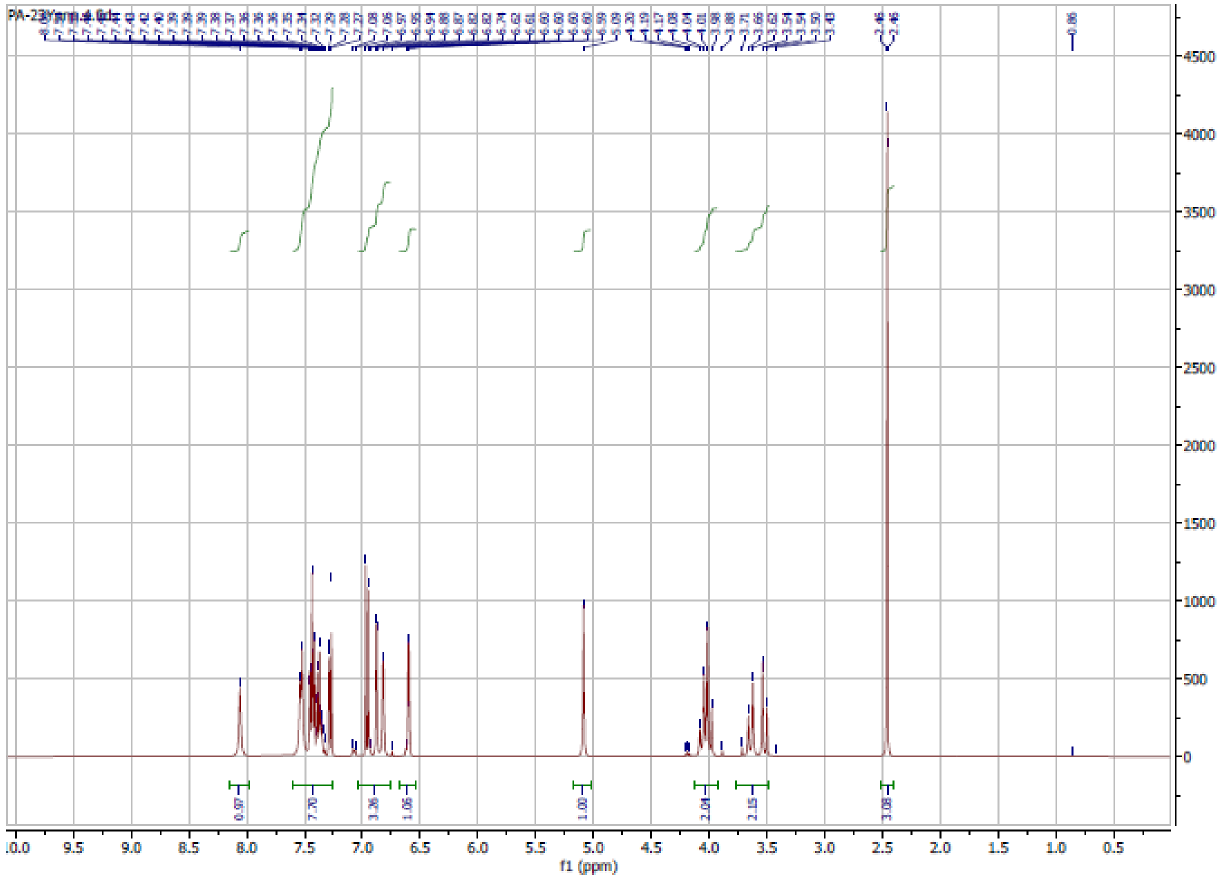

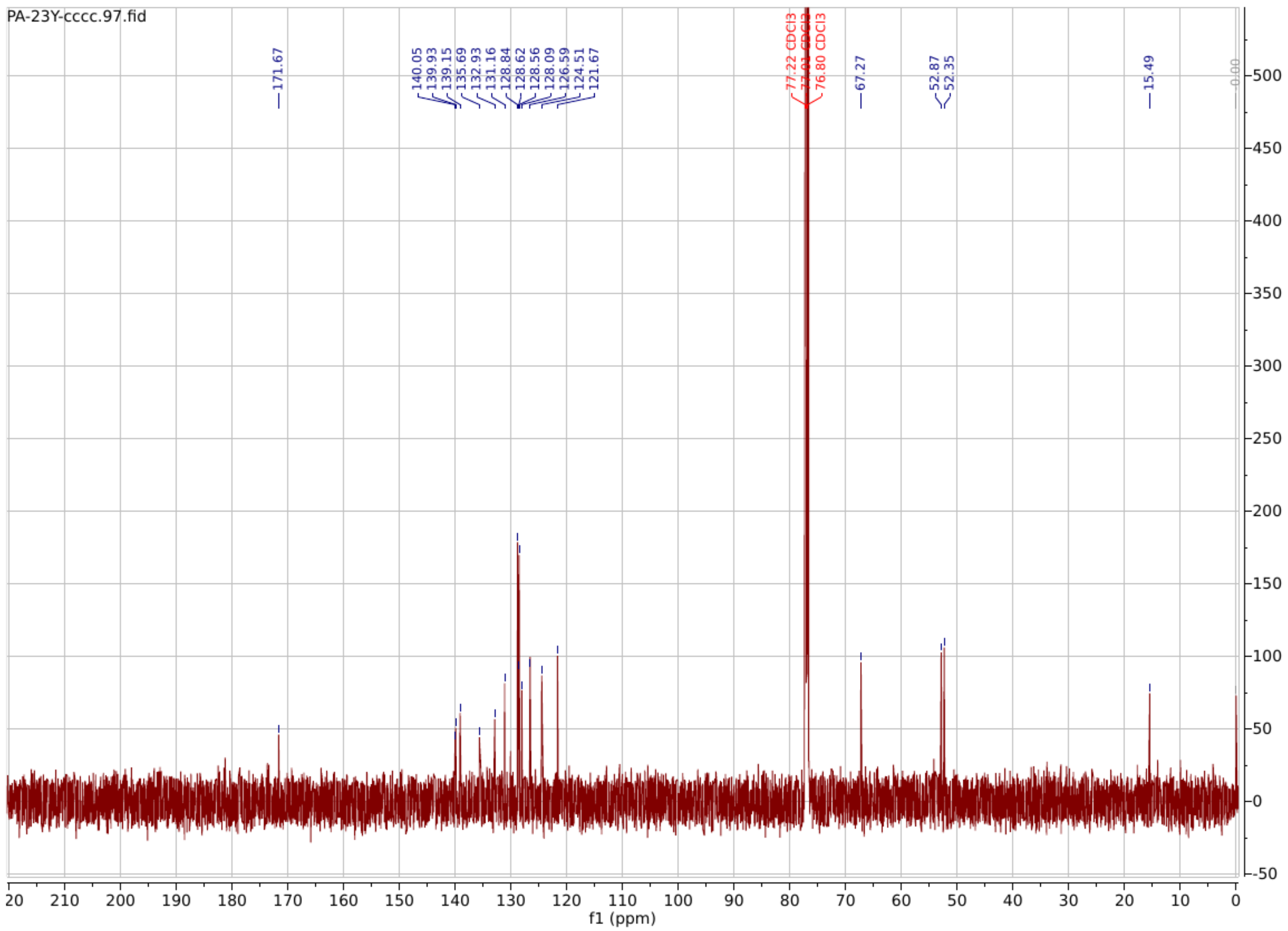

**PA89**

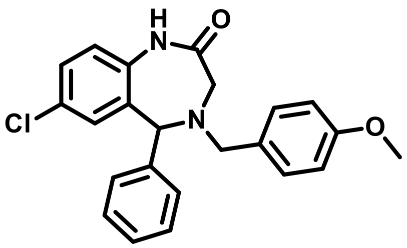

**7-chloro-4-(4-methoxybenzyl)-5-phenyl-1,3,4,5-tetrahydro-2H-benzo[e][1,4]diazepin-2-one**

**$^{1}$H NMR** (600 MHz, Chloroform-*d*) δ 8.39 (s, 1H), 7.45 – 7.18 (m, 8H), 6.99 (d, *J* = 8.5 Hz, 1H), 6.94 – 6.86 (m, 2H), 6.81 (d, *J* = 2.4 Hz, 1H), 4.93 (s, 1H), 3.82 (d, *J* = 16.8 Hz, 4H), 3.70 (d, *J* = 13.2 Hz, 1H), 3.48 (d, *J* = 15.5 Hz, 1H), 3.32 (dd, *J* = 15.4, 1.5 Hz, 1H).

**$^{13}$C NMR** (151 MHz, $CDCl_3$) δ 173.38, 160.23, 139.63, 136.63, 133.92, 131.55, 130.27, 129.96, 129.91, 128.81, 128.66, 128.53, 127.10, 121.77, 112.76, 66.75, 60.14, 55.69, 51.51.

**HRMS (ESI):** m/z calculated for $C_{23}H_{21}ClN_2O_2$ $[M+H]^+$: 392.1292, found: 393.1356.

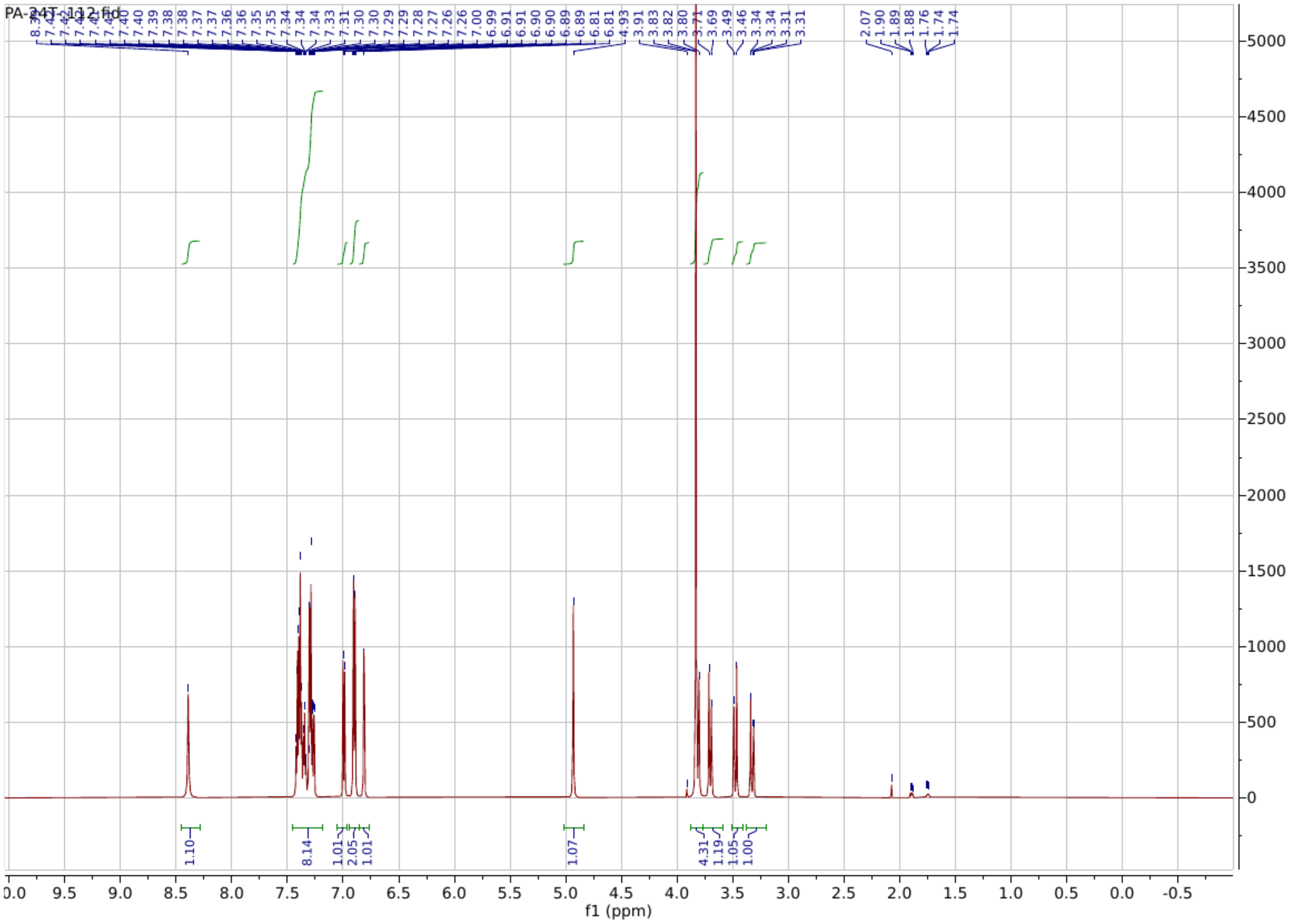

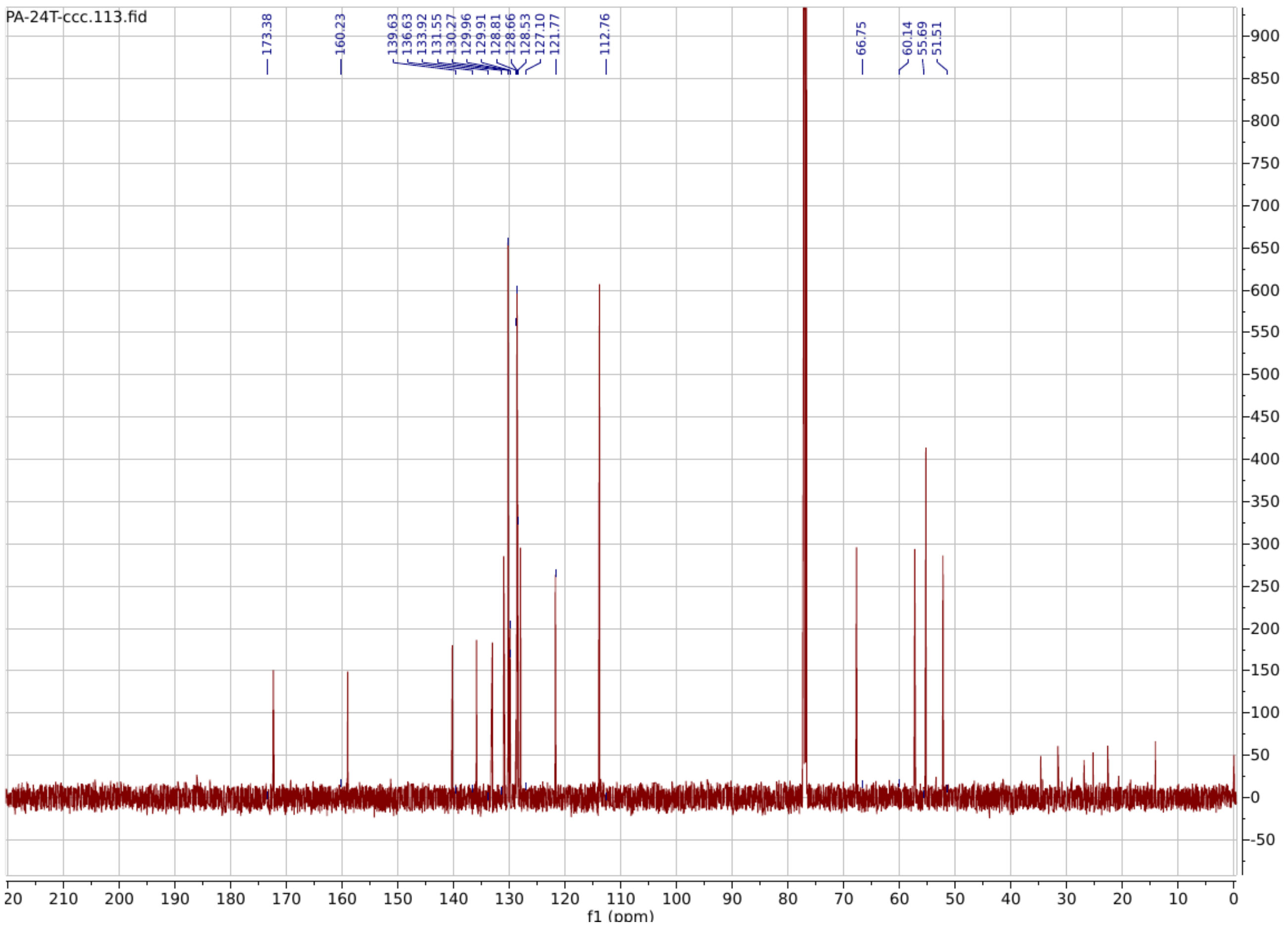

**PA90**

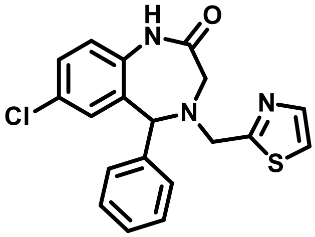

**7-chloro-5-phenyl-4-(thiazol-2-ylmethyl)-1,3,4,5-tetrahydro-2H-benzo[e][1,4]diazepin-2-one**

**$^1$H NMR** (600 MHz, $CDCl_3$) δ 8.31 – 8.06 (m, 1H), 7.72 (d, *J* = 3.3 Hz, 1H), 7.50 – 7.15 (m, 10H), 6.94 (dd, *J* = 14.4, 8.5 Hz, 1H), 6.86 (d, *J* = 2.4 Hz, 1H), 5.09 (s, 1H), 4.25 (d, *J* = 15.4 Hz, 1H), 4.13 (d, *J* = 15.4 Hz, 1H), 3.64 – 3.55 (m, 1H), 3.43 (dd, *J* = 15.6, 1.4 Hz, 1H).

**$^{13}$C NMR** (151 MHz, $CDCl_3$) δ 171.68, 169.81, 142.79, 139.41, 135.58, 132.23, 131.19, 130.20, 128.96, 128.94, 128.79, 128.51, 128.43, 128.35, 121.98, 119.59, 77.23, 77.02, 76.81, 68.09, 63.41, 55.29, 53.53.

**HRMS (ESI):** m/z calculated for $C_{19}H_{16}ClN_3OS$ $[M+H]^+$: 369.0703, found: 370.0776.

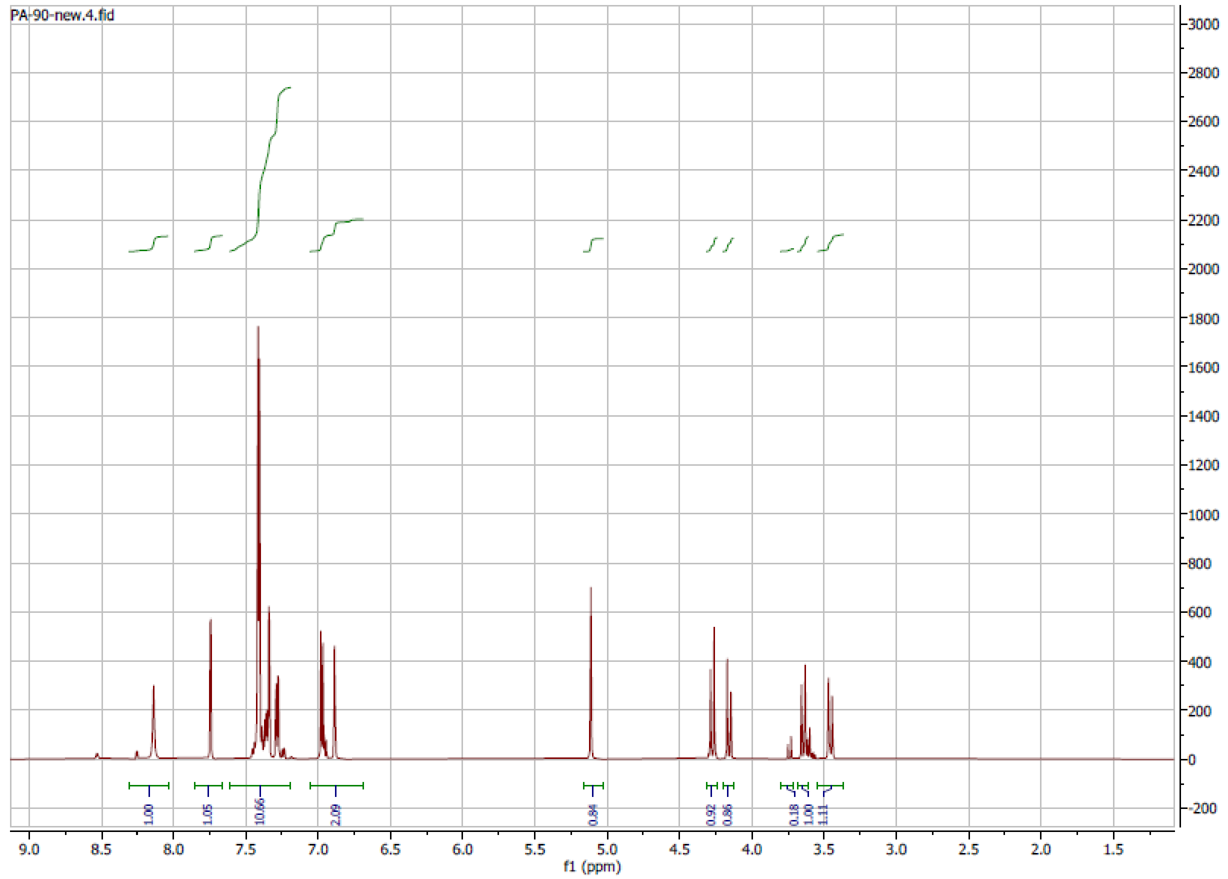

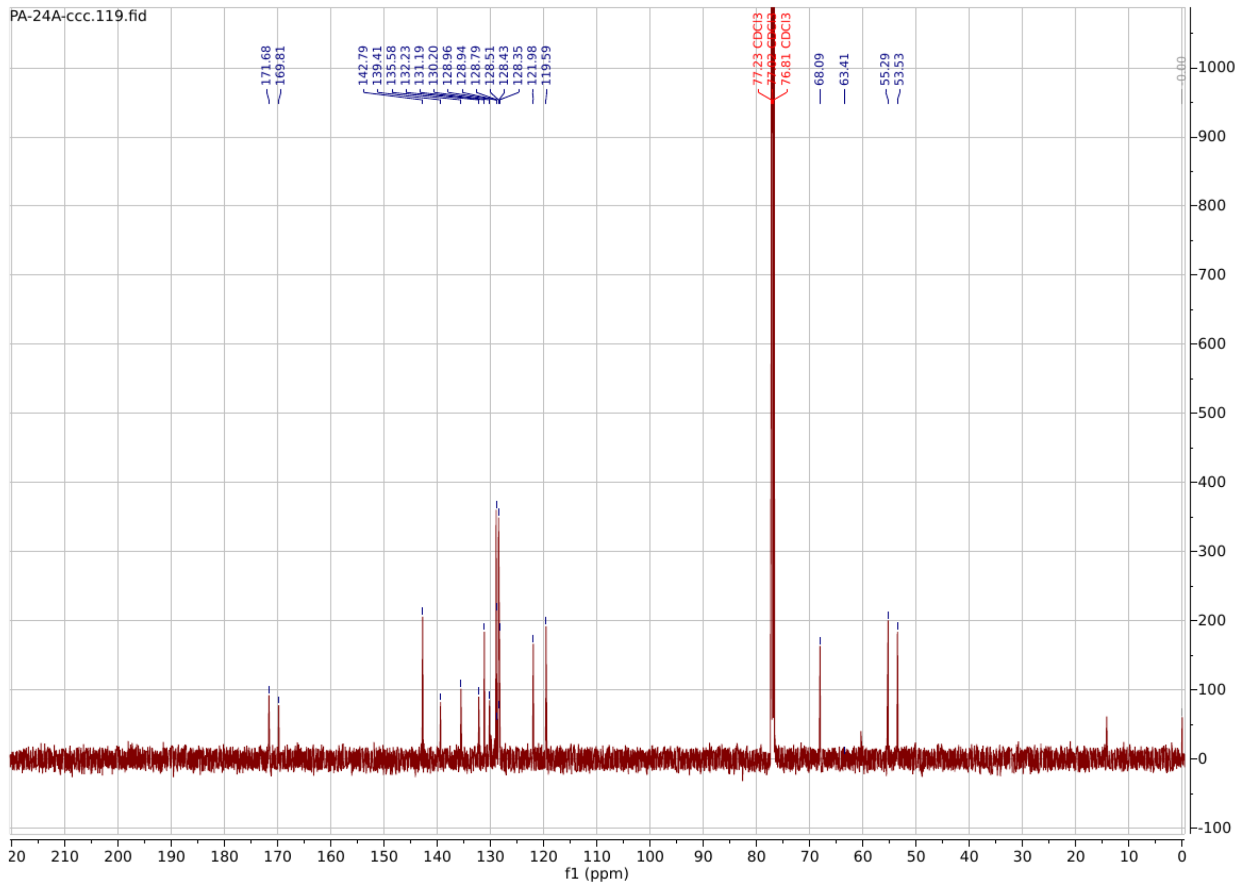

**PA91**

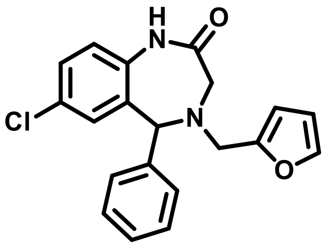

**7-chloro-4-(furan-2-ylmethyl)-5-phenyl-1,3,4,5-tetrahydro-2H-benzo[e][1,4]diazepin-2-one**

$^{1}$**H NMR** (600 MHz, Chloroform-*d*) δ 7.98 (s, 1H), 7.45 – 7.31 (m, 5H), 7.26 (dd, *J* = 8.4, 2.4 Hz, 1H), 6.94 (d, *J* = 8.5 Hz, 1H), 6.83 (d, *J* = 2.4 Hz, 1H), 6.35 (dd, *J* = 3.2, 1.9 Hz, 1H), 6.29 (d, *J* = 3.2 Hz, 1H), 4.98 (s, 1H), 3.91 – 3.78 (m, 2H), 3.52 (d, *J* = 15.4 Hz, 1H), 3.43 (dd, *J* = 15.3, 1.5 Hz, 1H).

**$^{13}$C NMR** (151 MHz, $CDCl_3$) δ 171.79, 151.43, 142.46, 139.79, 135.67, 133.07, 131.11, 130.13, 128.82, 128.62, 128.56, 128.07, 121.76, 110.23, 109.33, 77.22, 77.01, 76.80, 67.64, 52.74, 50.64.

**HRMS (ESI):** m/z calculated for $C_{20}H_{17}ClN_2O_2$ $[M+H]^+$: 352.0979, found: 353.1048.

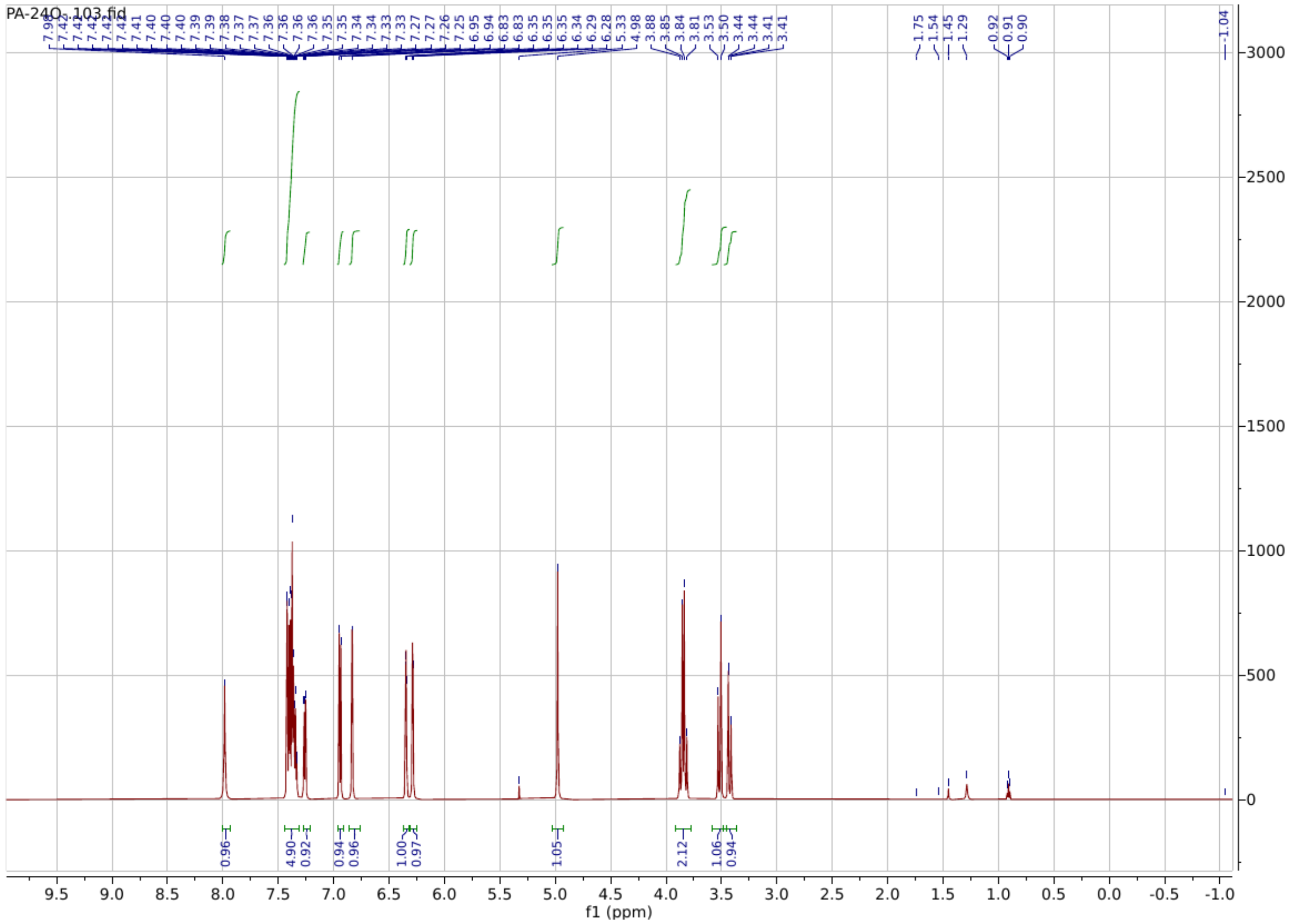

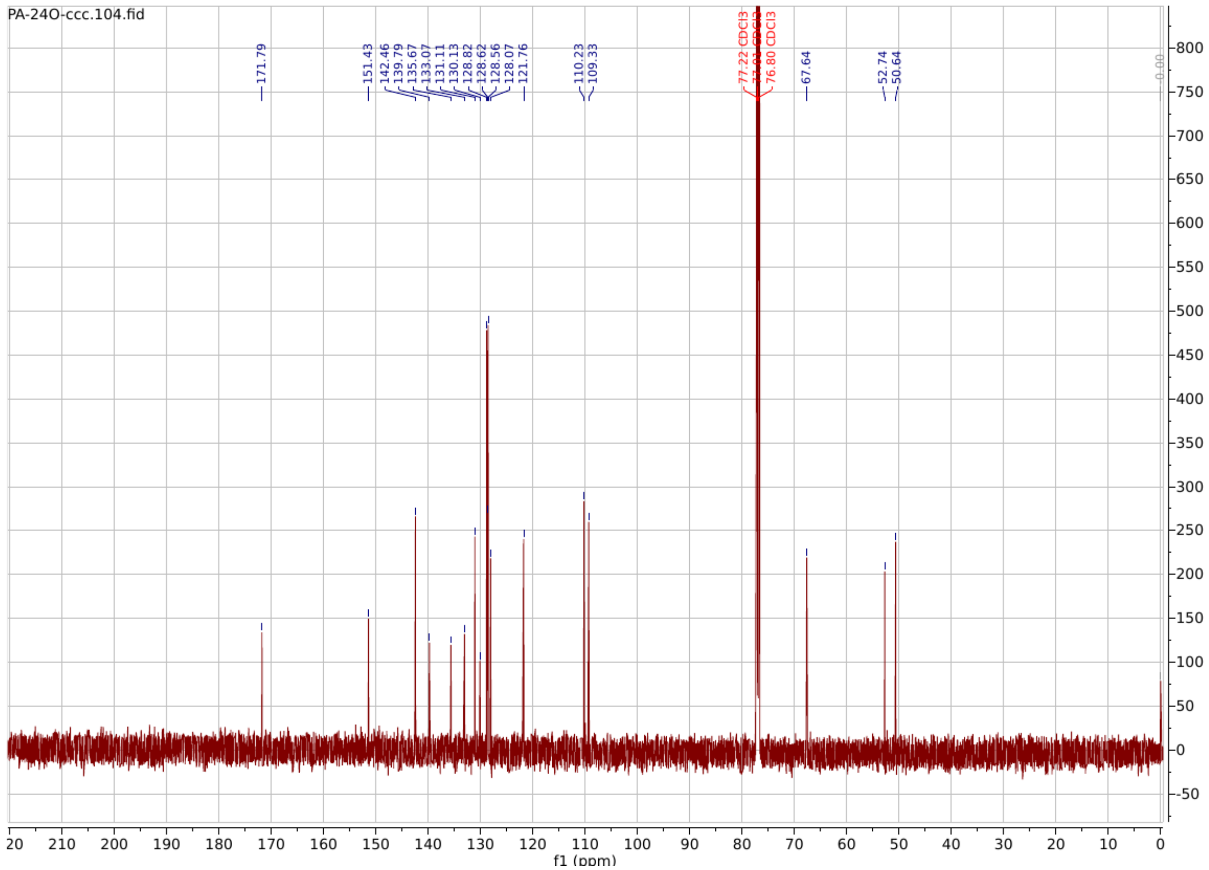

**PA92**

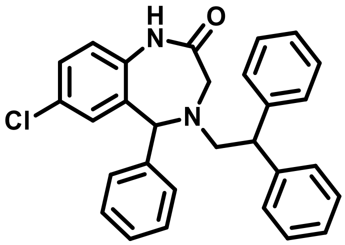

**7-chloro-4-(2,2-diphenylethyl)-5-phenyl-1,3,4,5-tetrahydro-2H-benzo[e][1,4]diazepin-2-one**

**$^1$H NMR** (400 MHz, $CDCl_3$) δ 8.39 (s, 1H), 7.39 – 7.13 (m, 17H), 6.93 (dd, $J$ = 8.1, 4.5 Hz, 3H), 6.86 (d, $J$ = 2.4 Hz, 1H), 4.97 (s, 1H), 4.39 (t, $J$ = 7.9 Hz, 1H), 3.50 (d, $J$ = 1.8 Hz, 2H), 3.37 (d, $J$ = 7.9 Hz, 2H).

**$^{13}$C NMR** (151 MHz, $CDCl_3$) δ 174.16, 142.61, 142.46, 140.12, 135.43, 132.02, 131.11, 129.29, 128.75, 128.58, 128.49, 128.41, 128.40, 128.17, 127.81, 126.56, 121.75, 77.25, 77.04, 76.83, 68.93, 60.43, 58.31, 52.69, 49.43.

**HRMS (ESI):** m/z calculated for $C_{29}H_{25}ClN_2O$ $[M+H]^+$: 452.1655, found: 453.1731.

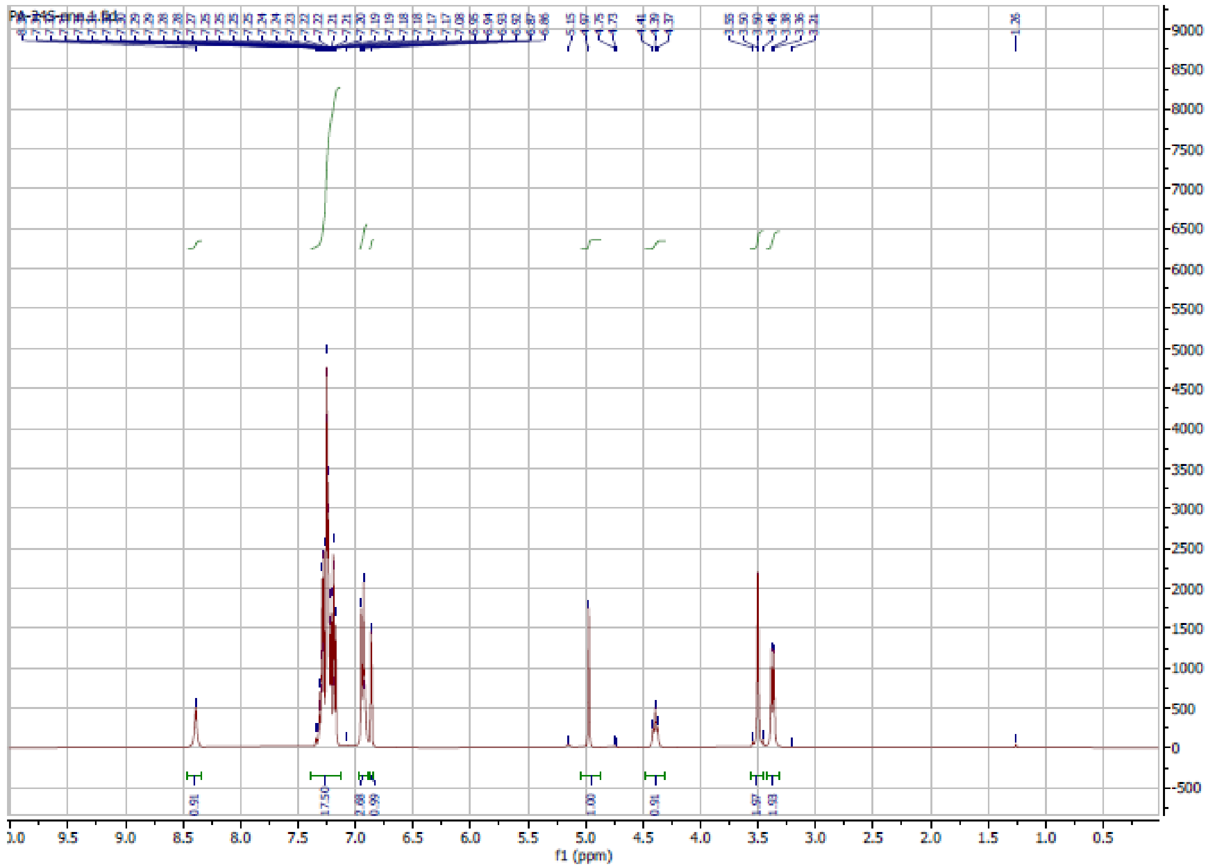

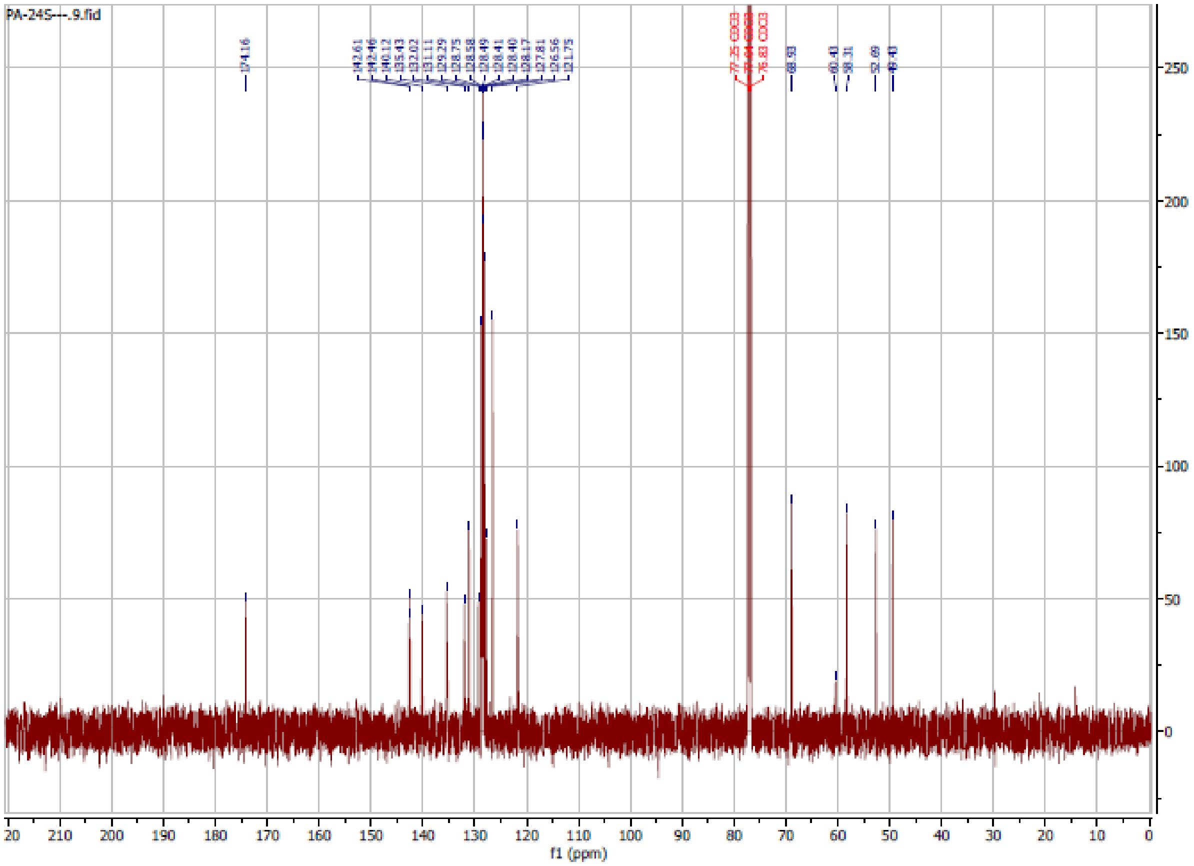

**PA93**

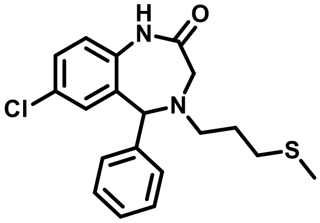

**7-chloro-4-(3-(methylthio)propyl)-5-phenyl-1,3,4,5-tetrahydro-2H-benzo[e][1,4]diazepin-2-one**

**$^{1}$H NMR** (600 MHz, $CDCl_3$) δ 9.34 (s, 1H), 7.46 – 7.18 (m, 6H), 7.05 (dd, *J* = 9.0, 3.2 Hz, 1H), 6.90 (s, 1H), 4.99 (s, 1H), 3.55 (dd, *J* = 16.2, 3.3 Hz, 1H), 3.46 – 3.35 (m, 1H), 2.82 (t, *J* = 7.3 Hz, 2H), 2.56 (td, *J* = 14.6, 13.7, 6.3 Hz, 2H), 2.10 (s, 3H), 1.87 (q, *J* = 7.8 Hz, 2H).

**$^{13}$C NMR** (151 MHz, $CDCl_3$) δ 173.90, 140.38, 135.74, 132.58, 130.98, 129.59, 128.75,

128.64, 128.55, 127.94, 121.92, 77.28, 77.07, 76.86, 68.73, 52.83, 52.17, 31.57, 26.95, 15.49.

**HRMS (ESI):** m/z calculated for $C_{19}H_{21}ClN_2OS$ $[M+H]^+$: 360.1063, found: 361.1133.

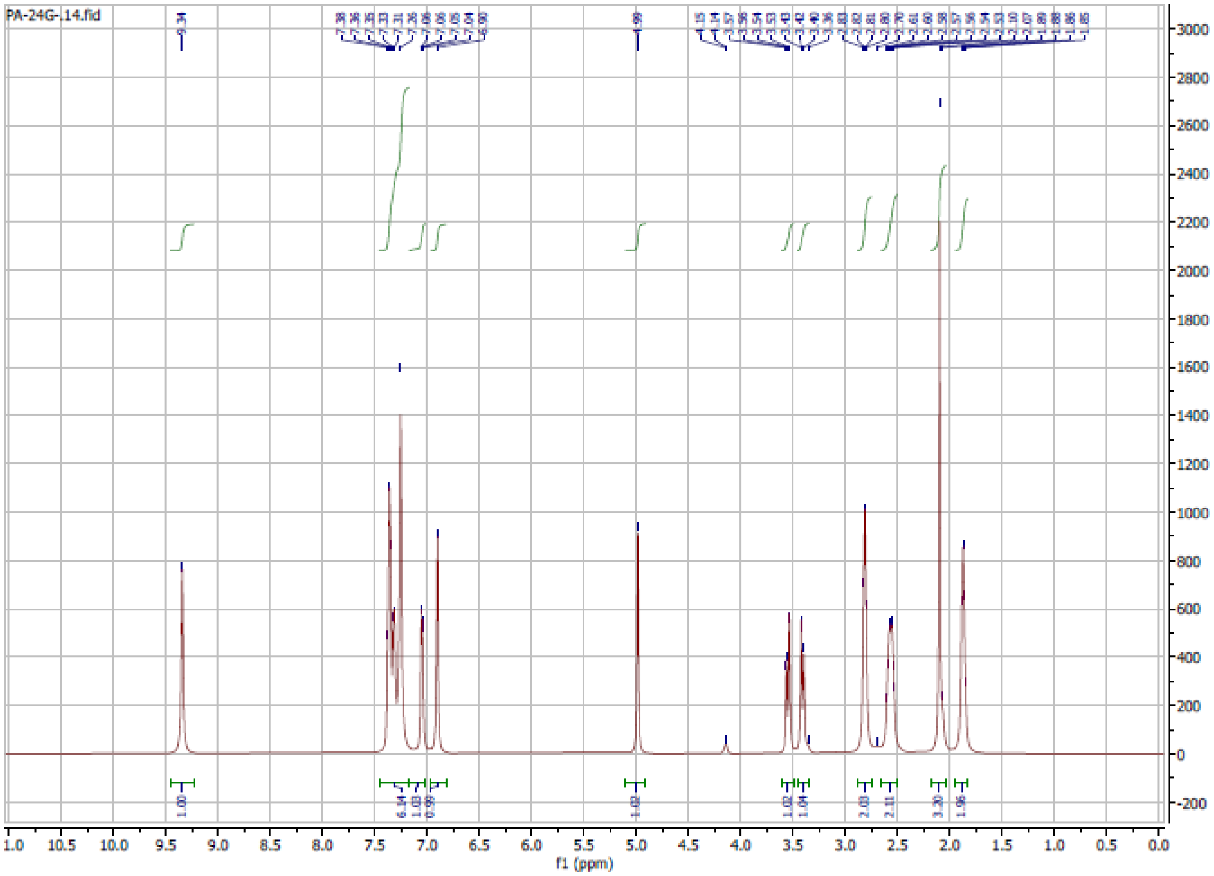

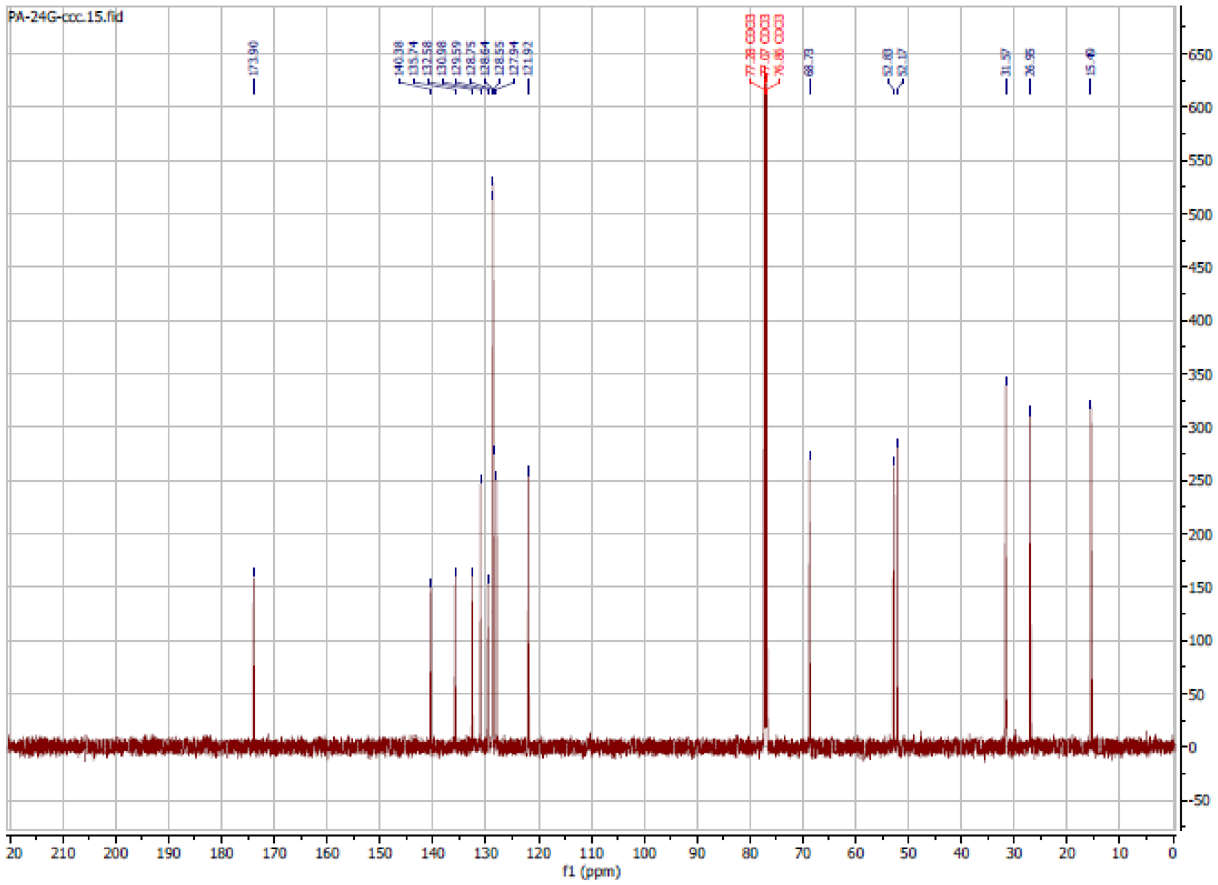

**PA97**

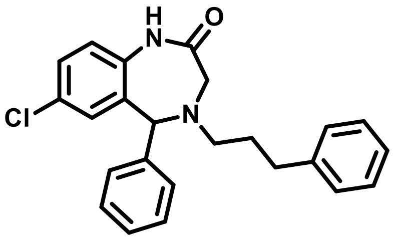

**7-chloro-5-phenyl-4-(3-phenylpropyl)-1,3,4,5-tetrahydro-2H-benzo[e][1,4]diazepin-2-one**

**$^{1}$H NMR** (600 MHz, Chloroform-*d*) δ 8.18 (s, 1H), 7.42 – 7.12 (m, 12H), 6.97 – 6.74 (m, 2H), 4.98 (s, 1H), 3.56 (d, *J* = 15.9 Hz, 1H), 3.43 (dd, *J* = 15.9, 1.4 Hz, 1H), 2.81 – 2.61 (m, 5H), 1.99 – 1.81 (m, 2H).

**$^{13}$C NMR** (151 MHz, $CDCl_3$) δ 173.06, 141.89, 141.82, 140.44, 135.52, 132.67, 131.13, 129.67, 128.73, 128.58, 128.51, 128.43, 128.41, 128.36, 127.91, 125.88, 125.84, 121.62, 77.23, 77.02, 76.80, 68.51, 62.31, 52.91, 52.86, 34.24, 33.08, 32.09, 29.21.

**HRMS (ESI):** m/z calculated for $C_{24}H_{23}ClN_2O$ $[M+H]^+$: 390.1499, found: 391.1573.

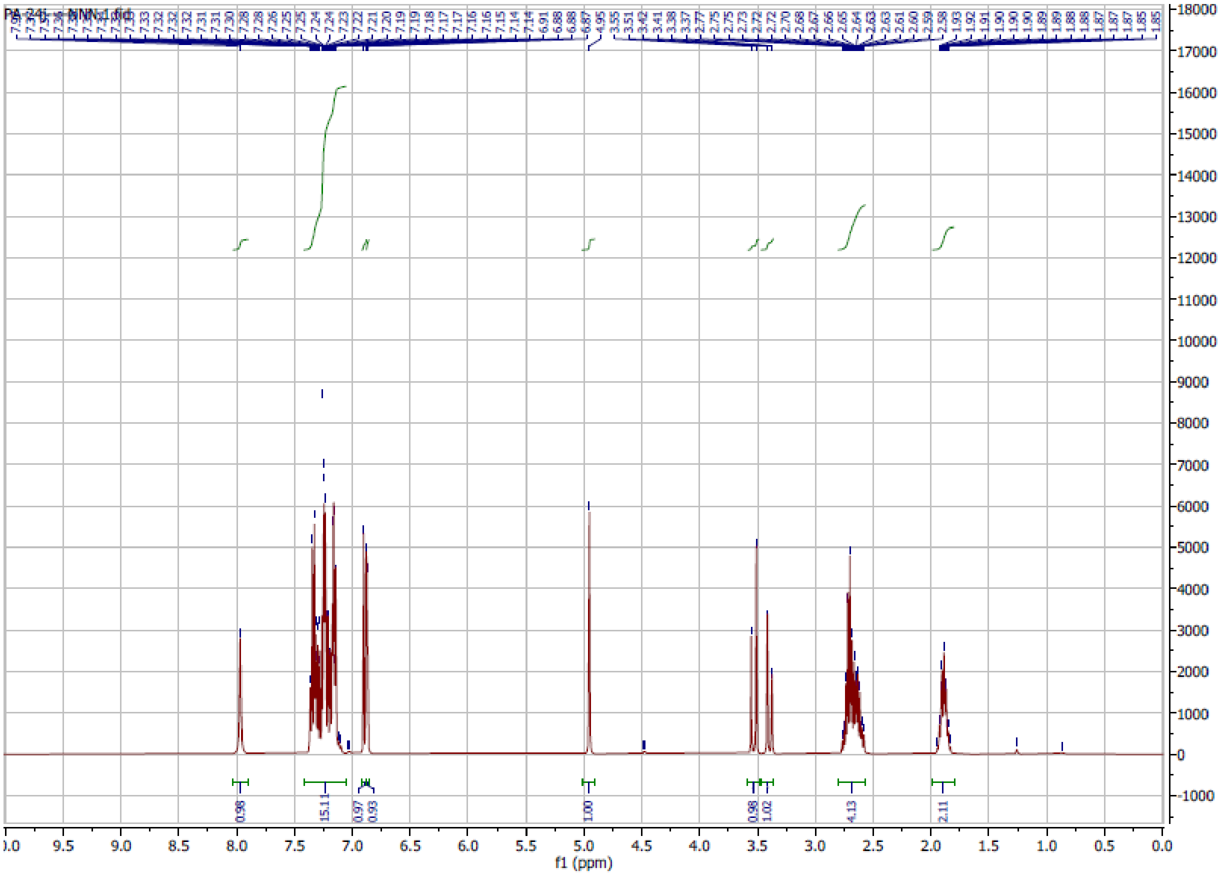

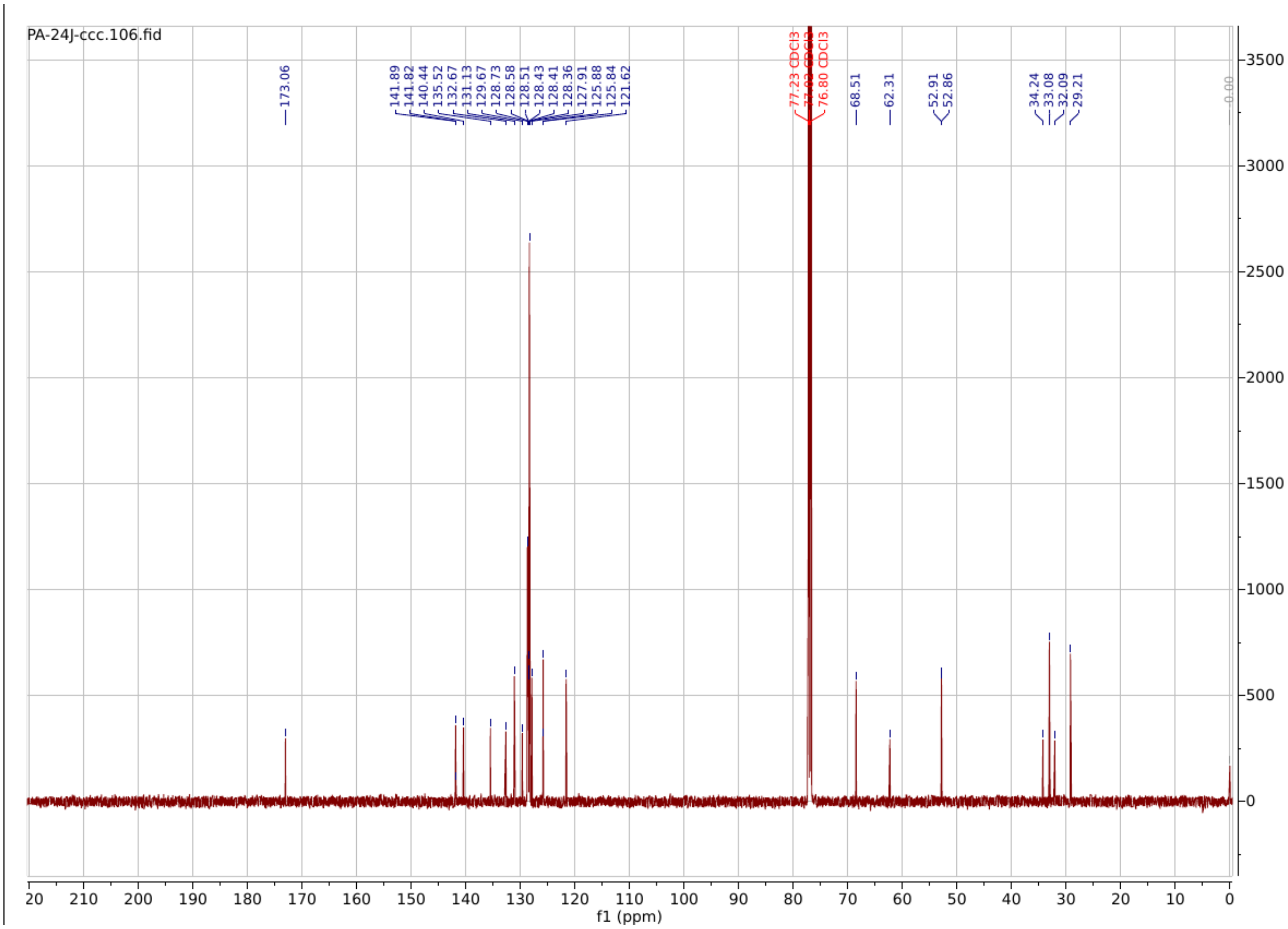

**PA99**

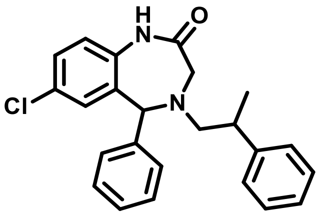

**7-chloro-5-phenyl-4-(2-phenylpropyl)-1,3,4,5-tetrahydro-2H-benzo[e][1,4]diazepin-2-one**

**$^1$H NMR** (600 MHz, Chloroform-*d*) δ 8.34 (d, *J* = 9.0 Hz, 1H), 7.39 – 7.06 (m, 9H), 7.06 – 6.87 (m, 3H), 6.79 (d, *J* = 2.4 Hz, 1H), 4.91 (d, *J* = 86.1 Hz, 1H), 3.60 – 3.27 (m, 2H), 3.16 – 2.64 (m, 3H), 1.30 (dd, *J* = 26.5, 6.9 Hz, 3H).

**$^{13}$C NMR** (151 MHz, $CDCl_3$) δ 173.69, 145.15, 145.00, 140.14, 135.45, 132.50, 131.15, 129.40, 128.71, 128.69, 128.65, 128.54, 128.45, 128.43, 128.29, 127.87, 127.72, 127.37, 127.31, 126.38, 126.30, 121.62, 121.57, 77.23, 77.02, 76.81, 69.15, 68.92, 60.85, 60.78, 53.42, 52.99, 52.72, 38.27, 37.93, 19.86, 19.11, 14.12.

**HRMS (ESI):** m/z calculated for $C_{24}H_{23}ClN_2O$ $[M+H]^+$: 390.1499, found: 391.1566.

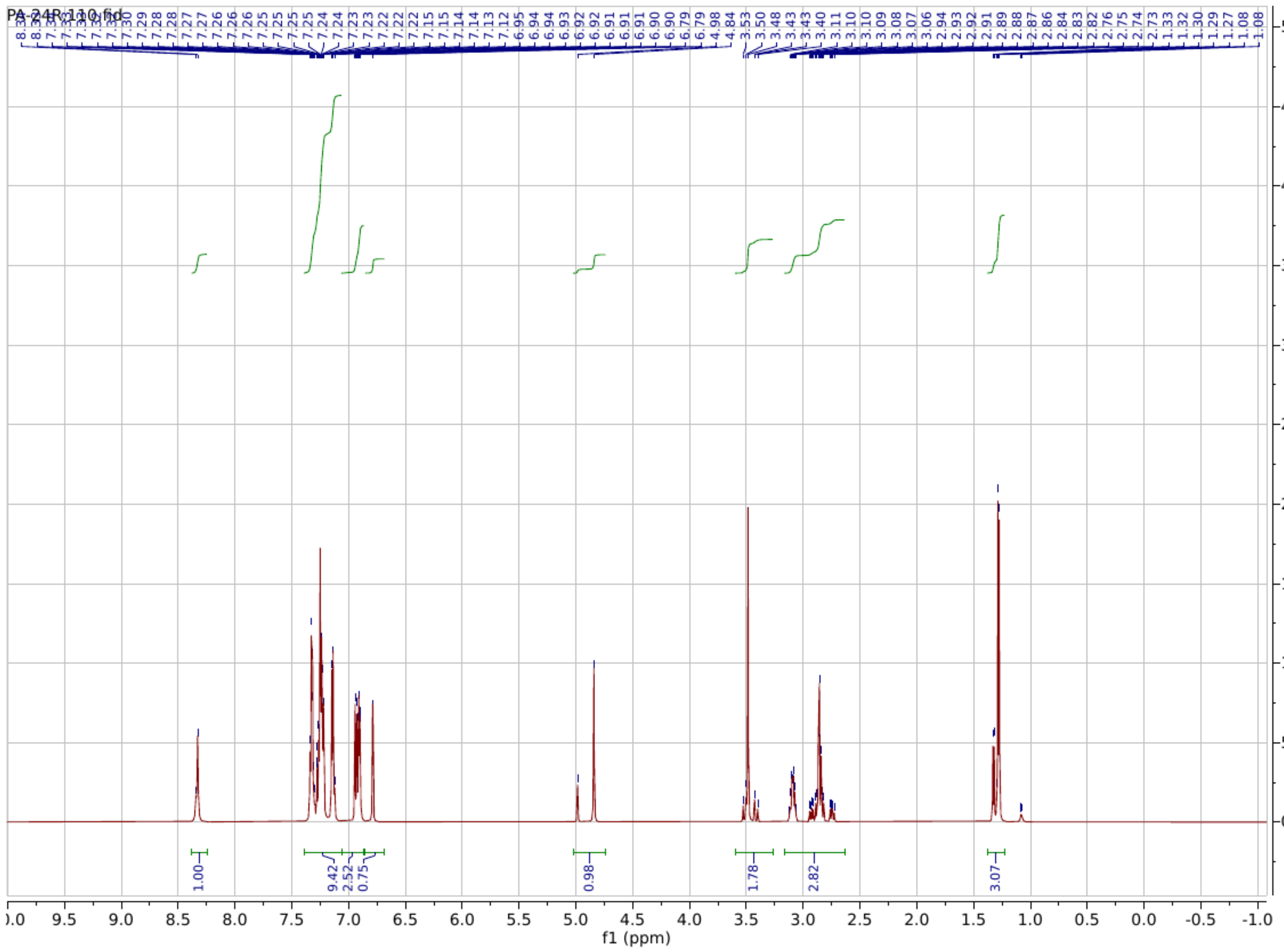

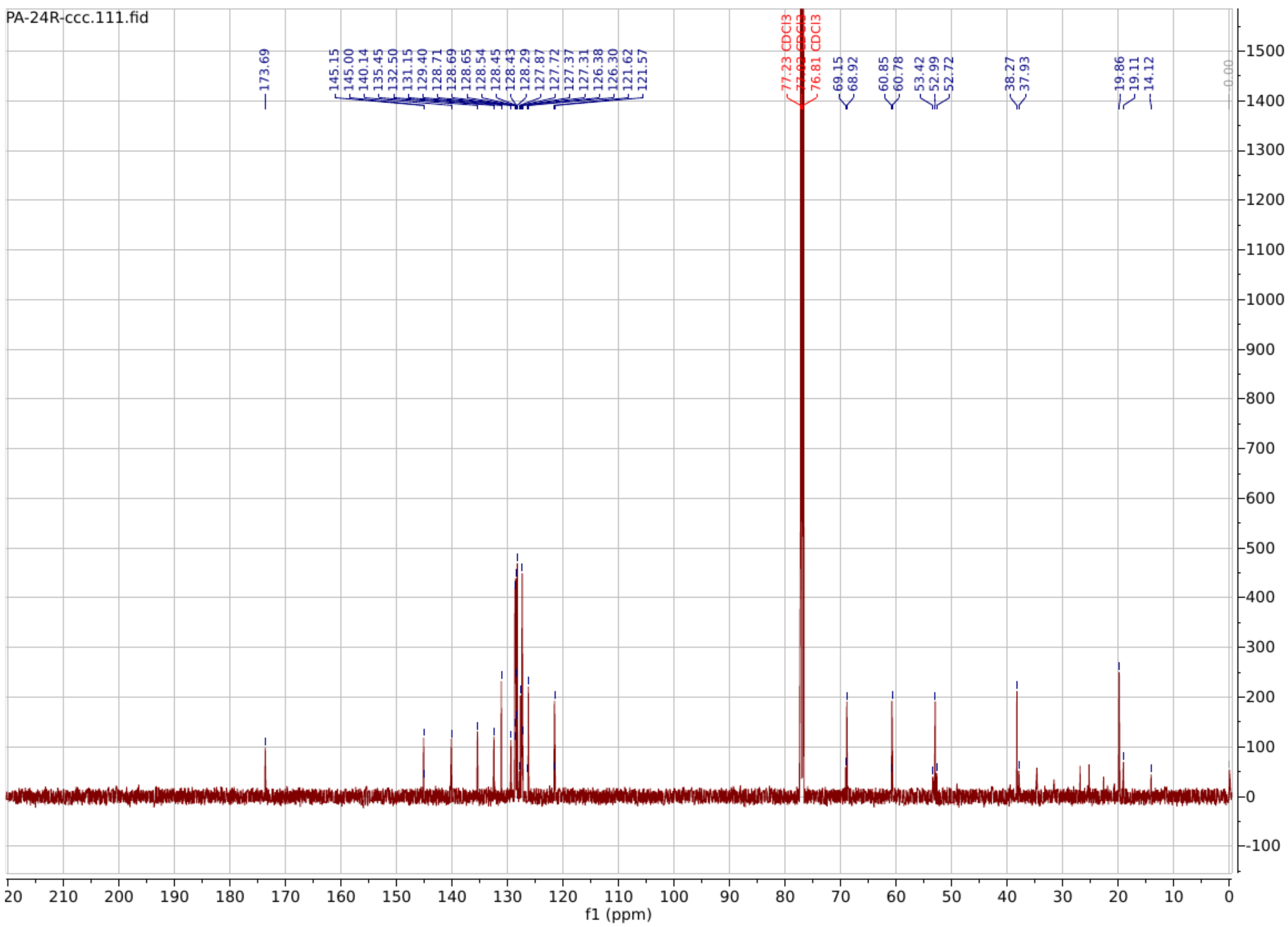

**PA133**

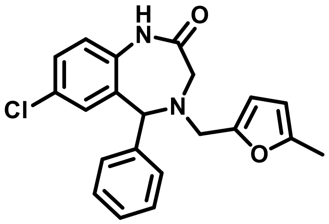

**7-chloro-4-((5-methylfuran-2-yl)methyl)-5-phenyl-1,3,4,5-tetrahydro-2H-benzo[e][1,4]diazepin-2-one**

**$^1$H NMR** (600 MHz, Chloroform-*d*) δ 8.11 (s, 1H), 7.42 – 7.31 (m, 5H), 7.26 (dd, *J* = 8.5, 2.4 Hz, 2H), 6.94 (d, *J* = 8.5 Hz, 1H), 6.87 (d, *J* = 2.4 Hz, 1H), 6.14 (d, *J* = 3.0 Hz, 1H), 5.91 (dd, *J* = 3.1, 1.1 Hz, 1H), 5.01 (s, 1H), 3.88 – 3.73 (m, 2H), 3.57 – 3.44 (m, 2H), 2.35 – 2.25 (m, 3H).

**$^{13}$C NMR** (151 MHz, $CDCl_3$) δ 152.22, 145.89, 135.56, 131.21, 130.00, 128.77, 128.61, 128.53, 127.99, 121.78, 113.04, 110.32, 106.13, 77.23, 77.08, 77.02, 76.87, 76.81, 67.49, 52.91, 50.88, 13.66.

**HRMS (ESI):** m/z calculated for $C_{21}H_{19}ClN_2O_2$ $[M+H]^+$: 366.1135, found: 367.1198.

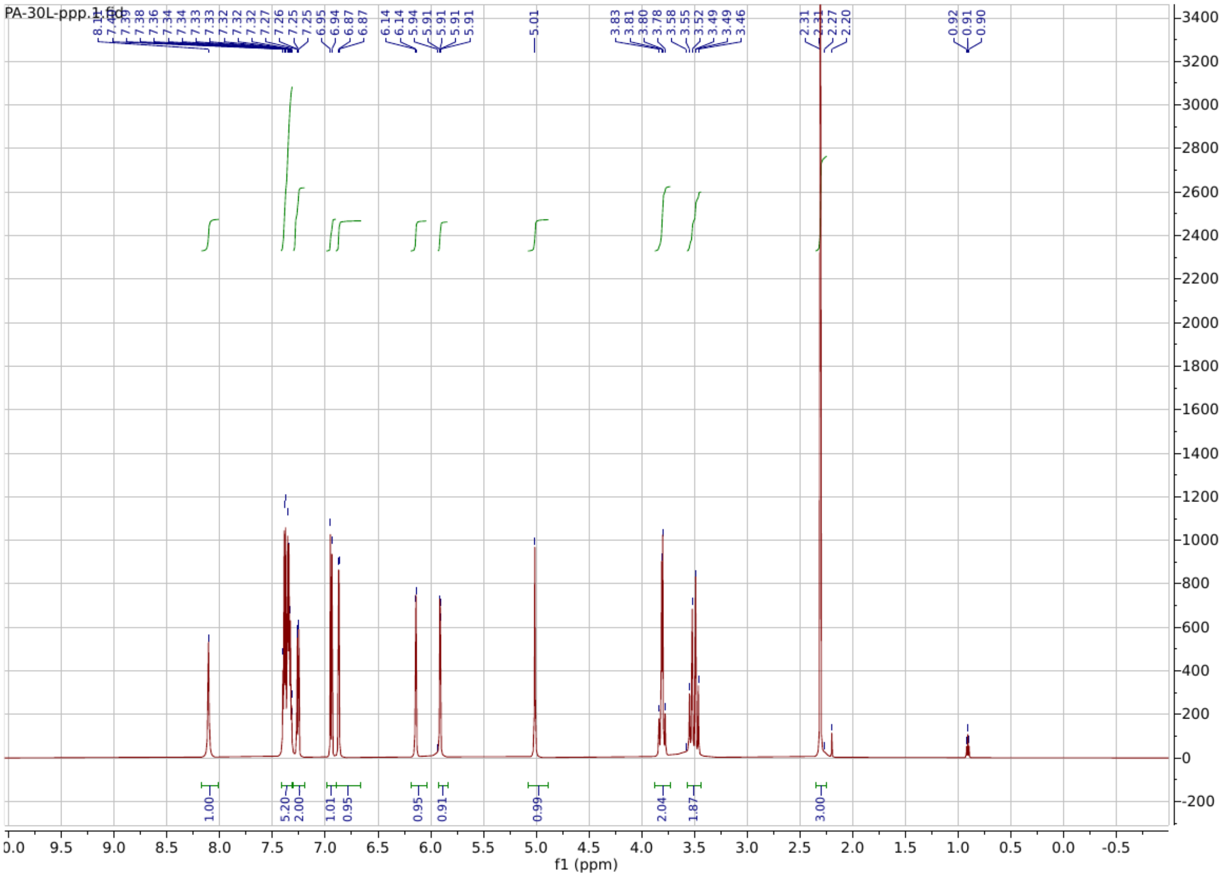

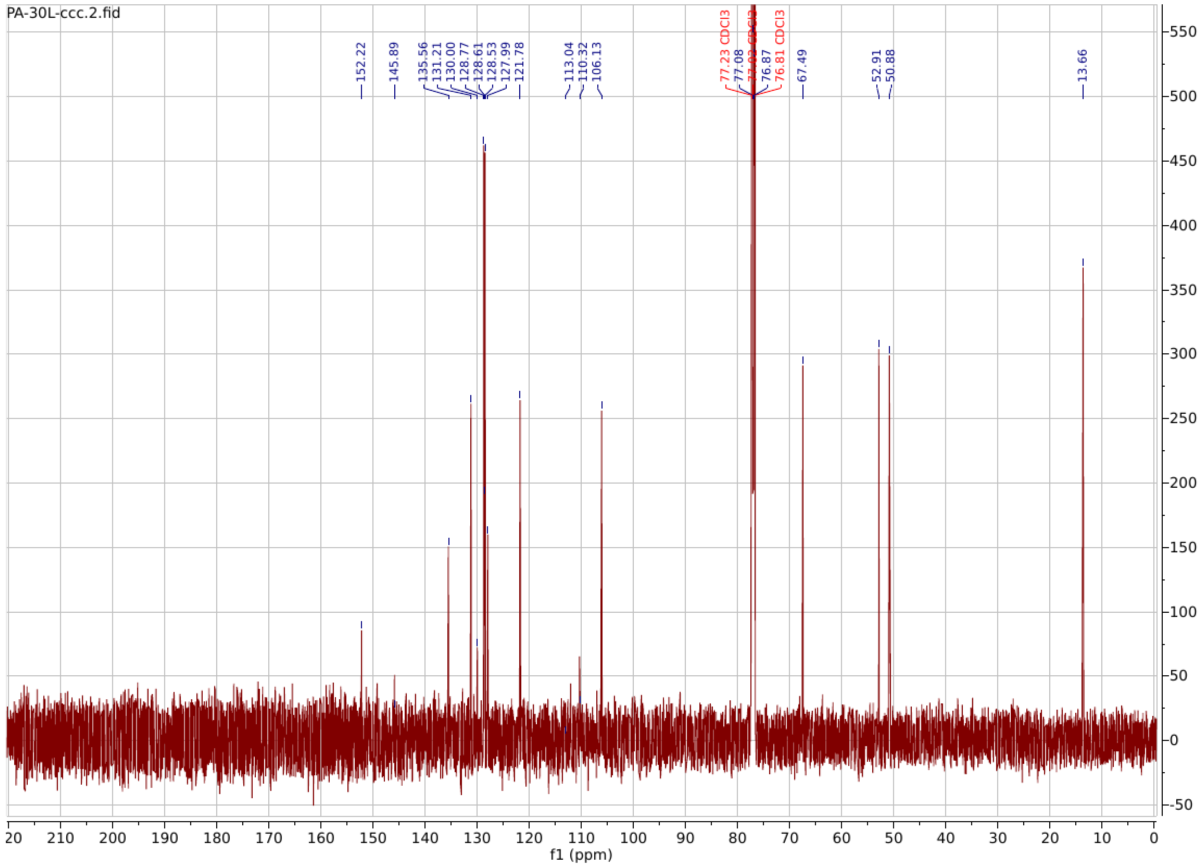

**PA135**

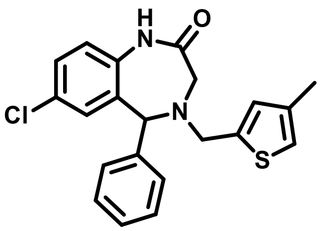

**7-chloro-4-((4-methylthiophen-2-yl)methyl)-5-phenyl-1,3,4,5-tetrahydro-2H-benzo[e][1,4]diazepin-2-one**

**$^{1}$H NMR** (600 MHz, Chloroform-*d*) δ 7.91 (s, 1H), 7.44 – 7.40 (m, 4H), 7.36 (ddt, *J* = 8.5, 5.5, 2.8 Hz, 1H), 7.28 – 7.26 (m, 1H), 6.95 (d, *J* = 8.4 Hz, 1H), 6.86 – 6.82 (m, 2H), 6.79 (d, *J* = 1.5 Hz, 1H), 5.02 (s, 1H), 3.97 (s, 2H), 3.54 (d, *J* = 15.4 Hz, 1H), 3.46 (dd, *J* = 15.4, 1.4 Hz, 1H), 2.23 (d, *J* = 1.0 Hz, 3H).

**$^{13}$C NMR** (151 MHz, $CDCl_3$) δ 137.21, 135.72, 132.80, 131.17, 130.26, 129.16, 128.92, 128.74, 128.63, 128.22, 121.76, 120.73, 77.23, 77.02, 76.80, 67.41, 52.75, 52.35, 15.72.

**HRMS (ESI):** m/z calculated for $C_{21}H_{19}ClN_2OS$ $[M+H]^+$: 382.0907, found: 383.0973.

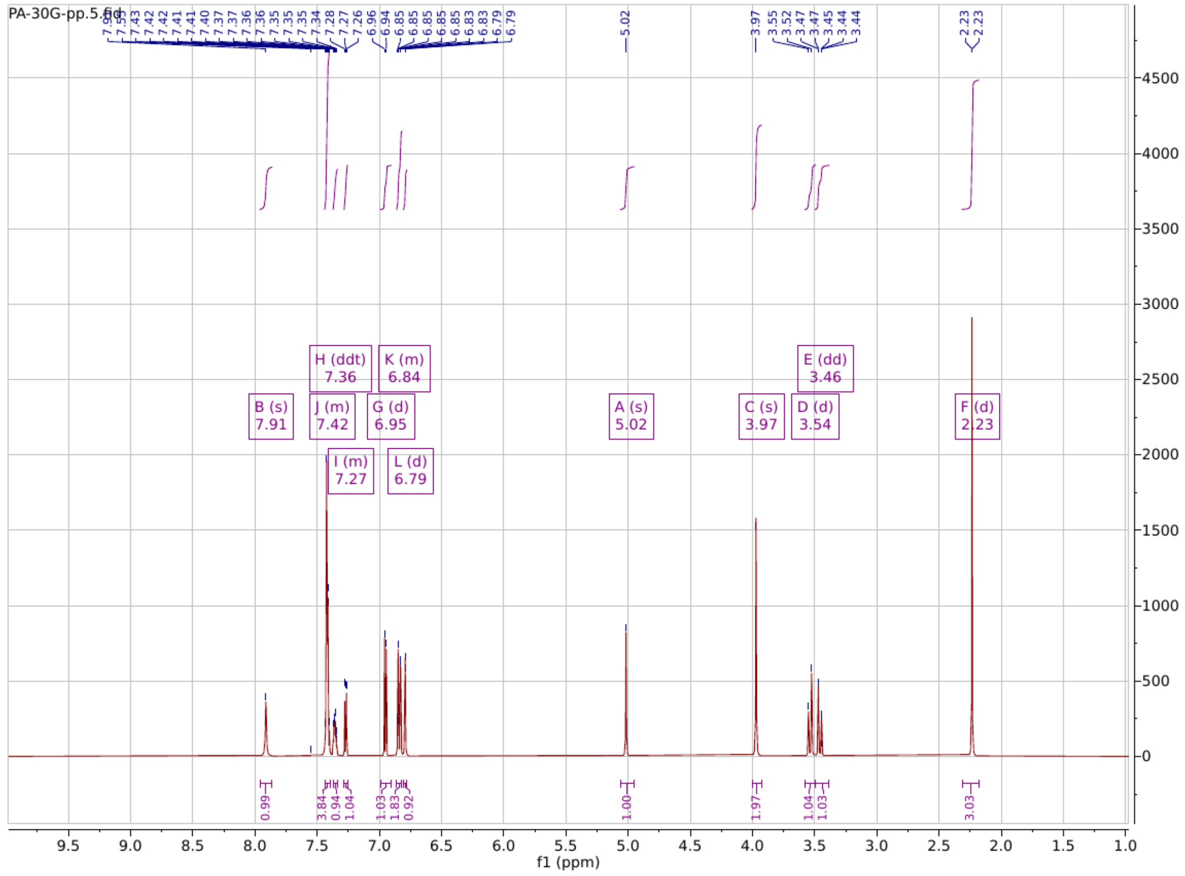

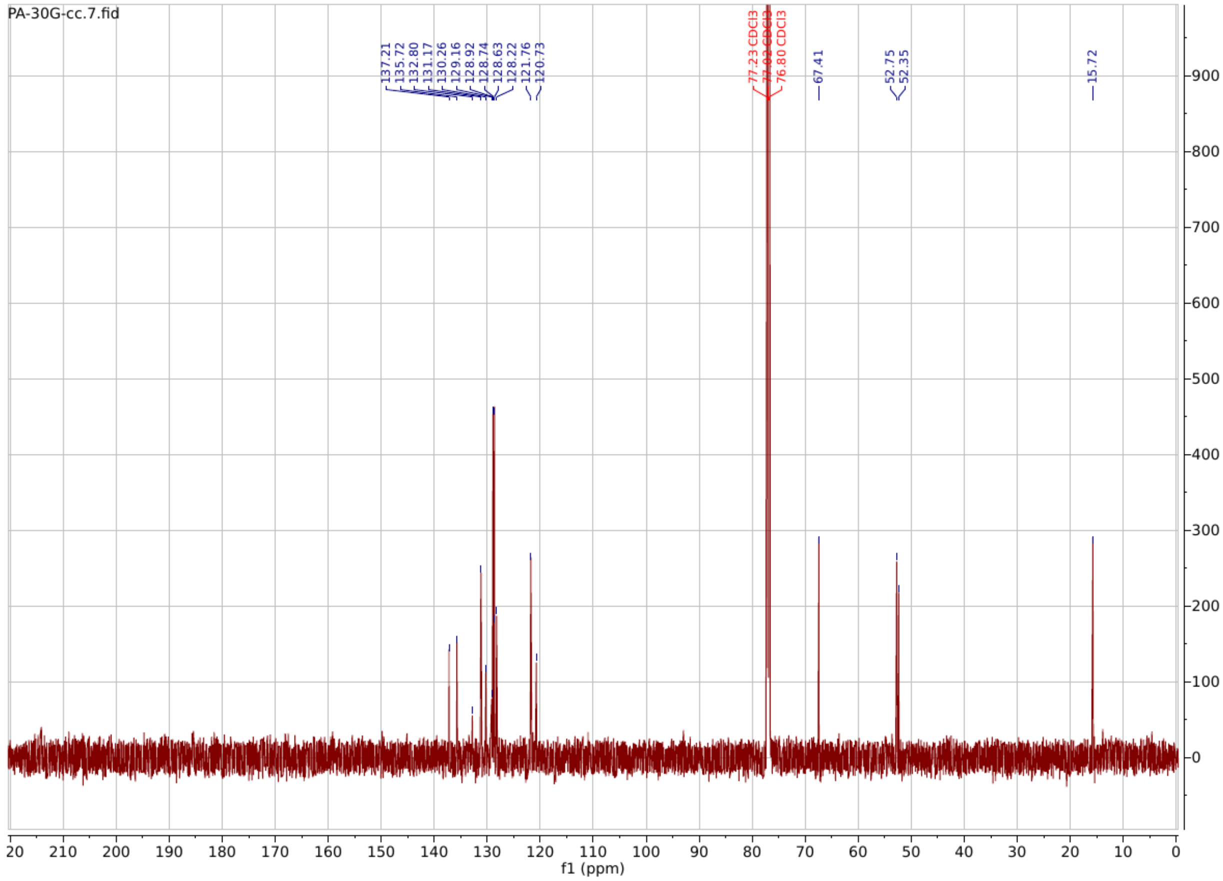

**PA136**

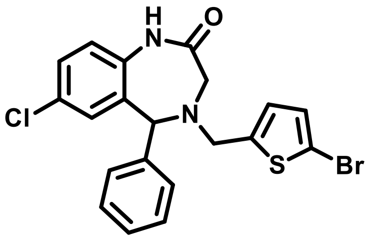

**4-((5-bromothiophen-2-yl)methyl)-7-chloro-5-phenyl-1,3,4,5-tetrahydro-2H-benzo[e][1,4]diazepin-2-one**

**$^{1}$H NMR** (600 MHz, Chloroform-*d*) δ 7.71 (s, 1H), 7.50 – 7.31 (m, 5H), 7.27 (d, *J* = 2.4 Hz, 6H), 6.98 – 6.85 (m, 2H), 6.80 (d, *J* = 2.4 Hz, 1H), 6.79 – 6.67 (m, 1H), 4.97 (s, 1H), 4.02 – 3.85 (m, 2H), 3.53 (d, *J* = 15.3 Hz, 1H), 3.41 (dd, *J* = 15.3, 1.6 Hz, 1H).

**$^{13}C$ NMR** (151 MHz, $CDCl_3$) δ 168.58, 135.72, 131.10, 130.50, 129.42, 129.05, 128.87, 128.60, 128.44, 127.02, 126.19, 121.81, 77.23, 77.01, 76.80, 67.52, 52.95, 52.20.

**HRMS (ESI):** m/z calculated for $C_{20}H_{16}BrClN_2OS$ $[M+H]^+$: 445.9855, found: 446.9919.

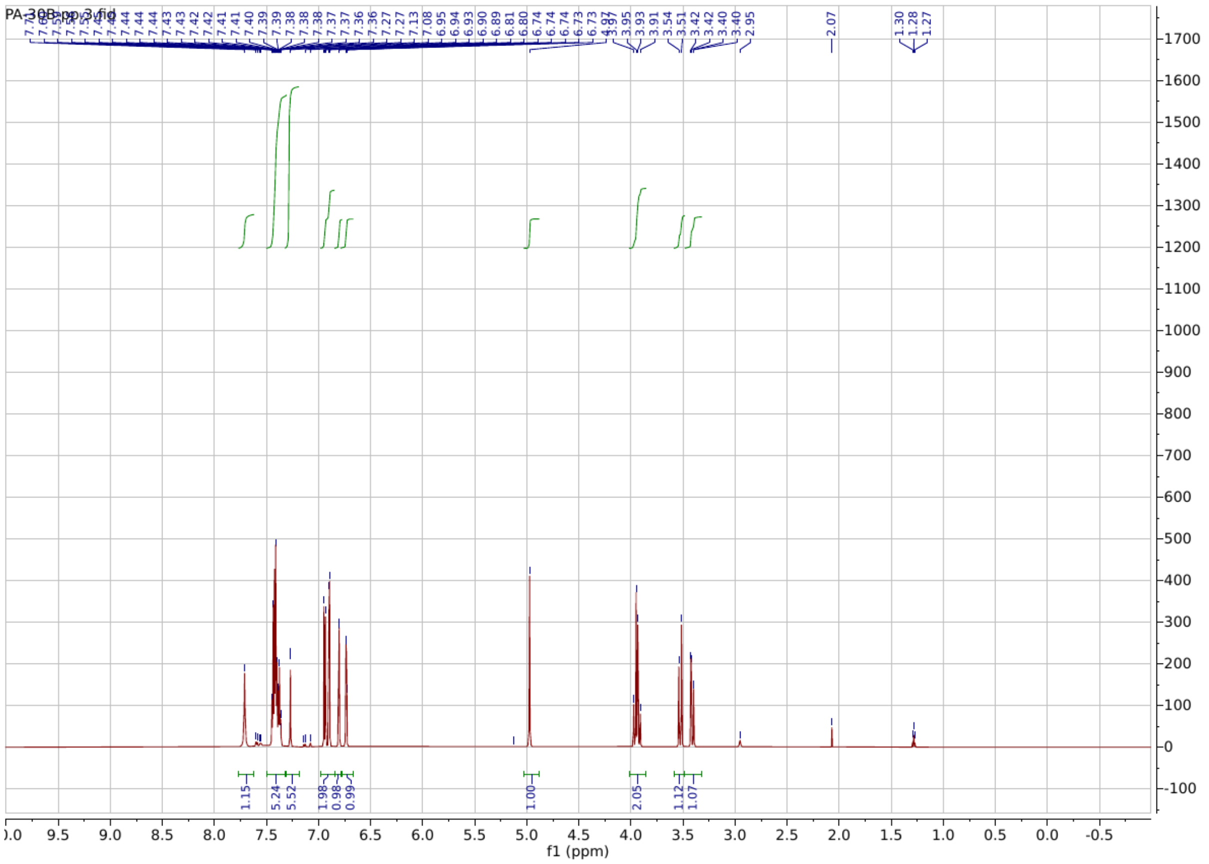

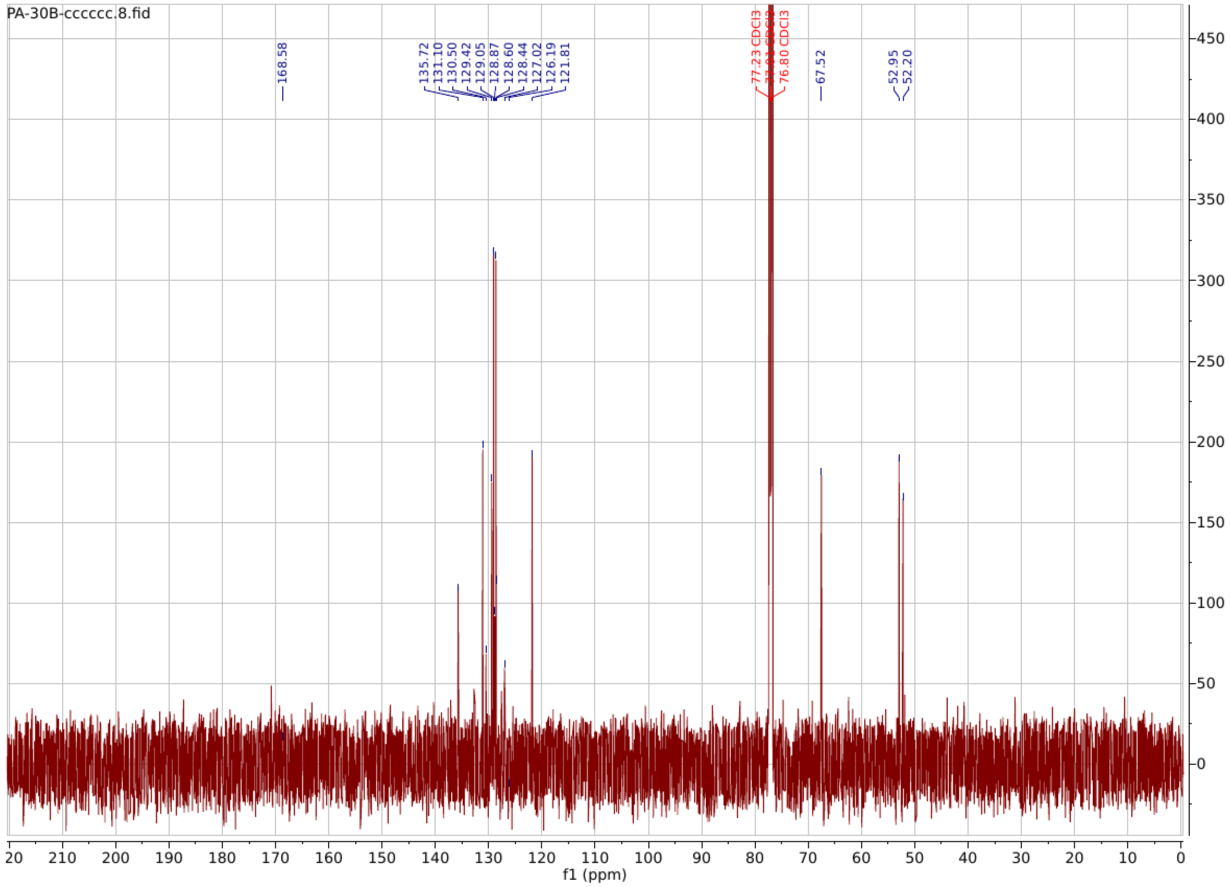

**PA138**

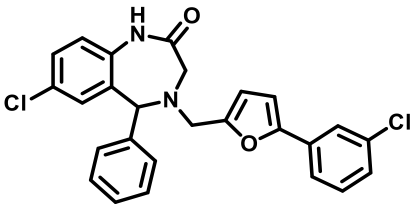

**7-chloro-4-((5-(3-chlorophenyl)furan-2-yl)methyl)-5-phenyl-1,3,4,5-tetrahydro-2H-benzo[e][1,4]diazepin-2-one**

**$^1$H NMR** (600 MHz, Chloroform-*d*) δ 8.06 (s, 1H), 7.65 (t, *J* = 1.9 Hz, 1H), 7.53 (dt, *J* = 7.9, 1.3 Hz, 1H), 7.45 – 7.40 (m, 4H), 7.37 (ddd, *J* = 8.6, 5.6, 2.5 Hz, 1H), 7.33 (t, *J* = 7.9 Hz, 1H), 7.29 – 7.17 (m, 3H), 6.94 (d, *J* = 8.4 Hz, 1H), 6.84 (d, *J* = 2.4 Hz, 1H), 6.62 (d, *J* = 3.3 Hz, 1H), 6.38 (d, *J* = 3.3 Hz, 1H), 5.04 (s, 1H), 3.92 (d, *J* = 3.5 Hz, 2H), 3.58 (d, *J* = 15.3 Hz, 1H), 3.49 (dd, *J* = 15.3, 1.4 Hz, 1H).

**$^{13}C$ NMR** (151 MHz, $CDCl_3$) δ 171.71, 152.33, 135.69, 134.71, 133.09, 132.41, 131.08, 130.27, 129.96, 128.86, 128.69, 128.66, 128.21, 127.18, 123.71, 121.85, 121.71, 111.56, 106.72, 77.23, 77.02, 76.81, 67.62, 53.04, 50.83.

**HRMS (ESI):** m/z calculated for $C_{26}H_{20}Cl_2N_2O_2$ $[M+H]^+$: 462.0902, found: 463.0964.

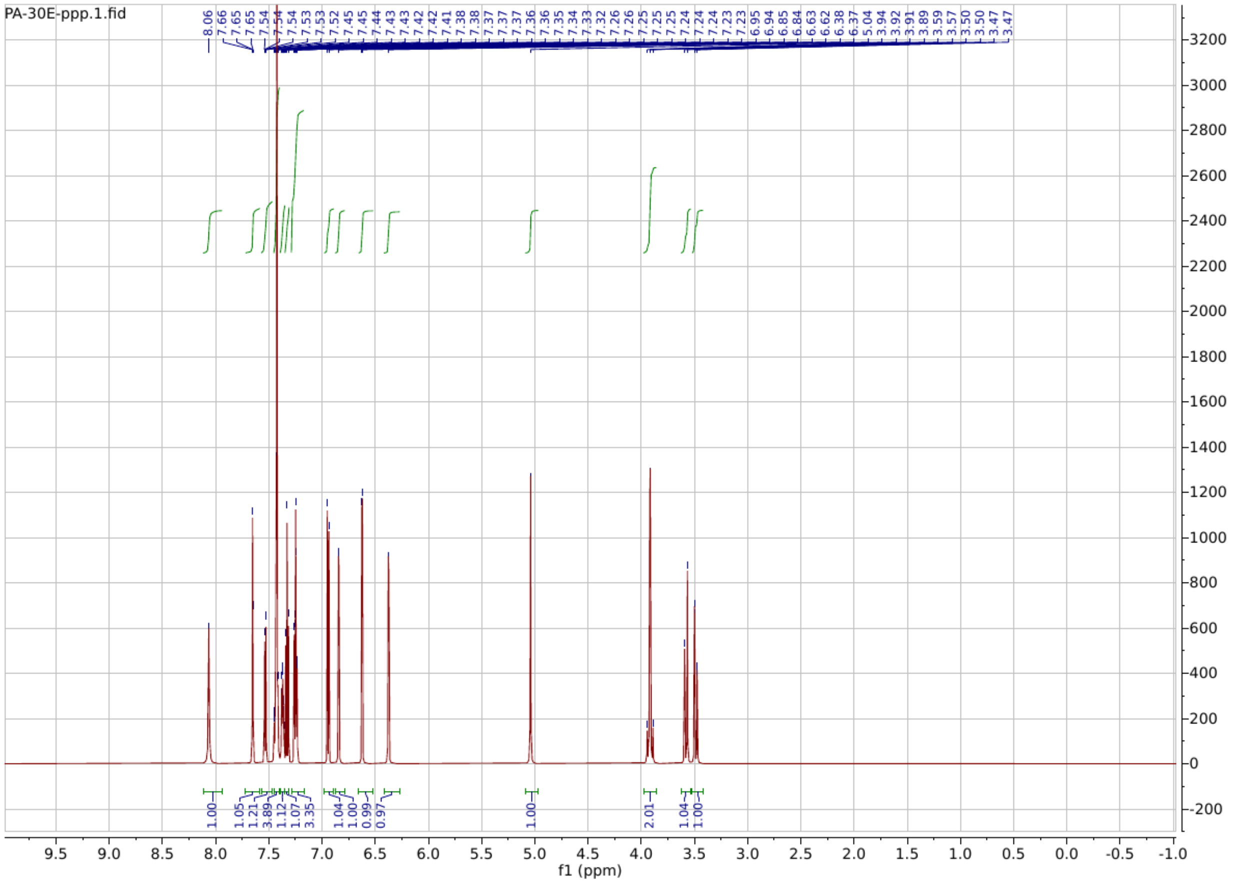

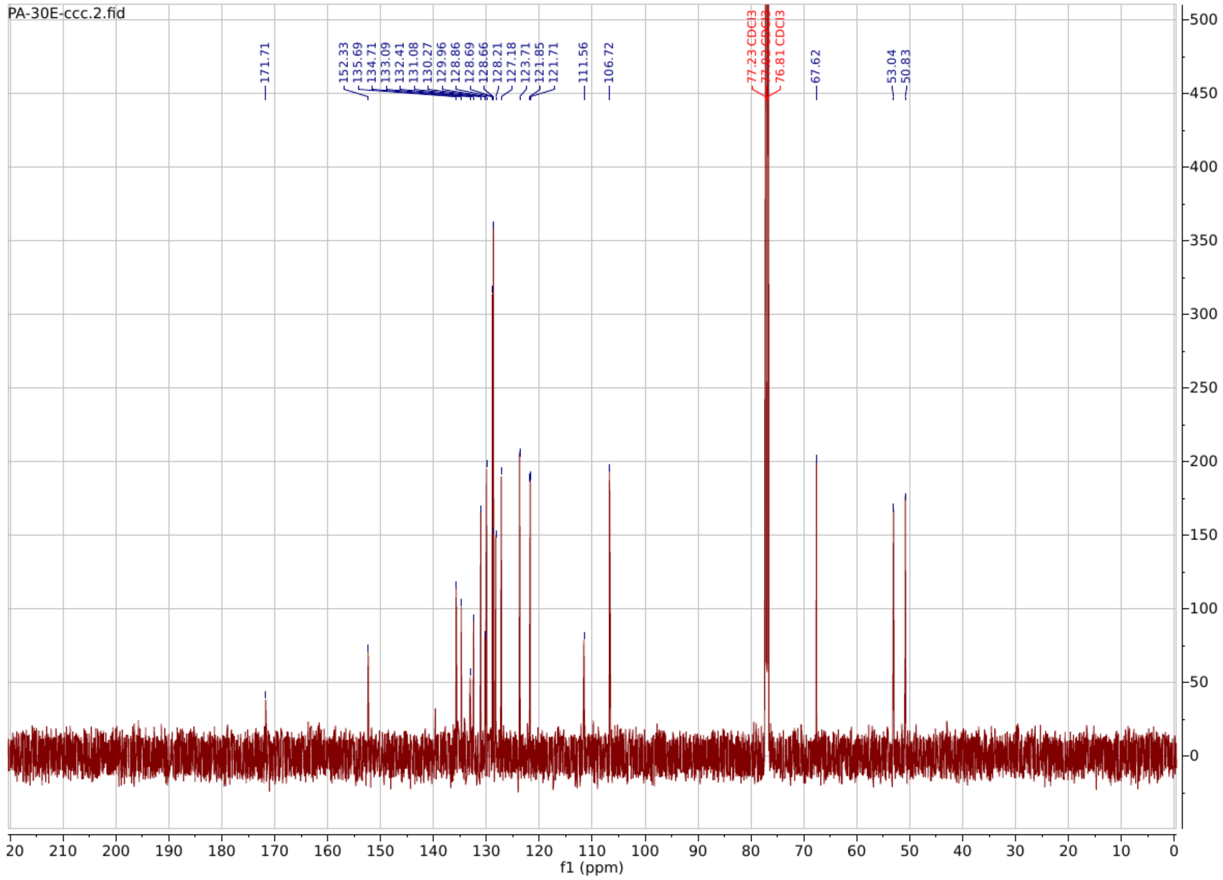

**PA146**

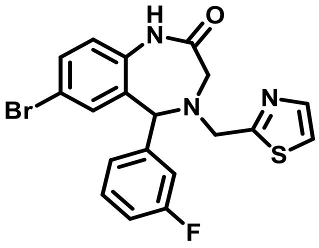

**7-bromo-5-(3-fluorophenyl)-4-(thiazol-2-ylmethyl)-1,3,4,5-tetrahydro-2H-benzo[e][1,4]diazepin-2-one**

$^{1}$H NMR (400 MHz, $CDCl_3$) δ 8.77 (s, 1H), 7.75 (d, *J* = 3.3 Hz, 1H), 7.42 (dd, *J* = 8.4, 2.3 Hz, 1H), 7.34 (dd, *J* = 7.6, 2.7 Hz, 2H), 7.20 – 7.12 (m, 2H), 7.06 – 6.95 (m, 3H), 5.08 (s, 1H), 4.35 – 4.10 (m, 2H), 3.51 (ddd, *J* = 78.8, 15.3, 1.1 Hz, 2H).

**$^{13}$C NMR** (151 MHz, $CDCl_3$) δ 171.82, 169.44, 142.77, 142.40, 136.15, 133.87, 132.20, 131.93, 130.49, 124.06, 122.67, 119.80, 119.23, 117.99, 115.45, 115.31, 77.25, 77.04, 76.83, 67.53, 61.99, 55.38, 53.66.

**HRMS (ESI):** m/z calculated for $C_{19}H_{15}BrFN_3OS$ $[M+H]^+$: 431.0103, found: 432.0193.

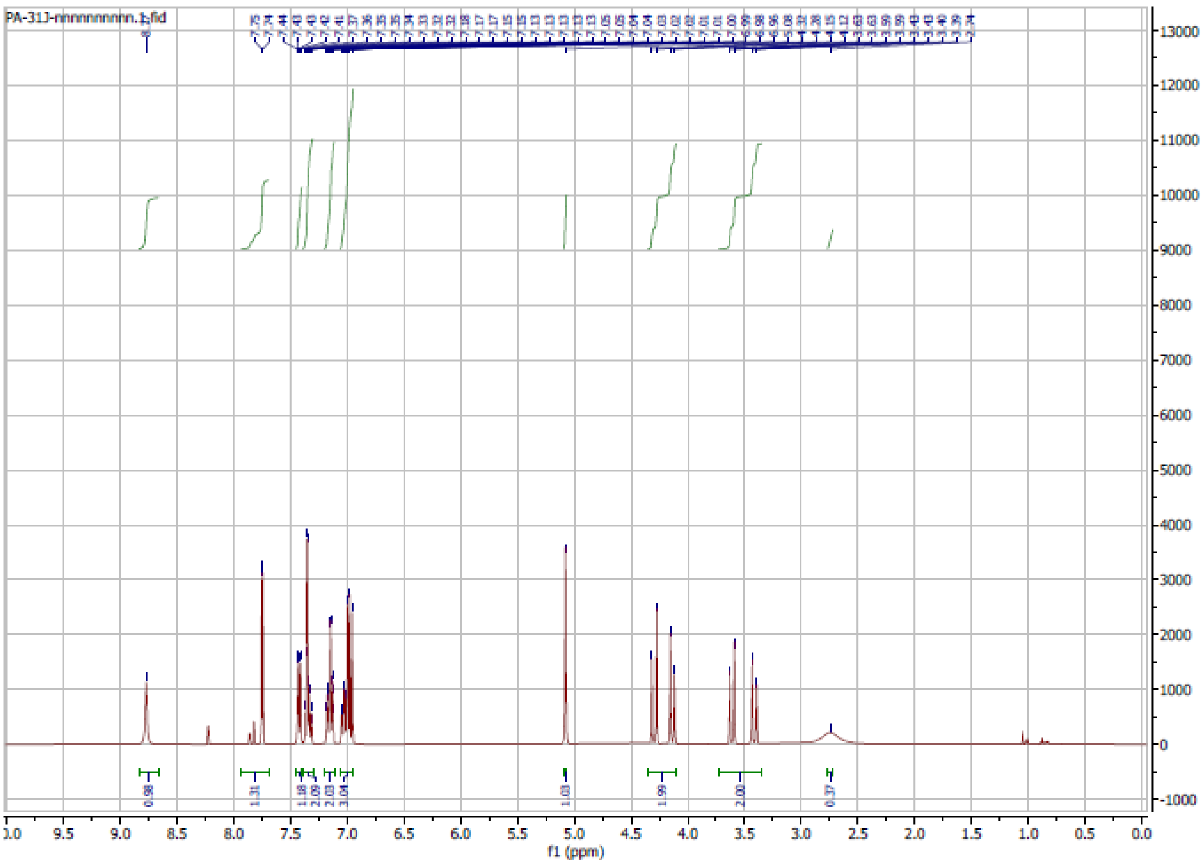

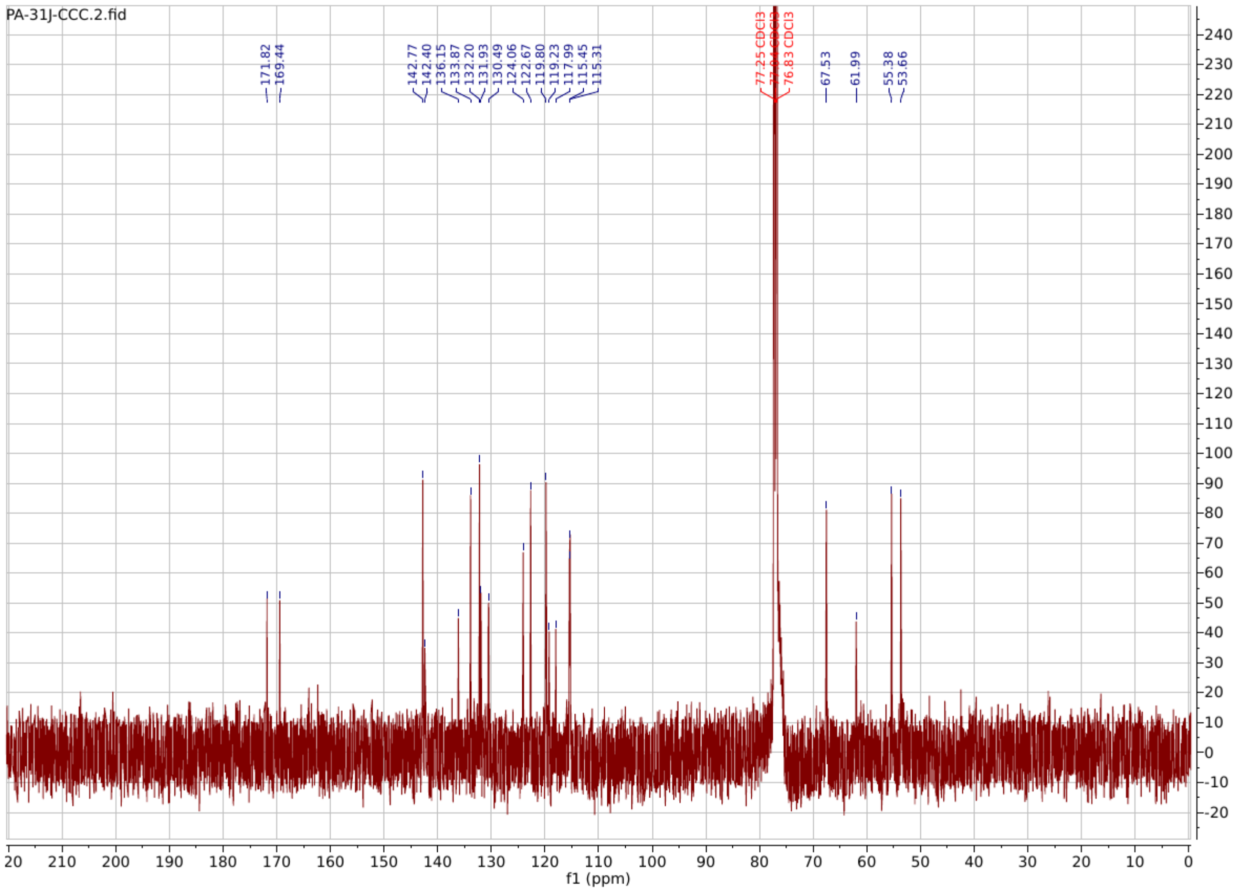

**PA149**

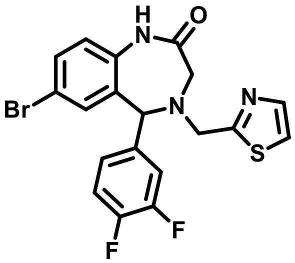

**7-bromo-5-(3,4-difluorophenyl)-4-(thiazol-2-ylmethyl)-1,3,4,5-tetrahydro-2H-benzo[e][1,4]diazepin-2-one**

$^{1}$**H NMR** (600 MHz, Chloroform-*d*) δ 7.88 – 7.70 (m, 1H), 7.55 – 6.81 (m, 8H), 5.05 (q, *J* = 20.0 Hz, 1H), 4.42 – 4.03 (m, 2H), 3.70 – 3.35 (m, 2H).

**$^{13}$C NMR** (151 MHz, $CDCl_3$) δ 173.38, 136.70, 136.53, 135.95, 133.88, 132.39, 131.75, 125.43, 124.47, 122.68, 119.99, 118.25, 117.80, 117.37, 117.25, 77.23, 77.02, 76.80, 69.40, 67.03, 55.20, 53.64.

**HRMS (ESI):** m/z calculated for $C_{19}H_{14}BrF_2N_3OS$ $[M+H]^+$: 449.0009, found: 450.0123.

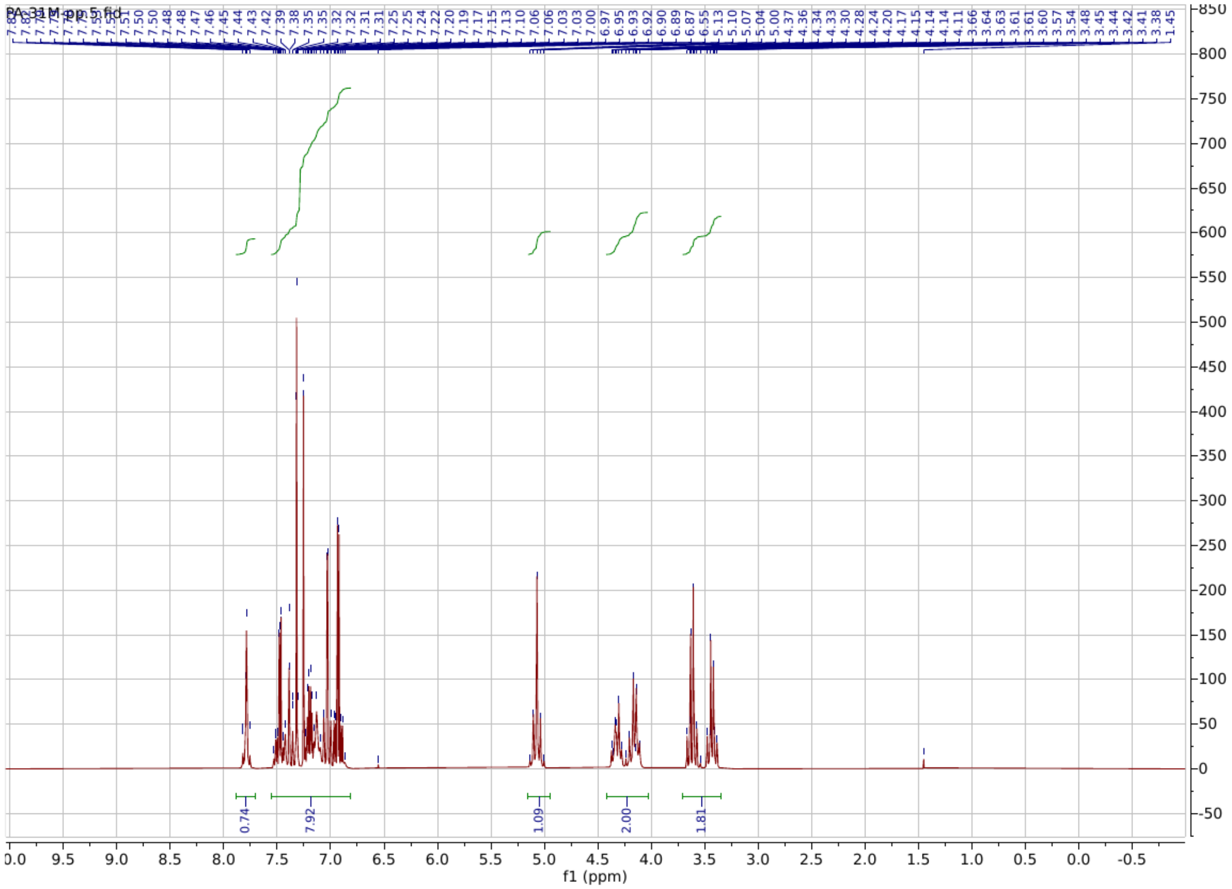

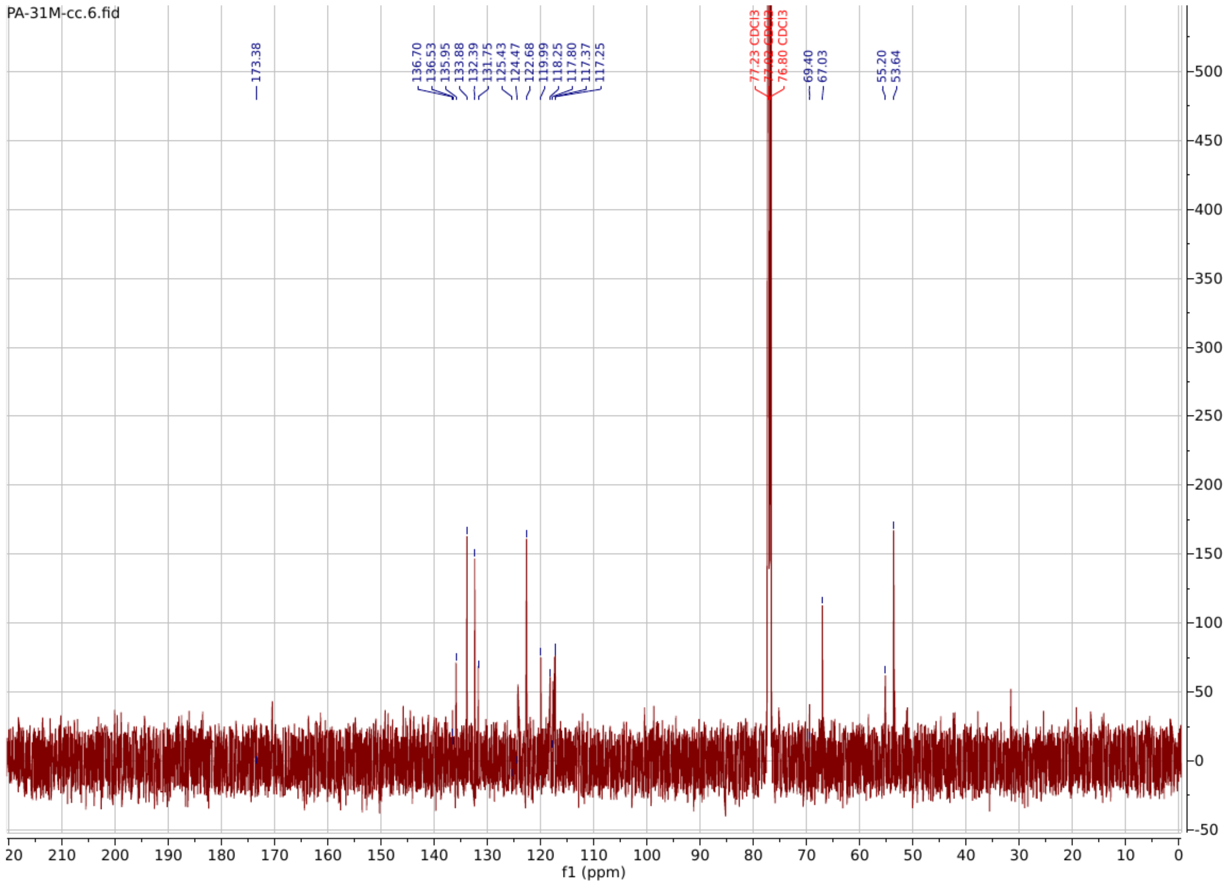

**PA151**

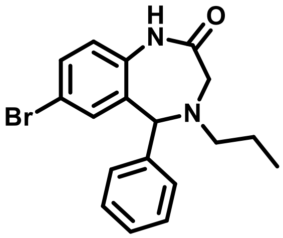

**7-bromo-5-phenyl-4-propyl-1,3,4,5-tetrahydro-2H-benzo[e][1,4]diazepin-2-one**

**$^{1}$H NMR** (600 MHz, Chloroform-*d*) δ 8.47 (s, 1H), 7.49 – 7.10 (m, 7H), 7.12 – 6.85 (m, 2H), 4.98 (s, 1H), 3.57 – 3.35 (m, 2H), 2.77 – 2.56 (m, 2H), 1.74 – 1.49 (m, 2H), 0.93 (q, *J* = 8.0, 7.2 Hz, 3H).

**$^{13}$C NMR** (151 MHz, $CDCl_3$) δ 173.58, 140.58, 136.12, 133.97, 133.19, 131.39, 128.68, 128.62, 127.84, 121.97, 117.21, 77.24, 77.02, 76.81, 68.53, 55.41, 52.78, 20.81, 11.54.

**HRMS (ESI):** m/z calculated for $C_{18}H_{19}BrN_2O$ $[M+H]^+$: 358.0681, found: 359.0738.

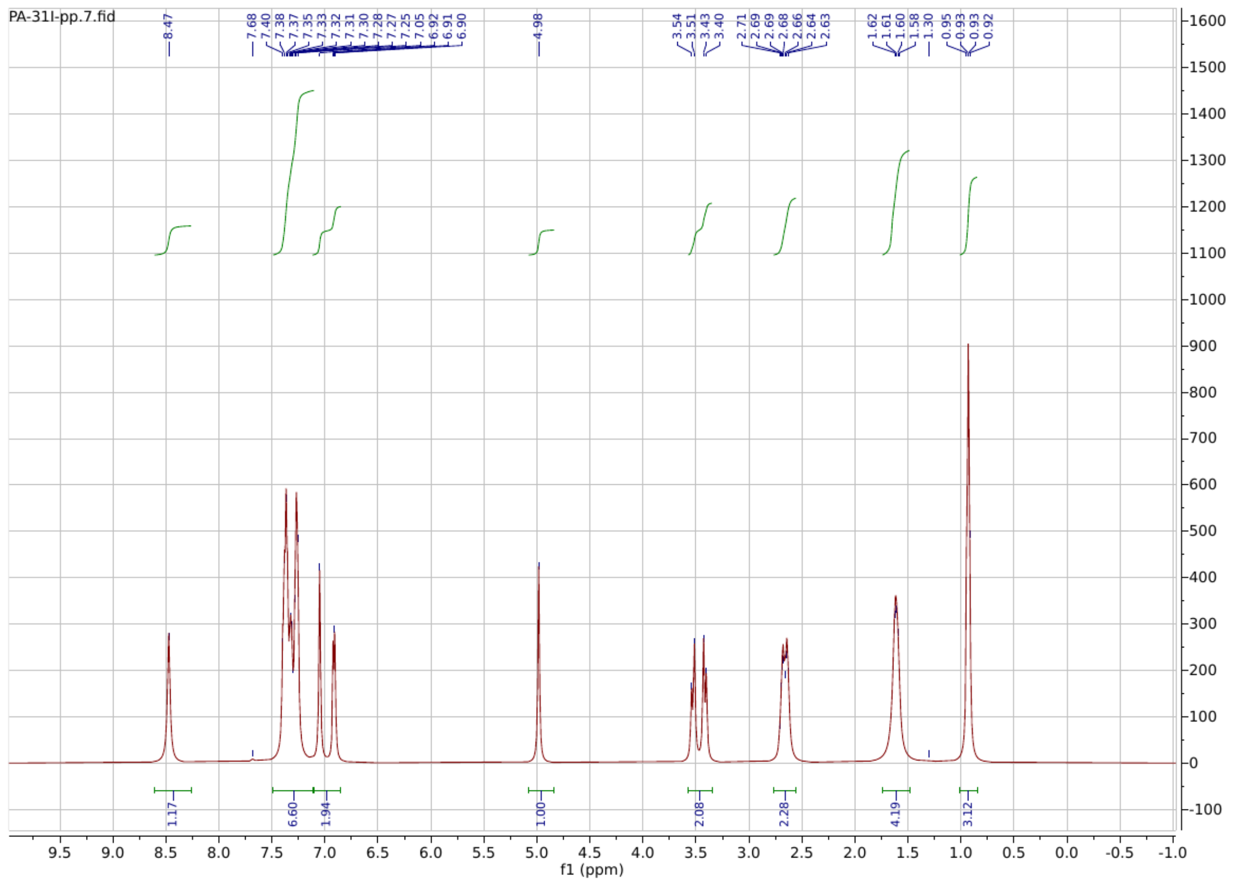

PA-31I-pp.7.fid
f1 (ppm)

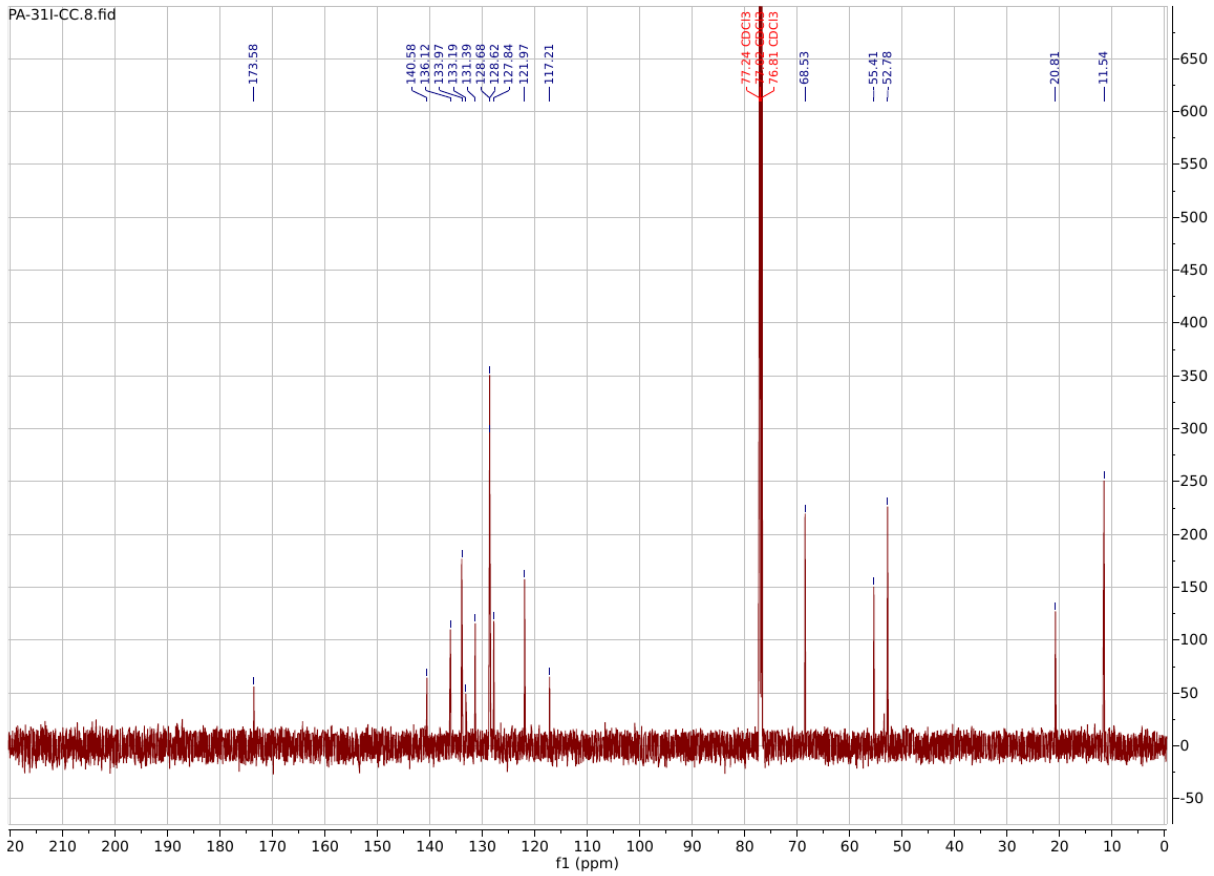

**PA156**

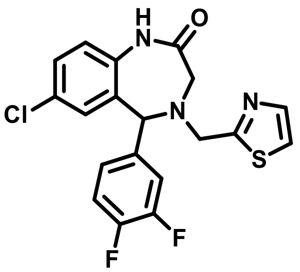

**7-chloro-5-(3,4-difluorophenyl)-4-(thiazol-2-ylmethyl)-1,3,4,5-tetrahydro-2H-benzo[e][1,4]diazepin-2-one**

$^{1}$**H NMR** (600 MHz, Chloroform-*d*) δ 8.17 (s, 1H), 7.91 – 7.67 (m, 1H), 7.57 – 7.20 (m, 9H), 7.00 (dd, *J* = 9.4, 4.5 Hz, 1H), 6.87 (s, 1H), 5.08 (s, 1H), 4.30 (d, *J* = 15.6 Hz, 1H), 4.14 (d, *J* = 15.3 Hz, 1H), 3.73 – 3.54 (m, 1H), 3.54 – 3.36 (m, 1H).

**$^{13}C$ NMR** (151 MHz, $CDCl_3$) δ 170.59, 169.44, 142.30, 139.29, 136.68, 135.53, 131.54, 130.95, 130.68, 129.42, 124.31, 122.45, 119.96, 117.83, 117.72, 117.39, 117.27, 77.23, 77.02, 76.81, 67.06, 55.22, 53.61, 53.43.

**HRMS (ESI):** m/z calculated for $C_{19}H_{14}ClF_2N_3OS$ $[M+H]^+$: 405.0514, found: 406.0577.

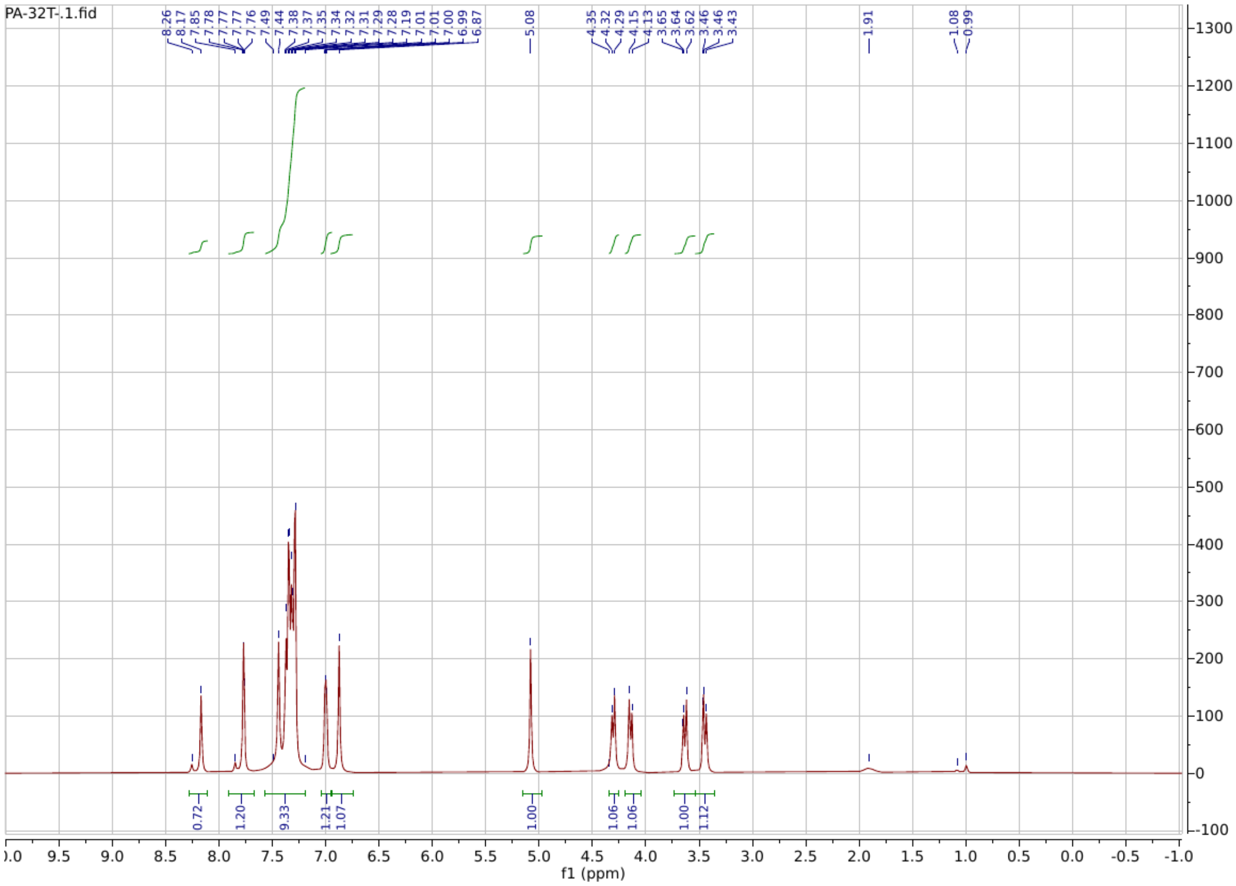

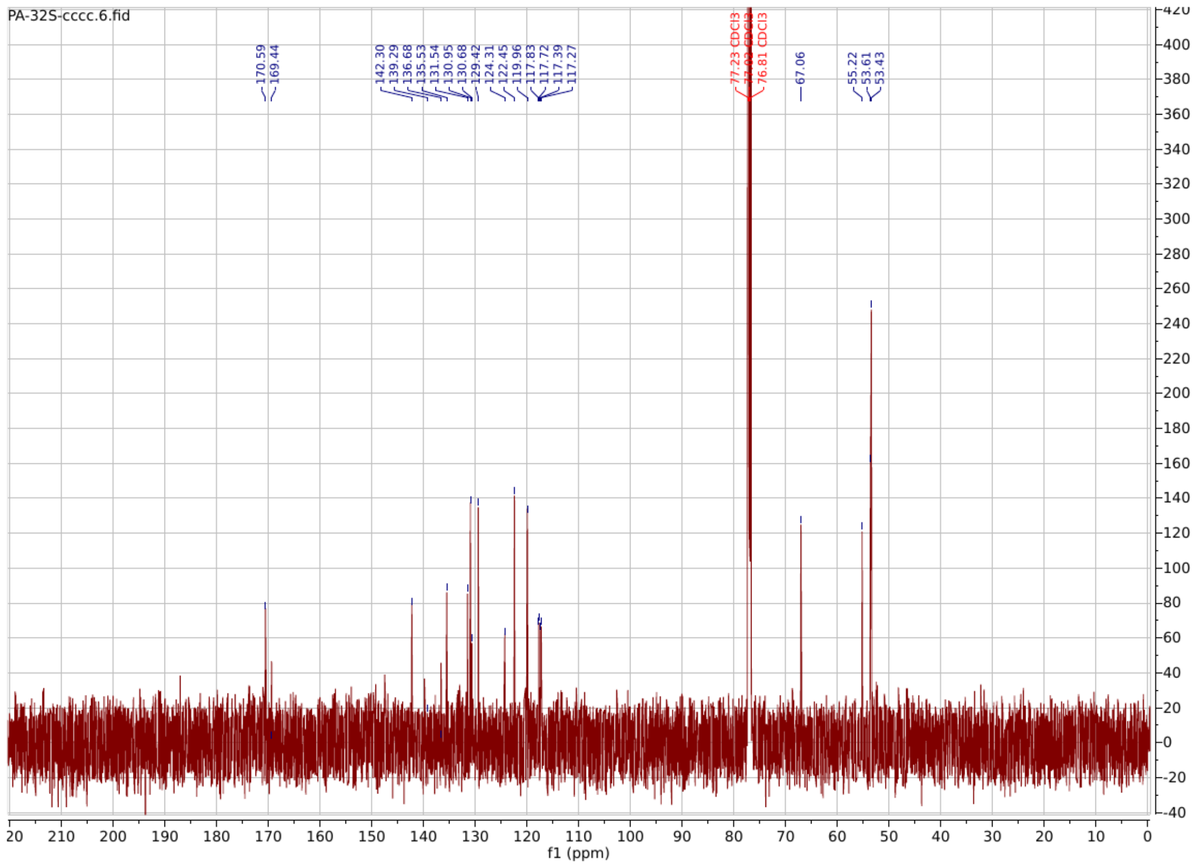

**PA157**

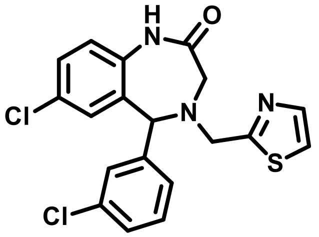

**7-chloro-5-(3-chlorophenyl)-4-(thiazol-2-ylmethyl)-1,3,4,5-tetrahydro-2H-benzo[e][1,4]diazepin-2-one**

**$^{1}$H NMR** (600 MHz, Chloroform-*d*) δ 8.17 (s, 1H), 7.86 – 7.70 (m, 1H), 7.52 – 7.17 (m, 9H), 7.00 (dd, *J* = 9.4, 4.5 Hz, 1H), 6.87 (s, 1H), 5.08 (s, 1H), 4.30 (d, *J* = 15.6 Hz, 1H), 4.14 (d, *J* = 15.3 Hz, 1H), 3.67 – 3.58 (m, 1H), 3.49 – 3.36 (m, 1H).

**$^{13}$C NMR** (151 MHz, $CDCl_3$) δ 170.96, 169.39, 142.50, 141.68, 135.58, 135.00, 131.63, 131.04, 130.54, 130.24, 129.30, 128.64, 128.51, 126.63, 122.31, 119.89, 77.24, 77.02, 76.81, 67.54, 55.29, 53.59.

**HRMS (ESI):** m/z calculated for $C_{19}H_{15}Cl_2N_3OS$ $[M+H]^+$: 403.0313, found: 404.0378.

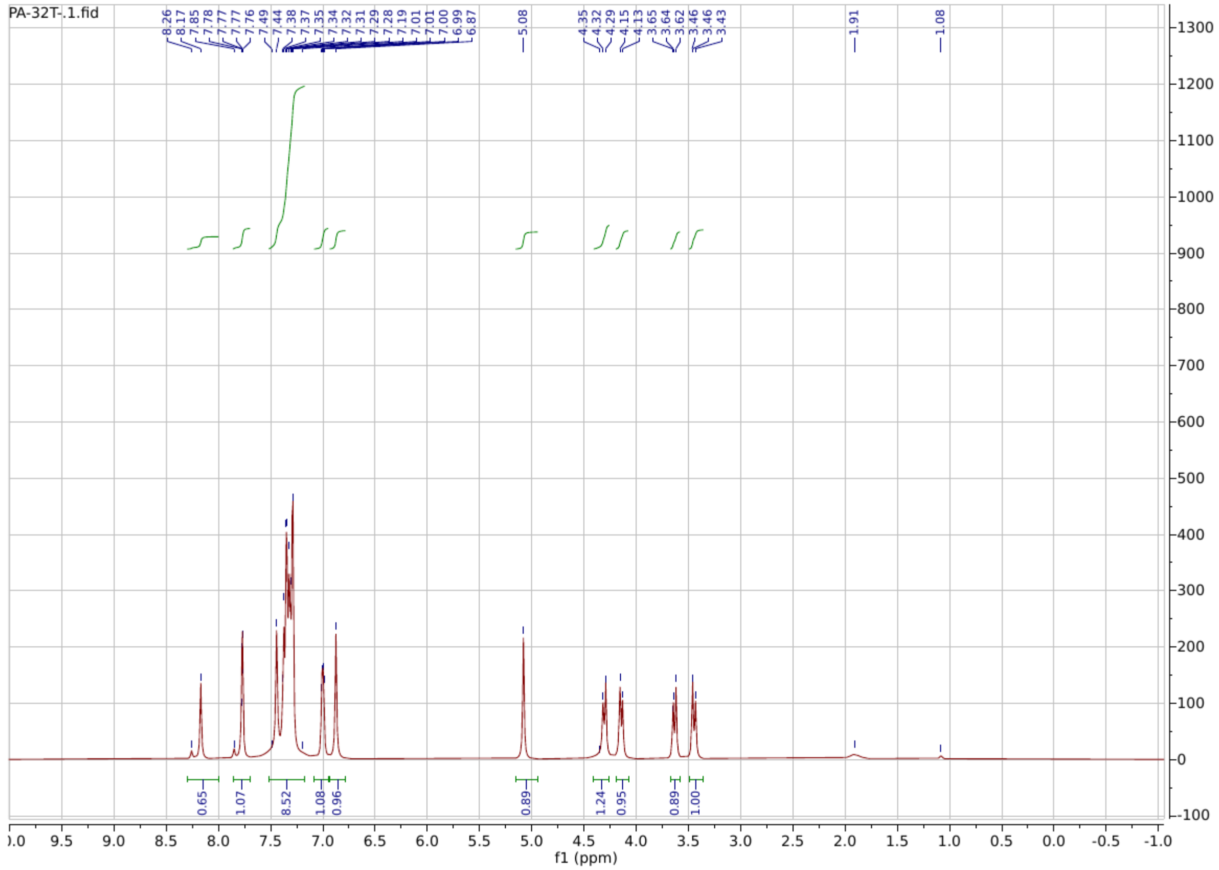

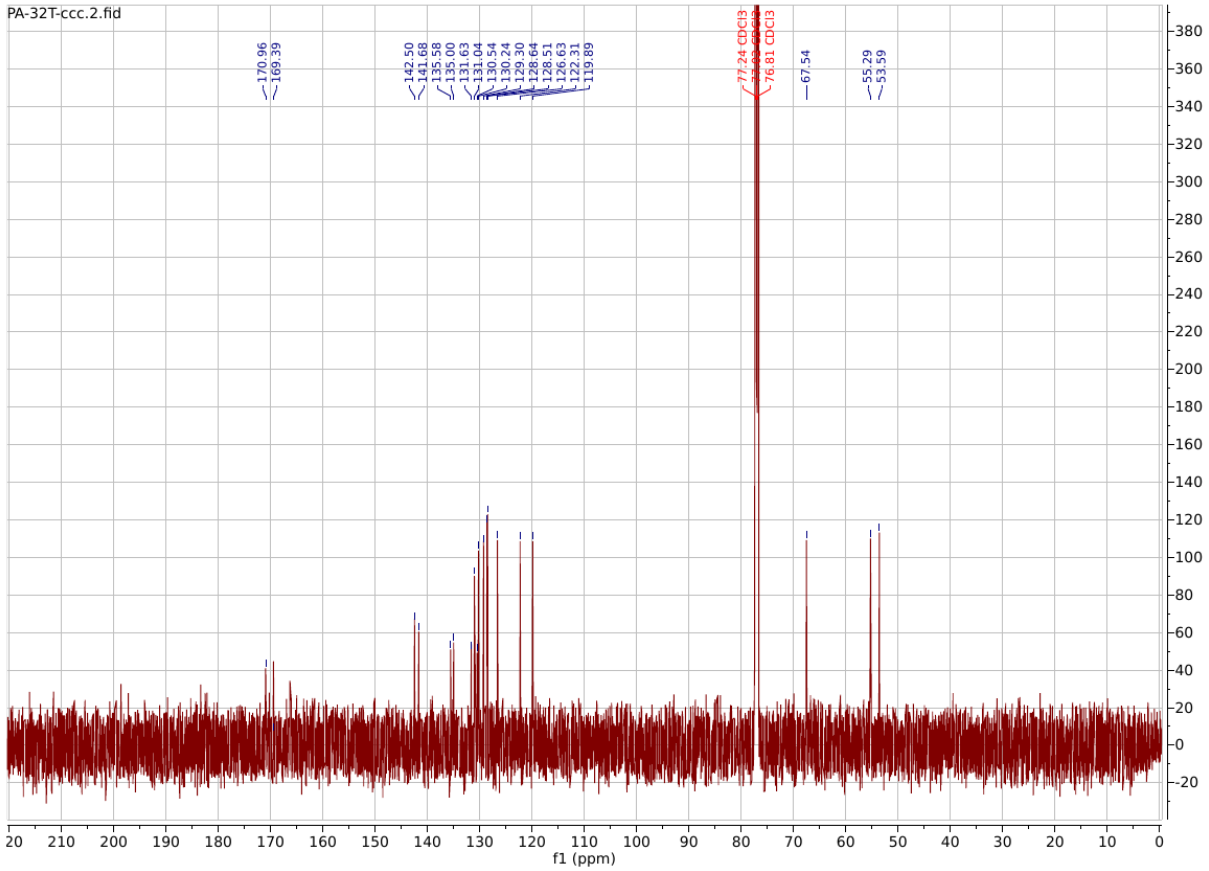

**PA158**

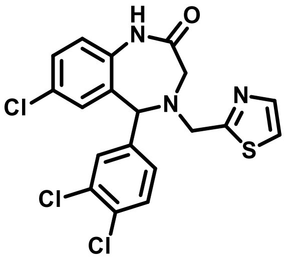

**7-chloro-5-(3,4-dichlorophenyl)-4-(thiazol-2-ylmethyl)-1,3,4,5-tetrahydro-2H-benzo[e][1,4]diazepin-2-one**

**$^{1}$H NMR** (600 MHz, Chloroform-*d*) δ 8.18 (d, *J* = 184.5 Hz, 1H), 7.97 – 7.71 (m, 2H), 7.60 – 7.43 (m, 3H), 7.35 (d, *J* = 6.2 Hz, 1H), 7.01 (ddd, *J* = 47.2, 9.6, 4.3 Hz, 1H), 6.86 (d, *J* = 9.9 Hz, 1H), 5.15 (d, *J* = 5.6 Hz, 1H), 4.65 – 4.23 (m, 1H), 4.12 (dd, *J* = 38.5, 15.2 Hz, 1H), 3.63 (dd, *J* = 54.8, 15.1 Hz, 1H), 3.35 (dd, *J* = 36.2, 13.6 Hz, 1H).

**$^{13}$C NMR** (151 MHz, $CDCl_3$) δ 175.28, 170.38, 169.38, 143.08, 142.38, 138.76, 135.61, 133.63, 133.27, 131.43, 131.30, 131.23, 130.93, 130.78, 130.22, 130.16, 130.00, 129.51, 127.64, 127.60, 122.92, 122.55, 120.53, 120.02, 119.88, 77.24, 77.02, 76.81, 68.07, 66.98, 60.69, 55.35, 54.09, 53.85, 53.62.

**HRMS (ESI):** m/z calculated for $C_{19}H_{14}Cl_3N_3OS$ $[M+H]^+$: 436.9923, found: 437.9984.

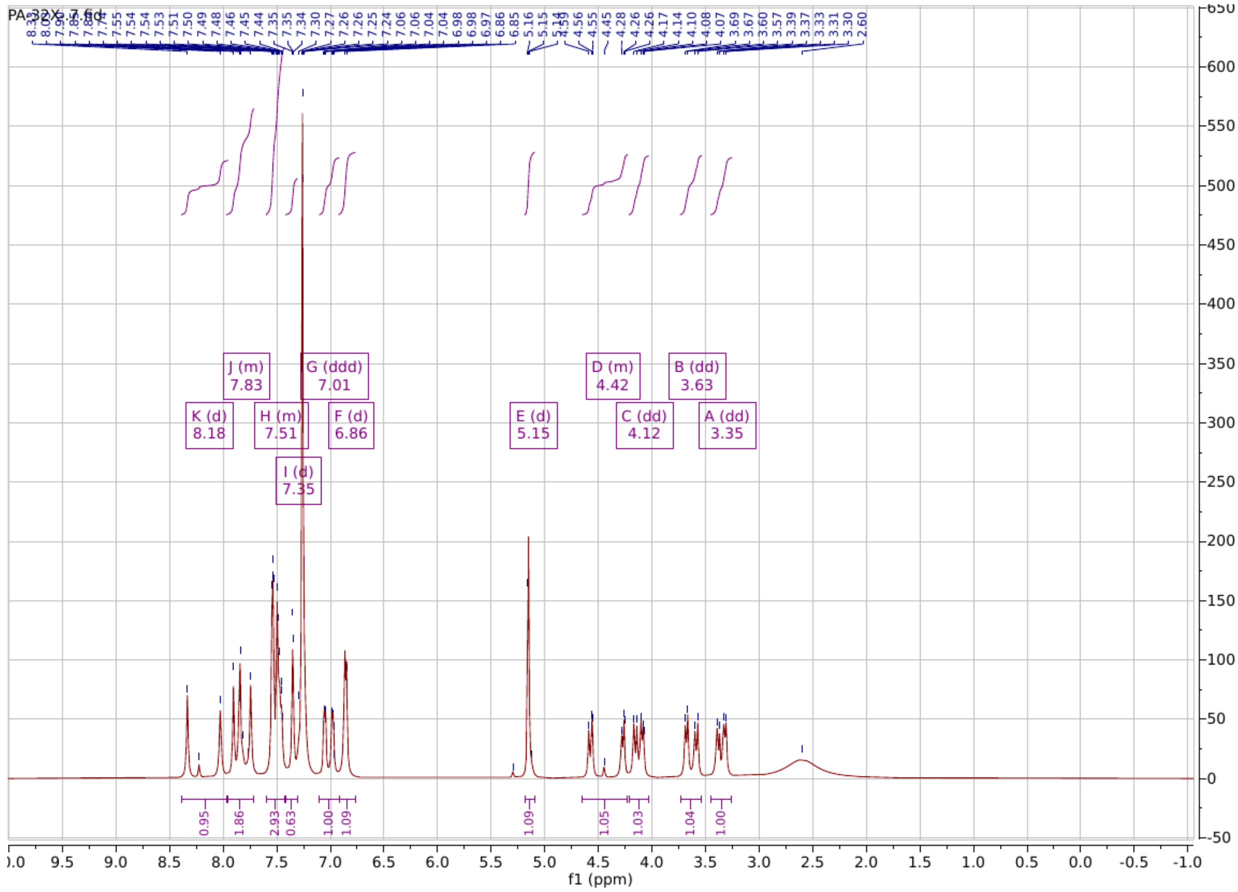

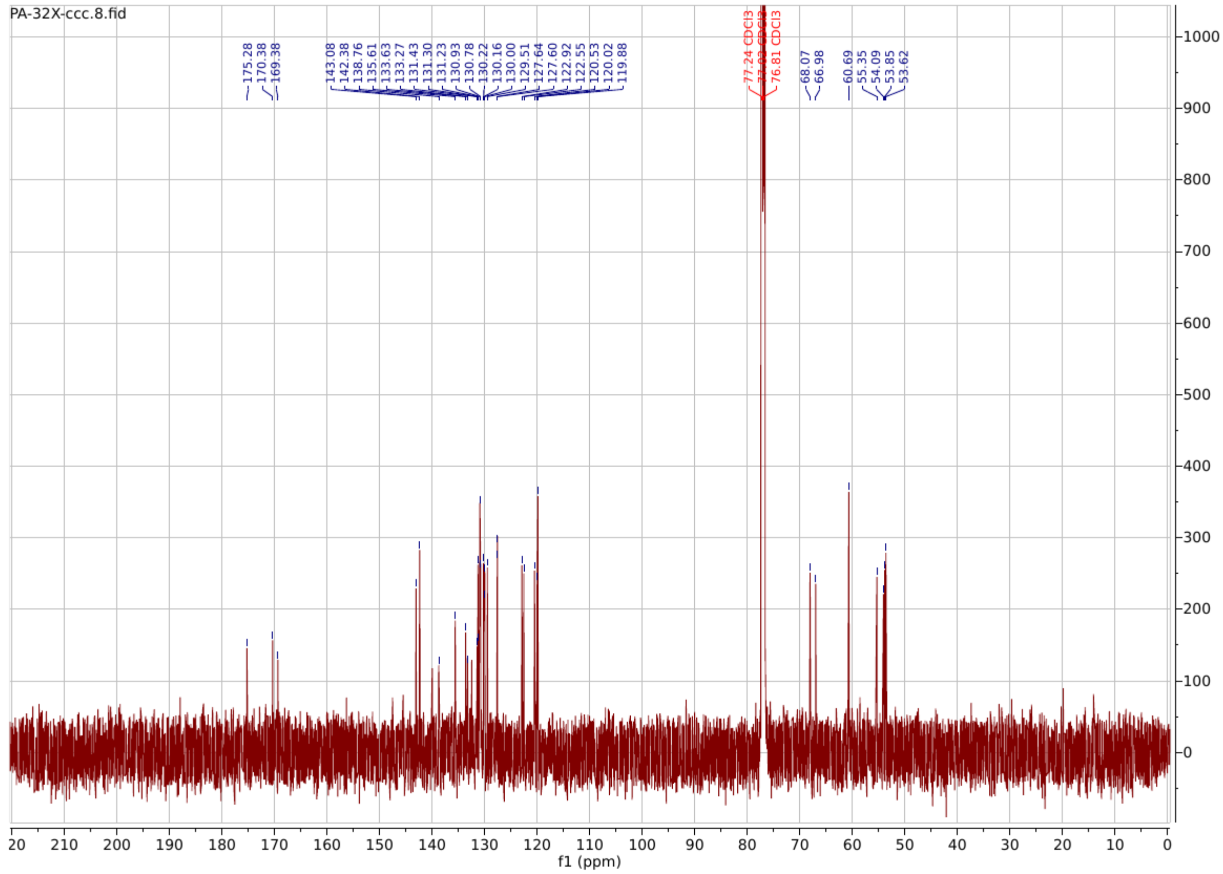

**PA159**

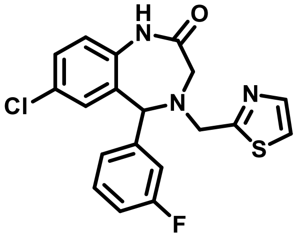

**7-chloro-5-(3-fluorophenyl)-4-(thiazol-2-ylmethyl)-1,3,4,5-tetrahydro-2H-benzo[e][1,4]diazepin-2-one**

**$^1$H NMR** (600 MHz, Chloroform-*d*) δ 8.17 (s, 1H), 7.85 (s, 1H), 7.49 – 7.31 (m, 5H), 7.10 (dt, *J* = 37.1, 6.4 Hz, 2H), 6.95 (s, 1H), 5.18 (s, 1H), 4.55 – 4.30 (m, 1H), 4.24 (d, *J* = 15.4 Hz, 1H), 3.71 (d, *J* = 15.1 Hz, 1H), 3.51 (d, *J* = 15.2 Hz, 1H).

**$^{13}$C NMR** (151 MHz, $CDCl_3$) δ 175.28, 170.38, 169.38, 143.08, 142.38, 138.76, 135.61, 133.63, 133.27, 131.43, 131.30, 131.23, 130.93, 130.78, 130.22, 130.16, 130.00, 129.51, 127.64, 127.60, 122.92, 122.55, 120.53, 120.02, 119.88, 77.24, 77.02, 76.81, 68.07, 66.98, 60.69, 55.35, 54.09, 53.85, 53.62.

**HRMS (ESI):** m/z calculated for $C_{19}H_{15}ClFN_3OS$ $[M+H]^+$: 387.0608, found: 388.0670.

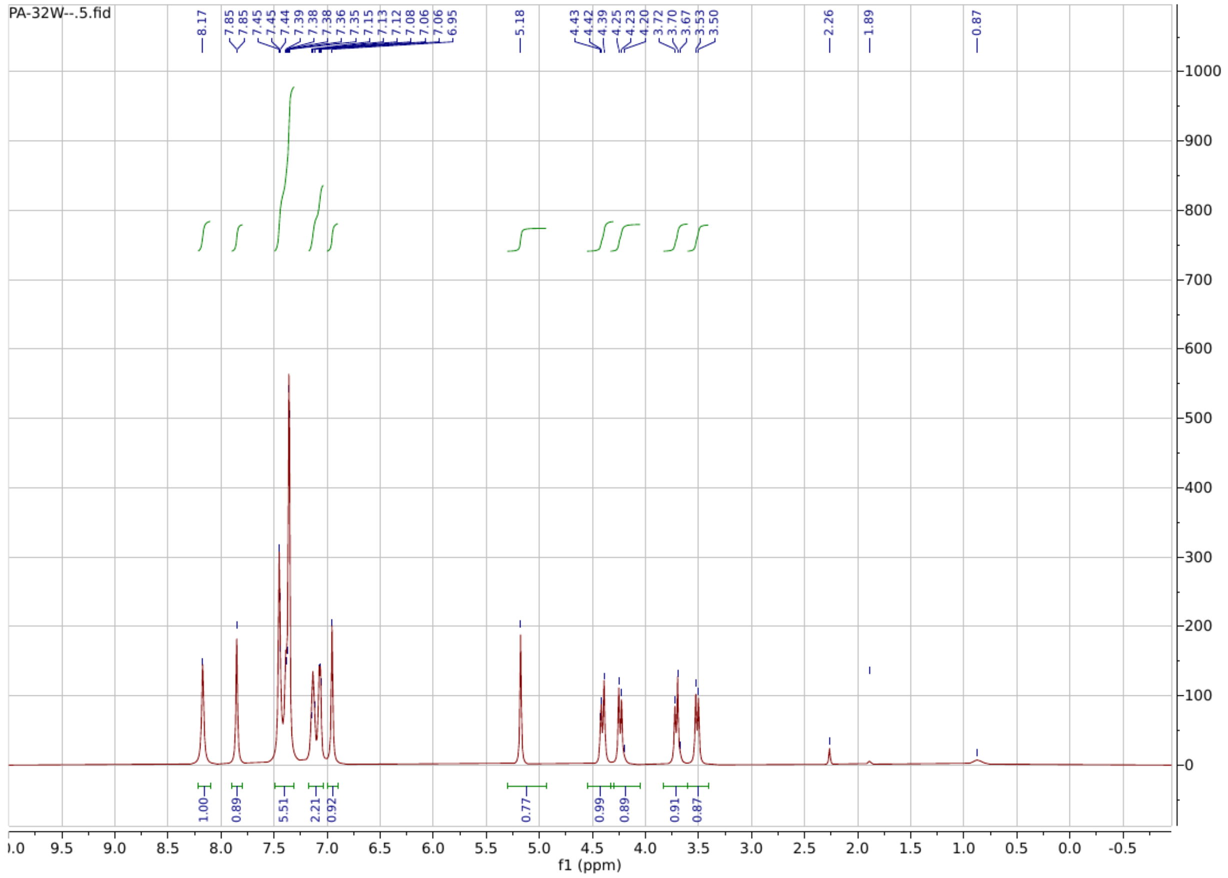

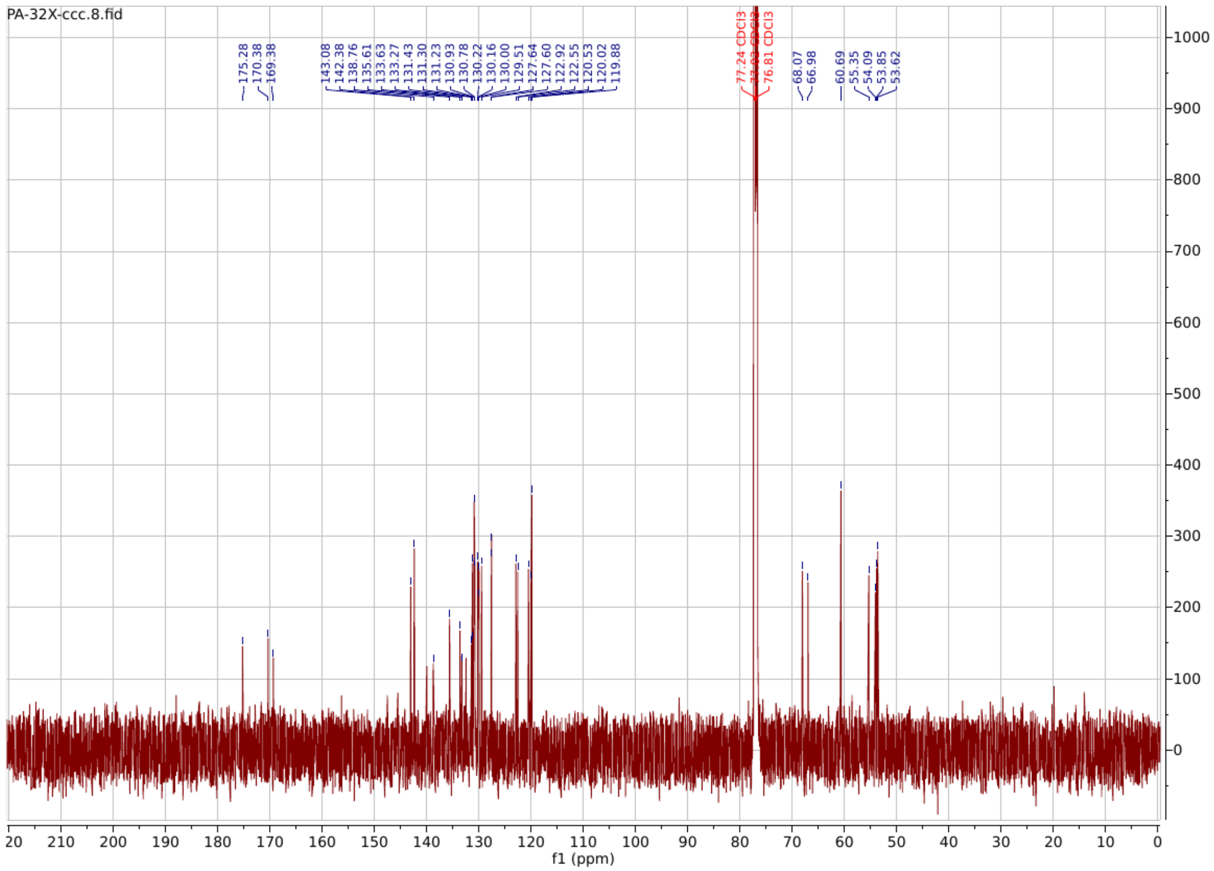

**PA160**

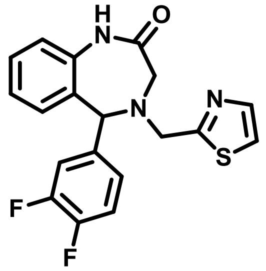

**5-(3,4-difluorophenyl)-4-(thiazol-2-ylmethyl)-1,3,4,5-tetrahydro-2H-benzo[e][1,4]diazepin-2-one**

**$^1$H NMR** (600 MHz, Chloroform-*d*) δ 8.17 (s, 1H), 7.76 (q, *J* = 3.5 Hz, 1H), 7.35 (tt, *J* = 11.0, 5.7 Hz, 3H), 7.22 – 6.98 (m, 4H), 6.96 – 6.83 (m, 1H), 5.08 (d, *J* = 4.6 Hz, 1H), 4.29 (dd, *J* = 15.5, 5.7 Hz, 1H), 4.15 (dd, *J* = 15.3, 5.6 Hz, 1H), 3.64 (dd, *J* = 14.9, 5.6 Hz, 1H), 3.43 (dd, *J* = 15.0, 5.3 Hz, 1H).

**$^{13}$C NMR** (151 MHz, $CDCl_3$) δ 170.99, 169.56, 151.88, 142.58, 137.54, 136.88, 131.25, 129.76, 129.35, 125.26, 124.32, 121.14, 119.74, 117.55, 117.40, 117.28, 77.24, 77.03, 76.81, 67.38, 55.44, 53.74.

**HRMS (ESI):** m/z calculated for $C_{19}H_{15}F_2N_3OS$ $[M+H]^+$: 371.0904, found: 372.0912.

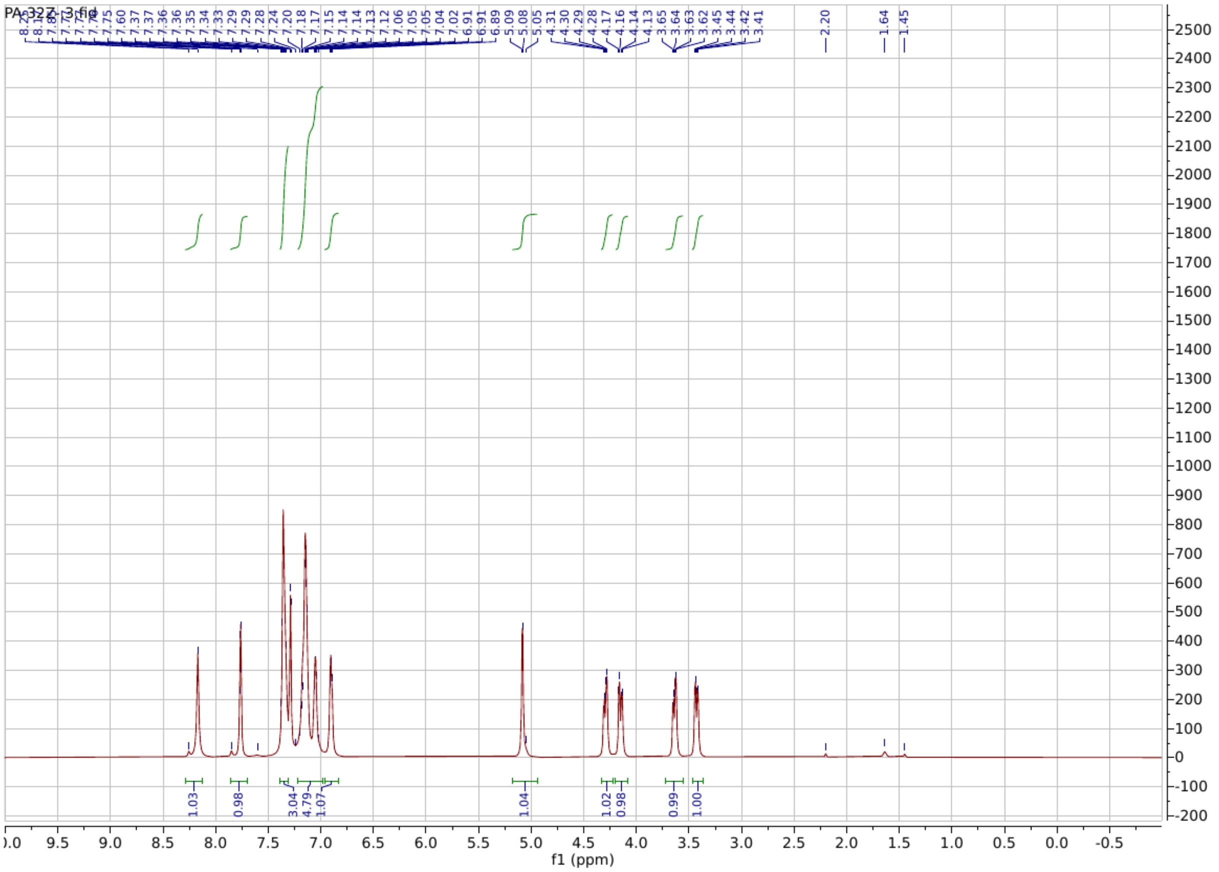

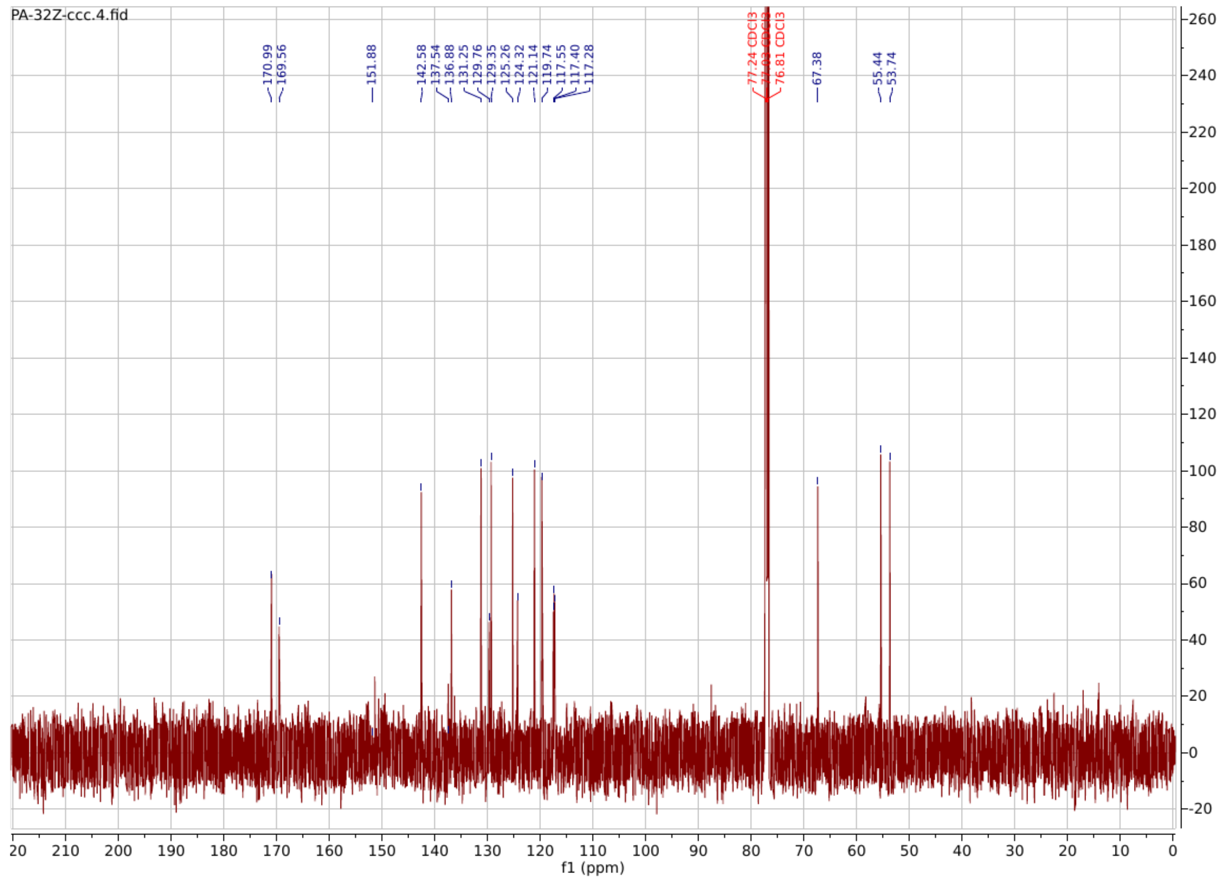

**PA162**

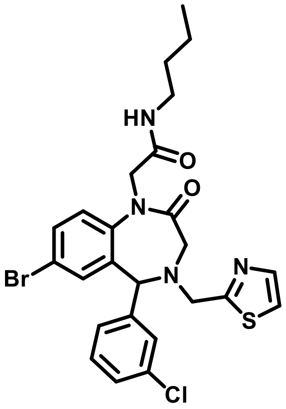

**2-(7-bromo-5-(3-chlorophenyl)-2-oxo-4-(thiazol-2-ylmethyl)-2,3,4,5-tetrahydro-1H-benzo[e][1,4]diazepin-1-yl)-N-butylacetamide**

**$^{1}$H NMR** (600 MHz, Chloroform-*d*) δ 7.75 (s, 1H), 7.59 – 7.38 (m, 3H), 7.31 (dd, *J* = 21.8, 7.3 Hz, 5H), 7.09 (s, 1H), 6.29 (s, 1H), 5.01 (s, 1H), 4.19 (d, *J* = 15.2 Hz, 1H), 4.11 – 3.96

(m, 2H), 3.57 (d, $J$ = 12.5 Hz, 1H), 3.36 – 3.19 (m, 4H), 1.43 (dt, $J$ = 88.5, 9.6 Hz, 4H), 1.08 – 0.81 (m, 3H).

**$^{13}$C NMR** (151 MHz, $CDCl_3$) δ 168.53, 168.04, 167.77, 142.64, 141.54, 134.75, 134.26, 133.24, 132.68, 129.98, 128.31, 127.66, 124.80, 120.36, 119.93, 77.24, 77.03, 76.81, 66.39, 56.06, 54.17, 52.28, 39.50, 31.50, 20.03, 13.71.

**HRMS (ESI):** m/z calculated for $C_{25}H_{26}BrClN_4O2S$ $[M+H]^+$: 560.0648, found: 561.0715.

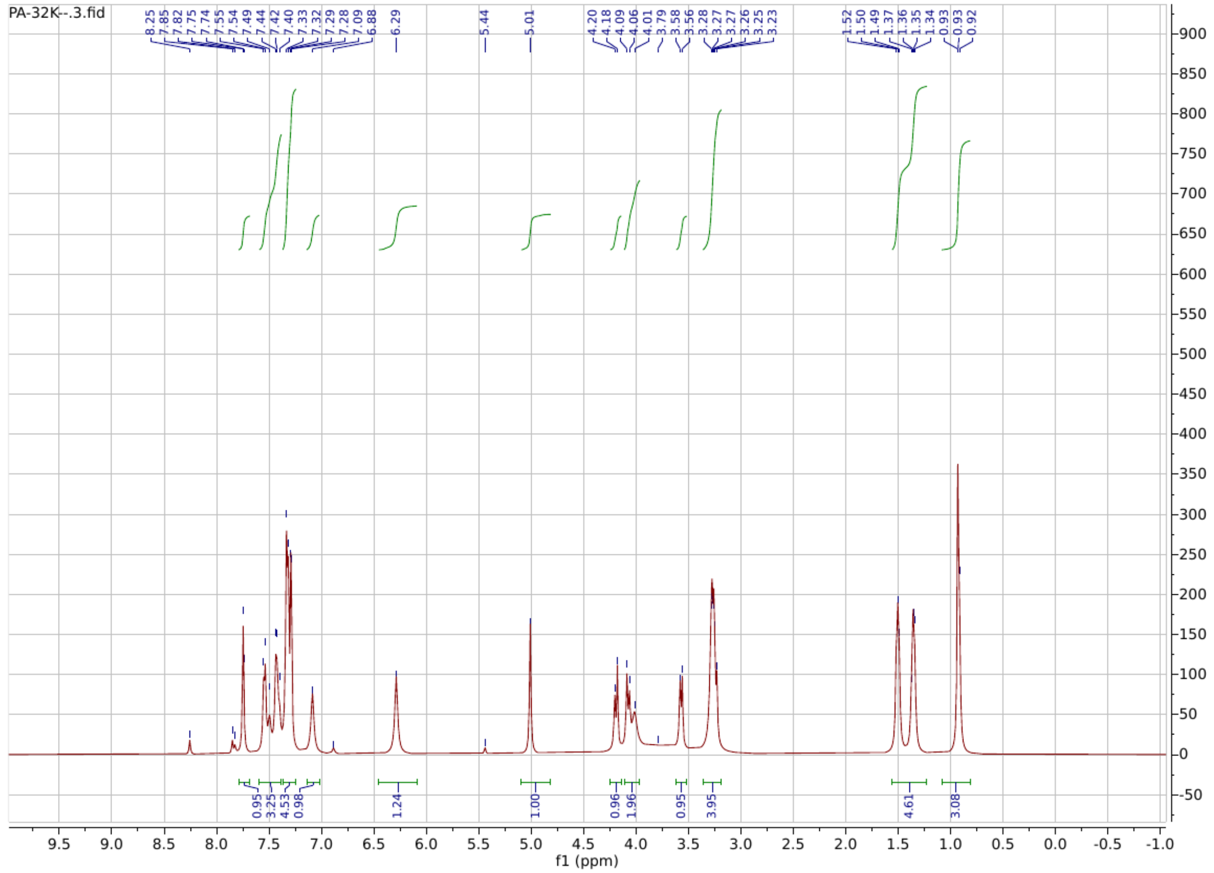

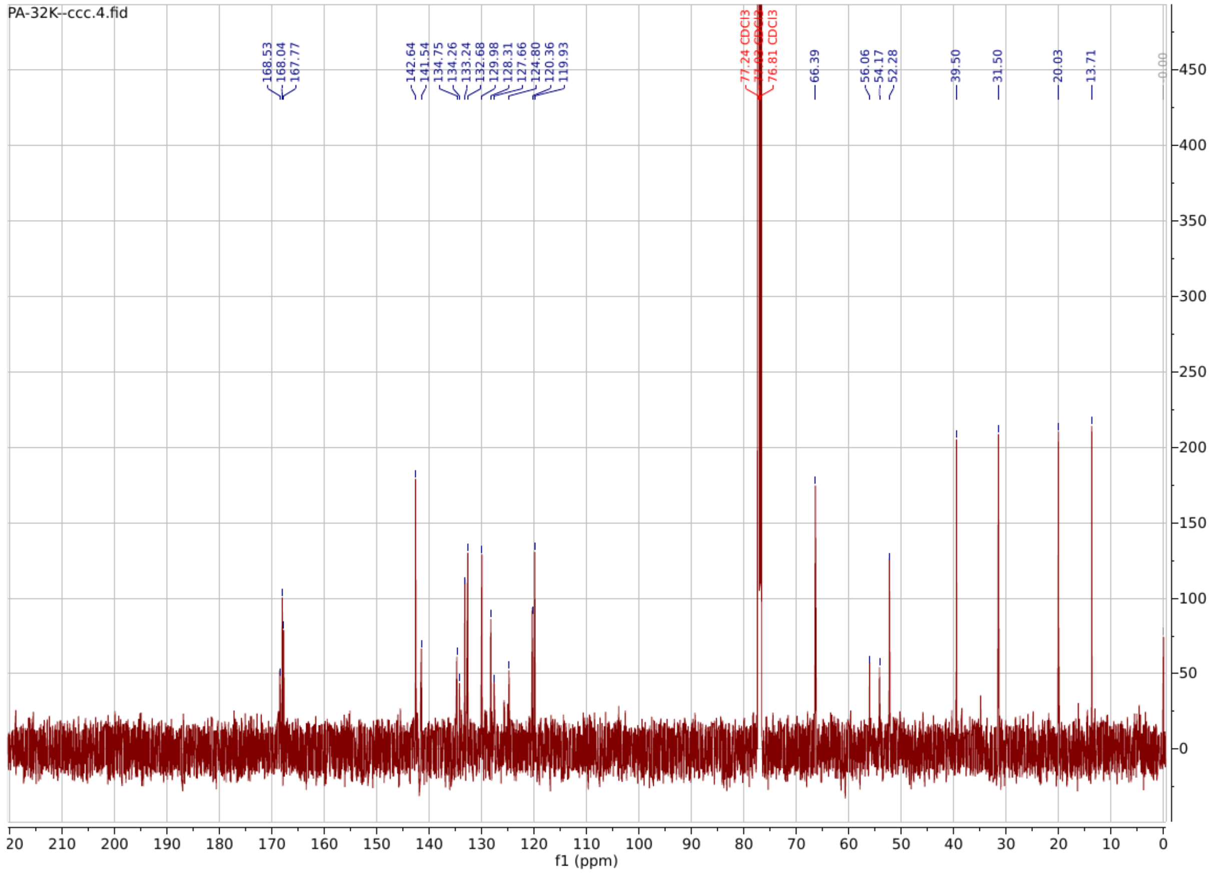
