## Supplemental Figures for "Small Molecules Restore Azole Activity Against Drug-Tolerant and Drug-Resistant *Candida* Isolates"

**Figure S1.**

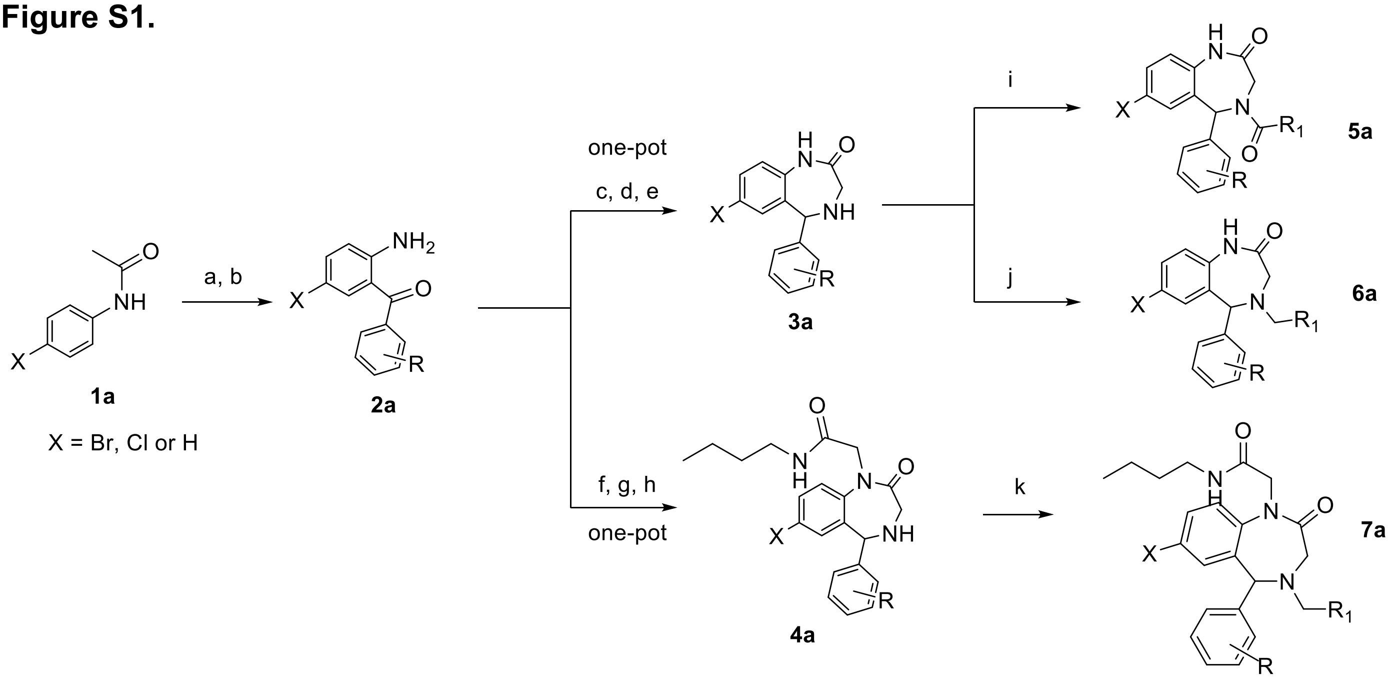

**Figure S2.**

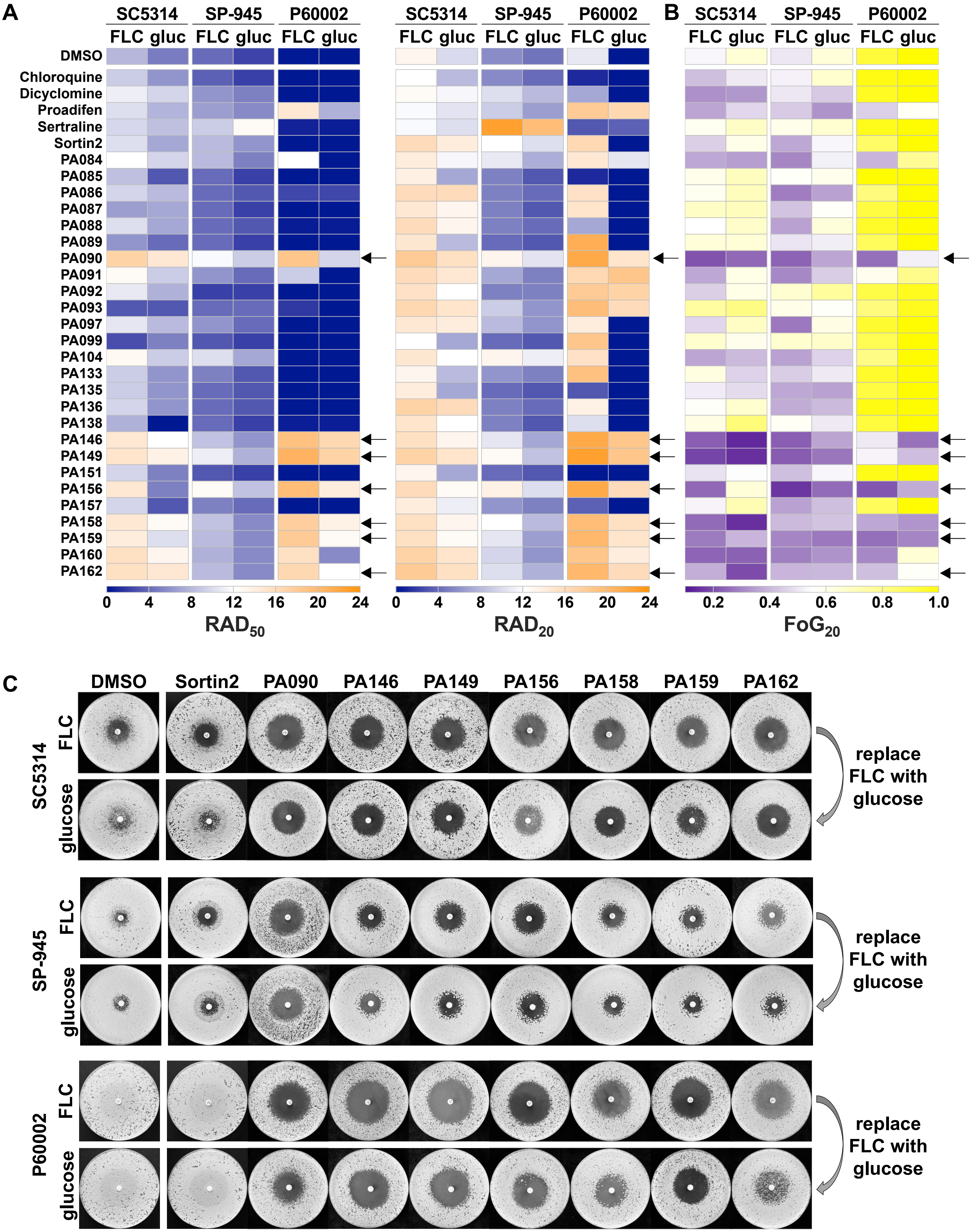
A
B
C
SC5314
SP-945
P60002
FLC gluc
DMSO
Chloroquine
Dicyclomine
Proadifen
Sertraline
Sortin2
PA084
PA085
PA086
PA087
PA088
PA089
PA090
PA091
PA092
PA093
PA097
PA099
PA104
PA133
PA135
PA136
PA138
PA146
PA149
PA151
PA156
PA157
PA158
PA159
PA160
PA162
0 4 8 12 16 20 24
RAD50
RAD20
0.2 0.4 0.6 0.8 1.0
FoG20
DMSO Sortin2 PA090 PA146 PA149 PA156 PA158 PA159 PA162
FLC
glucose
replace FLC with glucose

Figure S3.

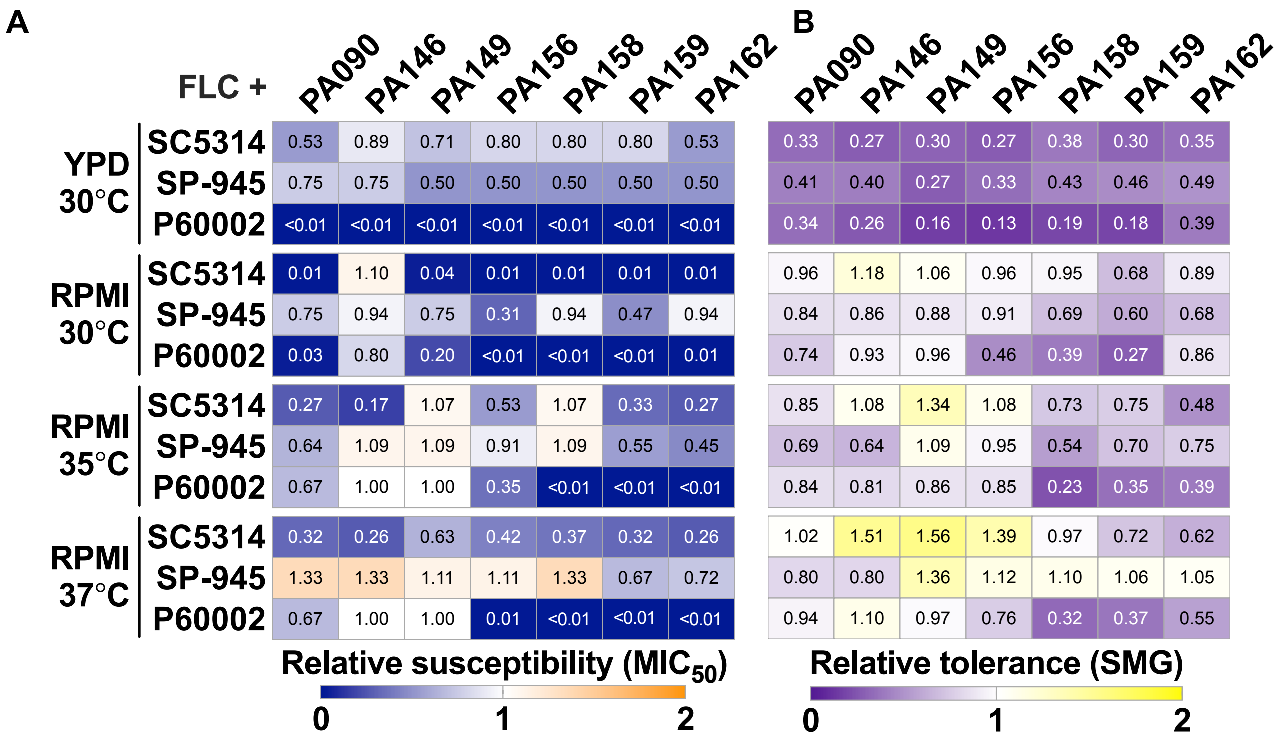
A
FLC +
PA090 PA146 PA149 PA156 PA158 PA159 PA162
YPD 30°C
SC5314 0.53 0.89 0.71 0.80 0.80 0.80 0.53
SP-945 0.75 0.75 0.50 0.50 0.50 0.50 0.50
P60002 <0.01 <0.01 <0.01 <0.01 <0.01 <0.01 <0.01
RPMI 30°C
SC5314 0.01 1.10 0.04 0.01 0.01 0.01 0.01
SP-945 0.75 0.94 0.75 0.31 0.94 0.47 0.94
P60002 0.03 0.80 0.20 <0.01 <0.01 <0.01 0.01
RPMI 35°C
SC5314 0.27 0.17 1.07 0.53 1.07 0.33 0.27
SP-945 0.64 1.09 1.09 0.91 1.09 0.55 0.45
P60002 0.67 1.00 1.00 0.35 <0.01 <0.01 <0.01
RPMI 37°C
SC5314 0.32 0.26 0.63 0.42 0.37 0.32 0.26
SP-945 1.33 1.33 1.11 1.11 1.33 0.67 0.72
P60002 0.67 1.00 1.00 0.01 <0.01 <0.01 <0.01
Relative susceptibility (MIC50)
0 1 2
B
PA090 PA146 PA149 PA156 PA158 PA159 PA162
0.33 0.27 0.30 0.27 0.38 0.30 0.35
0.41 0.40 0.27 0.33 0.43 0.46 0.49
0.34 0.26 0.16 0.13 0.19 0.18 0.39
0.96 1.18 1.06 0.96 0.95 0.68 0.89
0.84 0.86 0.88 0.91 0.69 0.60 0.68
0.74 0.93 0.96 0.46 0.39 0.27 0.86
0.85 1.08 1.34 1.08 0.73 0.75 0.48
0.69 0.64 1.09 0.95 0.54 0.70 0.75
0.84 0.81 0.86 0.85 0.23 0.35 0.39
1.02 1.51 1.56 1.39 0.97 0.72 0.62
0.80 0.80 1.36 1.12 1.10 1.06 1.05
0.94 1.10 0.97 0.76 0.32 0.37 0.55
Relative tolerance (SMG)
0 1 2

Figure S4.

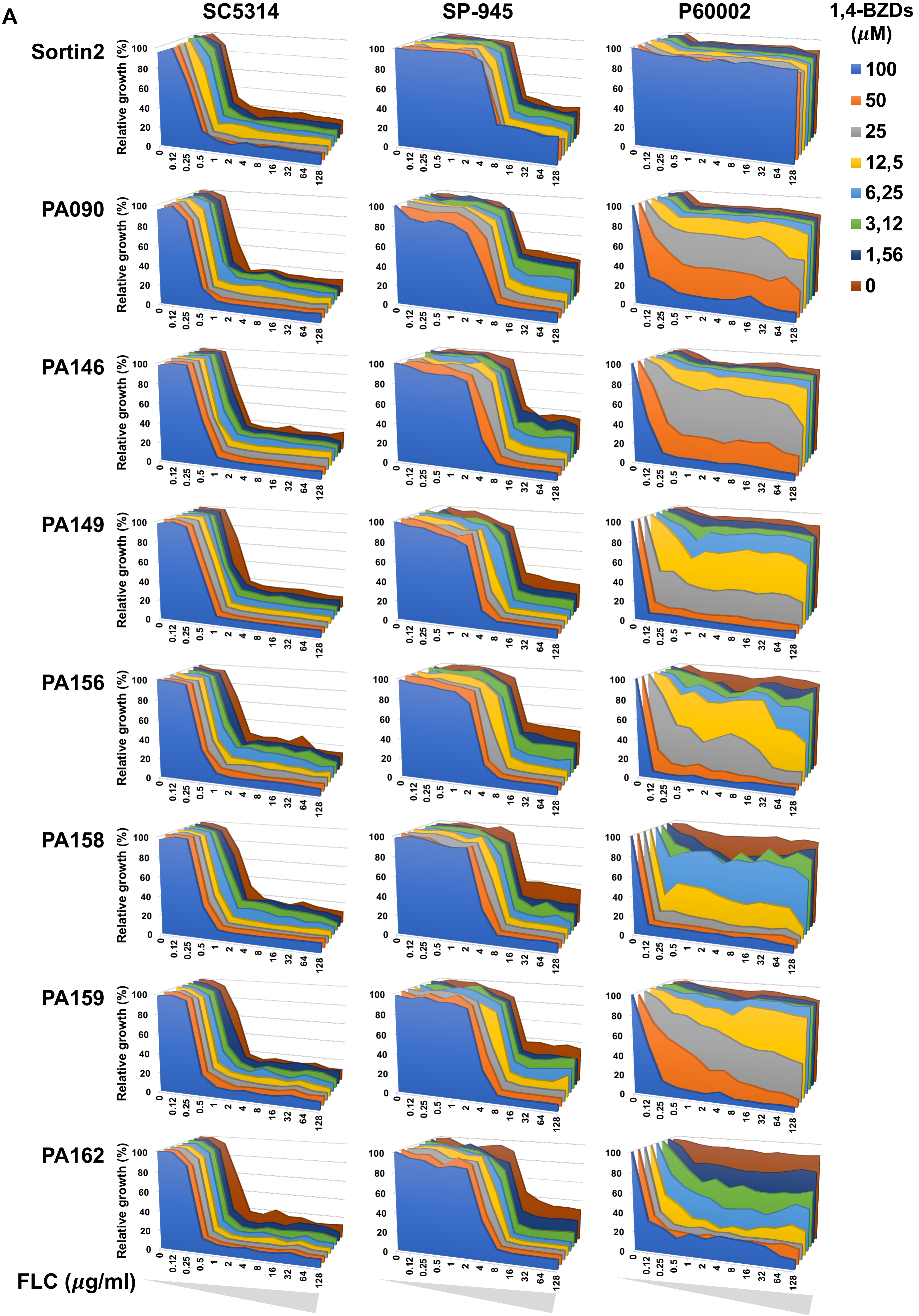
A
SC5314
SP-945
P60002
1,4-BZDs (μM)
100
50
25
12,5
6,25
3,12
1,56
0
Sortin2
PA090
PA146
PA149
PA156
PA158
PA159
PA162
Relative growth (%)
0
20
40
60
80
100
0
0.12
0.25
0.5
1
2
4
8
16
32
64
128
FLC (μg/ml)

**Figure S5.**

**A**

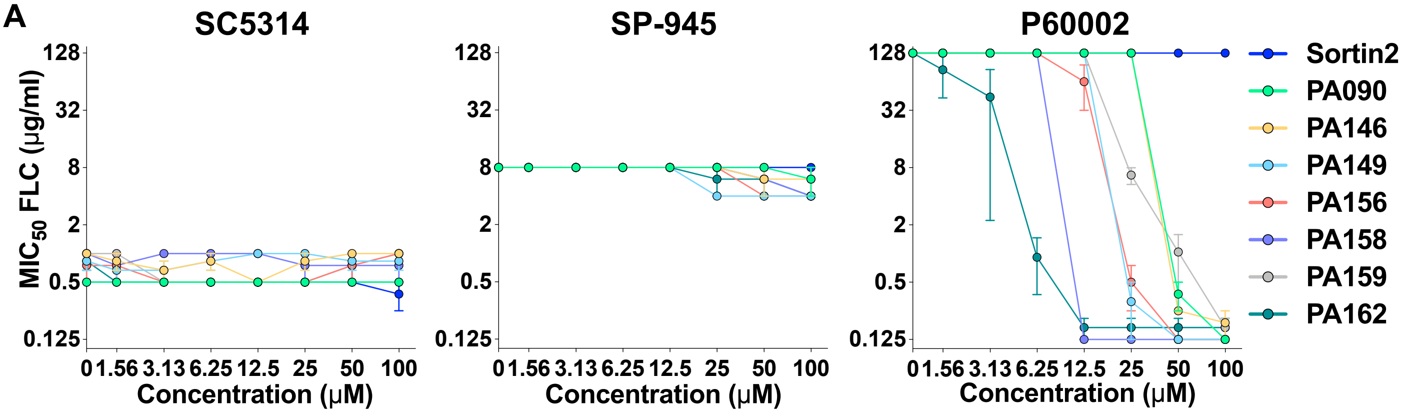


**B**

| Interaction | FICI |
|---|---|
| antagonistic | >4 |
| indifferent | 1-4 |
| additive | 0.5-1 |
| synergistic | <0.5 |

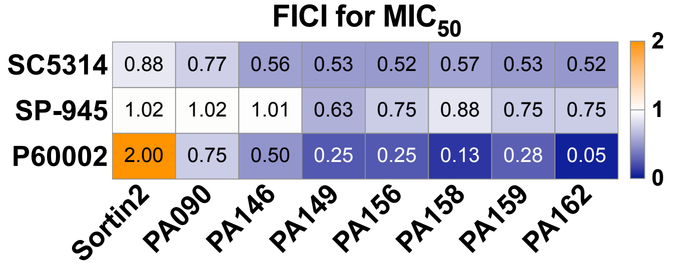

Figure S6.

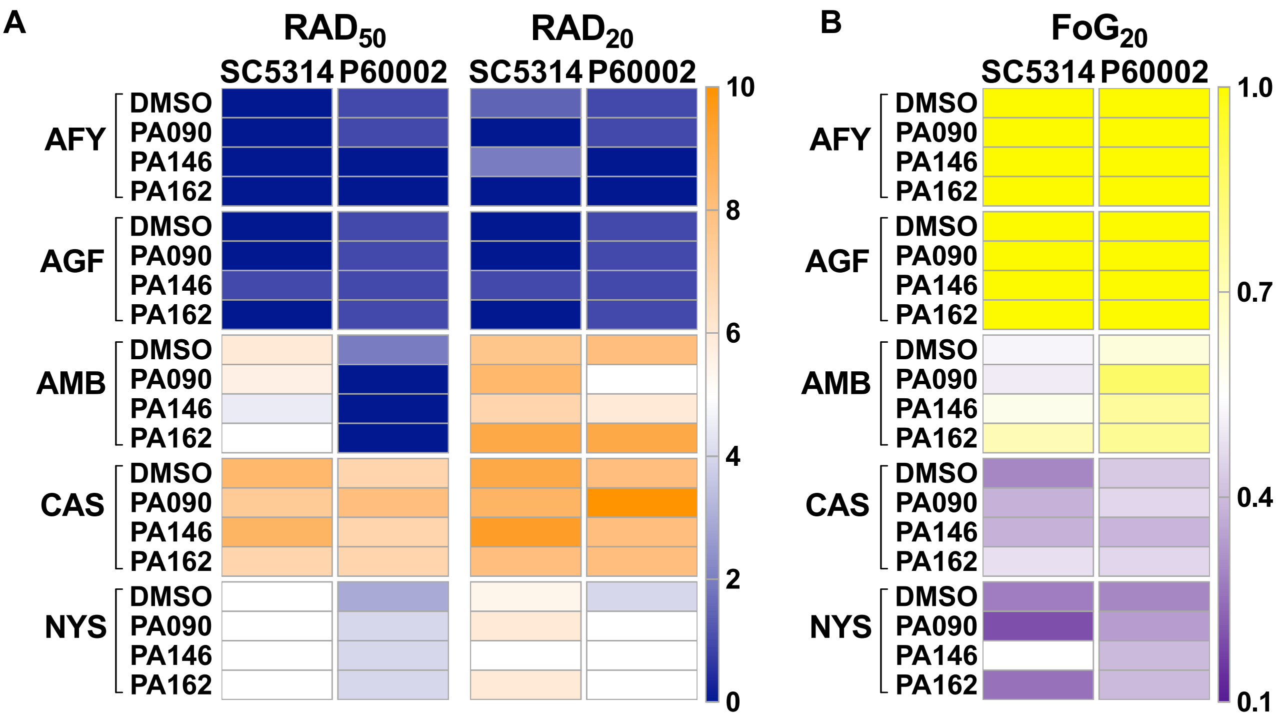

A
RAD50
RAD20
SC5314
P60002
SC5314
P60002
AFY
AGF
AMB
CAS
NYS
DMSO
PA090
PA146
PA162
10
8
6
4
2
0
B
FoG20
SC5314
P60002
1.0
0.7
0.4
0.1

### Figure S7.

**A**

| | | MIC$_{50}$ | SMG |
|---|---|---|---|
| FLC | SC5314 | 0.94 | 0.51 |
| | SP-945 | 8.00 | 0.78 |
| | P60002 | 128.00 | 0.87 |
| TERB | SC5314 | 4.00 | 0.54 |
| | SP-945 | 8.00 | 0.70 |
| | P60002 | 0.50 | 0.28 |
| FEN | SC5314 | 8.67 | 0.61 |
| | SP-945 | 32.00 | 0.85 |
| | P60002 | 7.33 | 0.56 |
| MYO | SC5314 | 16.00 | 0.54 |
| | SP-945 | 21.33 | 0.56 |
| | P60002 | 32.00 | 0.66 |
| FLC | ■ WT | 1.06 | 0.61 |
| | ▲ WT | 0.87 | 0.71 |
| | ■ *upc2* | 0.13 | 0.40 |
| | ▲ *erg5* | 0.40 | 0.87 |

**B**

| MIC$_{50}$ | SMG | |
|---|---|---|
| 1.06 | 0.61 | ■ WT |
| 1.00 | 0.47 | ▷ WT |
| 0.87 | 0.71 | ▲ WT |
| 1.00 | 0.68 | ◎ WT |
| 1.00 | 0.62 | ■ *mrr1* |
| 1.00 | 0.58 | ■ *tac1* |
| 1.00 | 0.87 | ▲ *adp1* |
| 1.00 | 0.95 | ▲ *mdl2* |
| 1.00 | 0.83 | ▲ *snq2* |
| 0.13 | 0.31 | ◎ *cdr1* |
| 0.13 | 0.22 | ▷ *cdr1* |
| 0.83 | 0.23 | ▷ *cdr2* |
| 0.08 | 0.24 | ▷ *flu1* |
| 1.00 | 0.50 | ▷ *mdr1* |
| 0.13 | 0.24 | ▷ *cdr1 cdr2* |
| 1.00 | 0.48 | ▷ *flu1 mdr1* |
| 0.50 | 0.22 | ▷ *cdr1 mdr1* |
| 1.00 | 0.30 | ▷ *cdr1 cdr2 flu1* |
| 0.06 | 0.24 | ▷ *cdr1 cdr2 mdr1* |
| 0.13 | 0.12 | ▷ *cdr1 cdr2 flu1 mdr1* |

**C**

| MIC$_{50}$ | SMG | |
|---|---|---|
| 1.06 | 0.61 | ■ WT |
| 6.67 | 0.74 | ■ *cph1* |
| 1.00 | 0.44 | ■ *efg1* |
| 1.00 | 0.44 | ■ *nrg1* |

**D**

| DMSO | |
|---|---|
| 17.8 | ▷ WT |
| 47.5 | ▷ *cdr1* |
| 30.8 | ▷ *cdr2* |
| 46.9 | ▷ *cdr1 cdr2* |

R6G fluorescence

0 25 50

**Figure S8.**

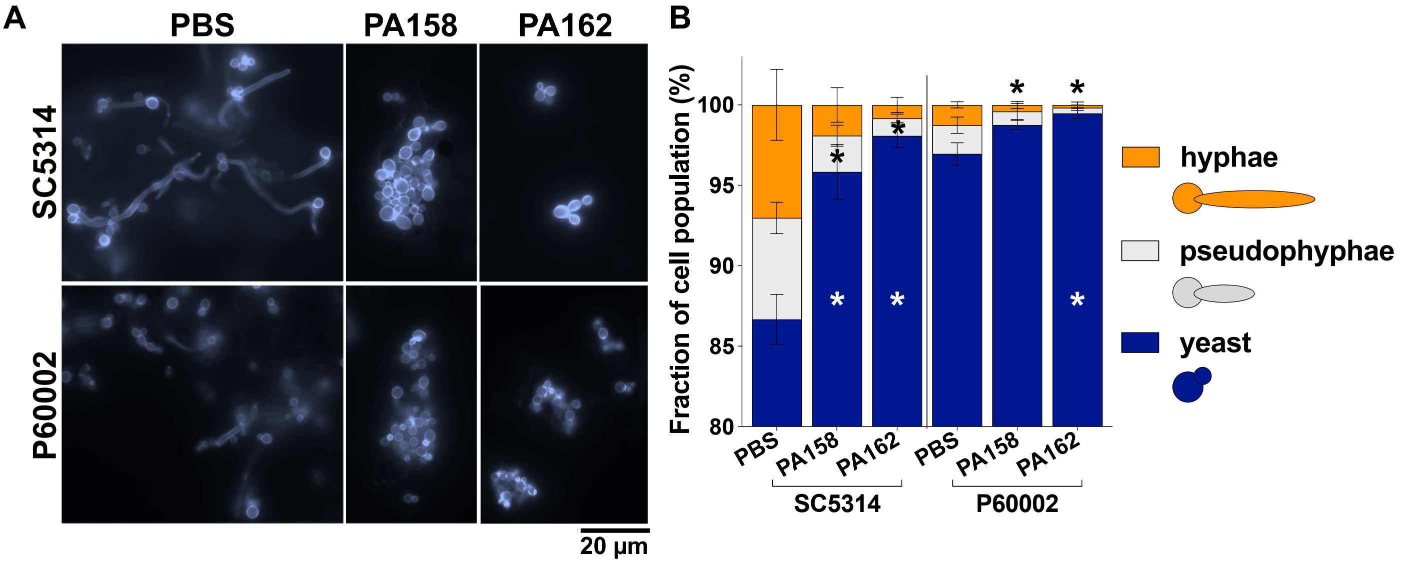

A
PBS
PA158
PA162
SC5314
P60002
20 µm
B
Fraction of cell population (%)
100
95
90
85
80
*
*
*
*
*
*
*
PBS
PA158
PA162
PBS
PA158
PA162
SC5314
P60002
hyphae
pseudophyphae
yeast
